# BysR, a LysR-type Pleiotropic Regulator, Controls Production of Occidiofungin by Activating the LuxR-type transcriptional regulator AmbR1 in *Burkholderia* sp. JP2-270

**DOI:** 10.1101/2022.07.06.499086

**Authors:** Lijuan Wu, Liqun Tang, Yuchang He, Cong Han, Lei Wang, Yunzeng Zhang, E Zhiguo

**Affiliations:** China National Rice Research Institute, Hangzhou, Zhejiang, People’s Republic of China; Joint International Research Laboratory of Agriculture and Agri-Product Safety, the Ministry of Education of China, Yangzhou University, Yangzhou, China

**Keywords:** Occidiofungin, LysR-type regulator, AmbR1, pleiotropic regulator, inhibitory activity

## Abstract

Occidiofungin is a highly effective antifungal glycopeptide produced by certain *Burkholderia* strains. The *ocf* gene cluster directing occidiofungin biosynthesis is regulated by cluster-specific regulators encoded by *ambR* homolog(s) within the same gene cluster, while it remains unknown to what extent occidiofungin biosynthesis is connected with the core regulation network. Here, we report that a LysR-type regulator BysR acts as a pleotropical regulator, and is essential for occidiofungin biosynthesis by directly activating *ambR1* transcription in *Burkholderia* sp. JP2-270. Deletion of *bysR* and *ocfE* both abolished the antifungal activity in JP2-270. This defect of the *bysR* mutant can be recovered by constitutively expressing *bysR* or *ambR1* but not by *ambR2*. The EMSA assay collectively showed that BysR regulates *ambR1* through direct binding to its promoter region. Taken together, occidiofungin produced by JP2-270 is the main substance inhibiting *M. oryzae*, and BysR controls occidiofungin production by directly activating the expression of *ambR1*. Besides, transcriptomic analysis revealed altered expression of 350 genes in response to *bysR* deletion, and the genes engaged in flagellar assembly and bacterial chemotaxis are the most enriched pathways. Also, 400 putative loci targeted by BysR were identified by DAP-seq in JP2-270. These loci not only include genes engaged in key metabolic pathways but also genes involved in secondary metabolic pathways. Collectively, we proposes that BysR may be a novel pleiotropic regulator, and *ambR1* is an important target gene of BysR, which is an intra-cluster transcriptional regulatory gene that further activates the transcription of *ocf* gene cluster.

**IMPORTANCE:** This study shows that BysR, a LysR-type transcriptional regulator (LTTR) from *Burkholderia* sp. JP2-270, activates the expression of *ambR1* gene responsible for regulating the synthesis of occidiofungin. BysR also acts as a pleiotropic regulator that controls primary and secondary metabolism, antibiotic resistance, motility, transport and other cellular processes in *Burkholderia* sp. JP2-270. This study provides insight into the regulatory mechanism of occidiofungin synthesis and enhances our understanding of the regulatory patterns of the LysR-type regulator.

## INTRODUCTION

The pathogenic fungi *Magnaporthe oryzae* is the causal agent of rice blast disease, which causes severe losses in grain yield annually. To manage blast disease, application of biocontrol strains of *Burkholderia* belonging to β-proteobacteria is a powerful strategy in addition to chemical fungicides and plant resistance breeding (1, 2). This is largely attributable to antifungal metabolites produced by certain *Burkholderia* isolates (2–4). The *Burkholderia cepacia* complex (Bcc) represents a diverse group of ubiquitously distributed bacteria in various niches, such as soil, water, plants, animals and humans (5). The Bcc has significant biotechnological potential as a source of novel antibiotics and bioactive secondary metabolites. Multiple bioactive secondary metabolites, such as pyrrolnitrin, occidiofungin, burkholdines, gladiolin, enacyloxin IIa and cepacins, which have shown great potential in biocontrol and pharmaceuticals, have been identified in members affiliated with Bcc (6–9).

Occidiofungin, a glycopeptide, was first identified from *Burkholderia contaminans* MS14 (10), which possesses significant antifungal and antiparasitic activities (2, 10, 11). Occidiofungin is biosynthesized by *ocfD, ocfE*, *ocfF*, *ocfH*, and *ocfJ*, which encode nonribosomal peptide synthetases (12, 13). Previous comparative genomics analysis suggested that the occidiofungin biosynthesis genes *ocfD*-*ocfJ* are recognized as a gene cluster and present in plant growth promoting *Burkholderia* strains but absent in pathogenic strains or non-plant-associated soil strains (14). The expression of *ocfD*-*ocfJ* is suggested to be specifically regulated by two LuxR-type regulatory genes, *ambR1* and *ambR2*, located in the same gene cluster with *ocfD*-*ocfJ*. AmbR1 plays a more critical role than AmbR2 (12, 13, 15). However, the integration mechanism of this secondary metabolism pathway with the core regulation network remains unknown.

In an elite biocontrol strain *Burkholderia* sp. JP2-270, we recently identified a transcriptional regulator BysR that is essential for the fungal inhibitory activity of JP2-270 that harbores a putative occidiofungin biosynthesis gene cluster (16). BysR belongs to the LysR-type transcriptional regulators (LTTRs). LTTRs are one of the most common transcription regulators in prokaryotes and can act as either positive or negative modulator for various genes involved in symbiotic nitrogen-fixation, virulence, quorum sensing, motility, and secondary metabolism (17–21). For example, the LTTR StgR inhibits the production of actinorhodin and prodigiosin by interacting with its pathway specific regulators in *Streptomyces coelicolor* (22), while the LTTR member FinR was recently reported to serve as an activator of phenazine and pyoluteorin biosynthesis in *Pseudomonas chlororaphis* G05 (21). Besides, ScmR was identified as a global LTTR controlling virulence and diverse secondary metabolites in *Burkholderia thailandensis*, like burkholdac, malleilactone, thailandamide, and acybolins (20).

This work aimed to investigate the regulon of the novel LTTR BysR, which is essential for the biocontrol activity of *Burkholderia* sp. JP2-270 against the pathogenic *M. oryzae*. By inactivating *ocfE*, we demonstrated that occidiofungin produced by JP2-270 mainly contributes the suppression activity towards *M. oryzae*. However, we found that BysR is required to inhibit the mycelial growth of *M. oryzae*. To understand the functions and regulatory mechanisms of BysR on regulation of occidiofungin production, as well as other pathways, we performed EMSA assays, RNA sequencing (RNA-seq) and DNA affinity purification sequencing (DAP-seq). The EMSA assays have showed that occidiofungin synthesis was regulated by BysR, and *ambR1*, the pathway specific regulatory gene of occidiofungin production, was the direct target of BysR. Aside from secondary metabolism, a series of regulatory targets, including genes engaged in core cellular processes, such as TCA cycle, DNA replication, amino acid synthesis, secretion (T6SS) system, efflux pump systems, outer membrane porins, transcriptional regulation (including H-NS histone family protein), transposase activity, quorum sensing system and TCSs, as well as c-di-GMP synthesis and secondary metabolites were identified in the chromosome of JP2-270 by DAP-seq. In general, our results identify a novel upstream component of the occidiofungin synthesis regulatory network, and provide new insight into the regulatory mechanisms of occidiofungin synthesis. The regulatory models have been proposed to help us understand the global regulatory role of LTTR transcriptional regulator BysR in diverse cellular processes.

## RESULTS

### BysR displays conserved domains of LTTR regulators

The *bysR* (DM992_17470) has an open reading frame of 984 bp, and encodes a putative 327 amino acid protein. The phylogenetic analysis of BysR with its most relevant LTTRs (protein sequence similarity >29%) revealed that these LTTRs were not conserved within a given genus, and the most homologous to BysR is BcaI3178 of *Burkholderia cenocepacia* H111, showing 93% identity (96% similarity) (Fig. 1A). LTTRs that have been reported to regulate secondary metabolite synthesis in *Burkholderia* include ScmR, ShvR and ToxR (20, 23, 24) (Fig. 1A). The domain homology search with Pfam (25) revealed that there is a typical HTH DNA binding domain (Pfam accession number PF00126) at the N-terminus and a typical C-terminus receiver domain (PF03466) (Fig. 1B). In addition, the homology comparison at the amino acid level between BysR and other identified LTTRs revealed that the high degree of similarity is within the N-terminal helix-turn-helix domain involved in binding DNA and the less conserved region was found in C-terminal, where the coinducer domain is located (Fig. 1C).

**Fig. 1.**
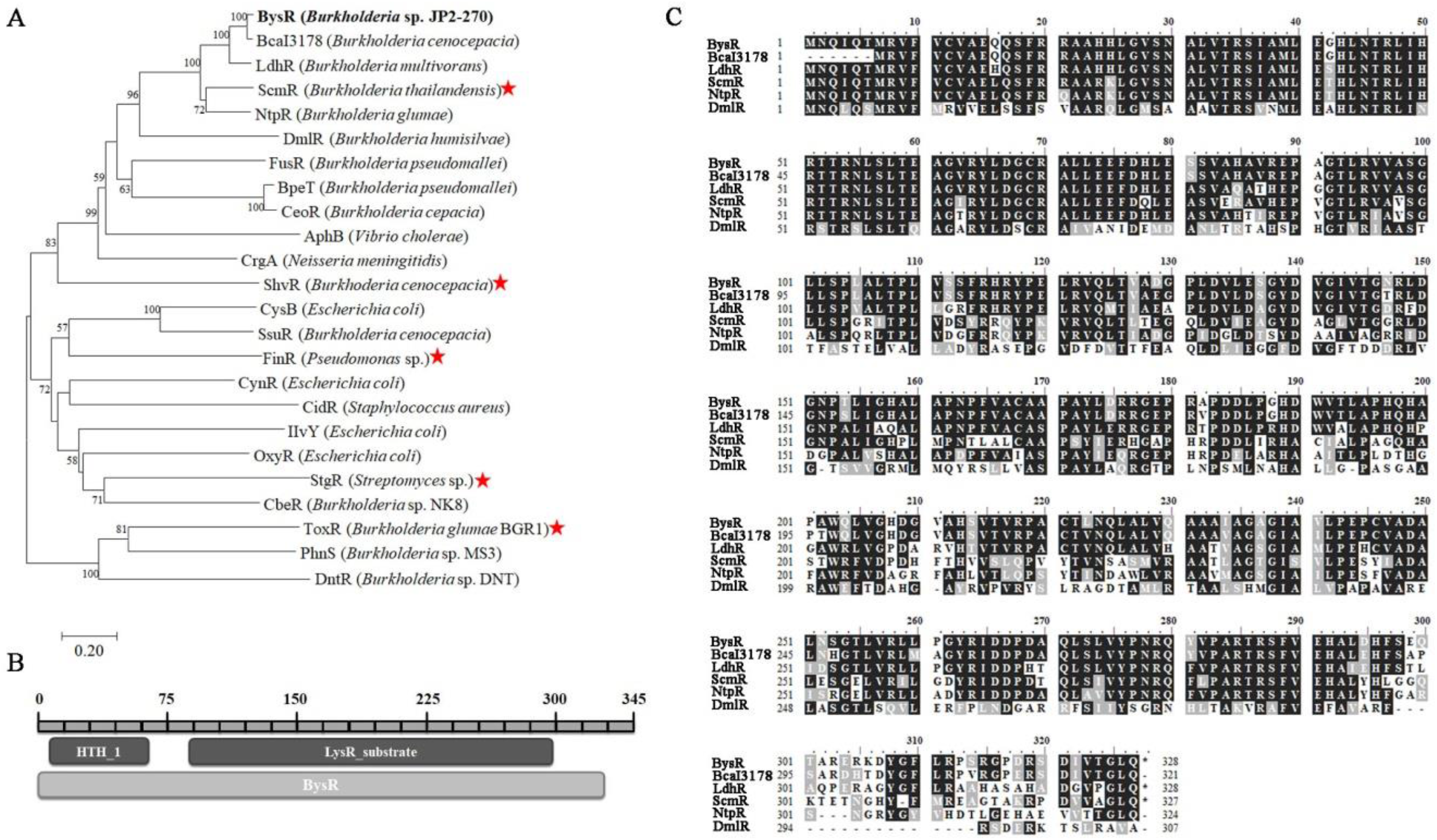
Structure and correlation of BysR and related LTTR proteins. A. Phylogenetic tree based on amino acid sequences of BysR and related LTTR proteins. The tree was generated with MEGA7 using the neighbor-joining method and a bootstrap value of 1,000 iterations. The LTTRs marked with red stars are involved in the regulation of secondary metabolites. B. Domain structure of BysR was analyzed by using the Pfam program at Pfam: Home page (xfam.org). C. Multiple sequence alignments for BysR. The alignments were conducted using MEGA7 (ClustalW algorithm). The consensus sequences are represented in black background. The similar sequences are shaded with grey background. The references protein BcaI3178 (I35_RS03450), LdhR (WP_012214022.1), ScmR (BTH_I1403), NtpR (WP_012734779.1) and DmlR (WP_175225518.1) were obtained from NCBI.

### BysR positively regulate multiple pathways in *Burkholderia* sp. JP2-270

Since BysR is a member of LTTRs, which are known as global regulators mediating gene expression of many pathways, we performed RNA-seq transcriptomic analysis to identify the potential regulated targets of BysR. Principal component analysis (PCA) revealed that three replicates of WT group and Δ*bysR* group are samples of high similarity, respectively, and the samples of different groups showed great dispersion (Fig. 2A). Besides, the high square (R^2^) of Pearson correlation coefficient between replicates of WT group (> 0.963) and Δ*bysR* group (> 0.944) also indicated that the data had a good reproducibility (Supplementary Fig. S1). The high consistence between the RNA-seq and qRT-PCR results further demonstrated the reliability of the RNA-seq results (Supplementary Fig. S2).

**Fig. 2.**
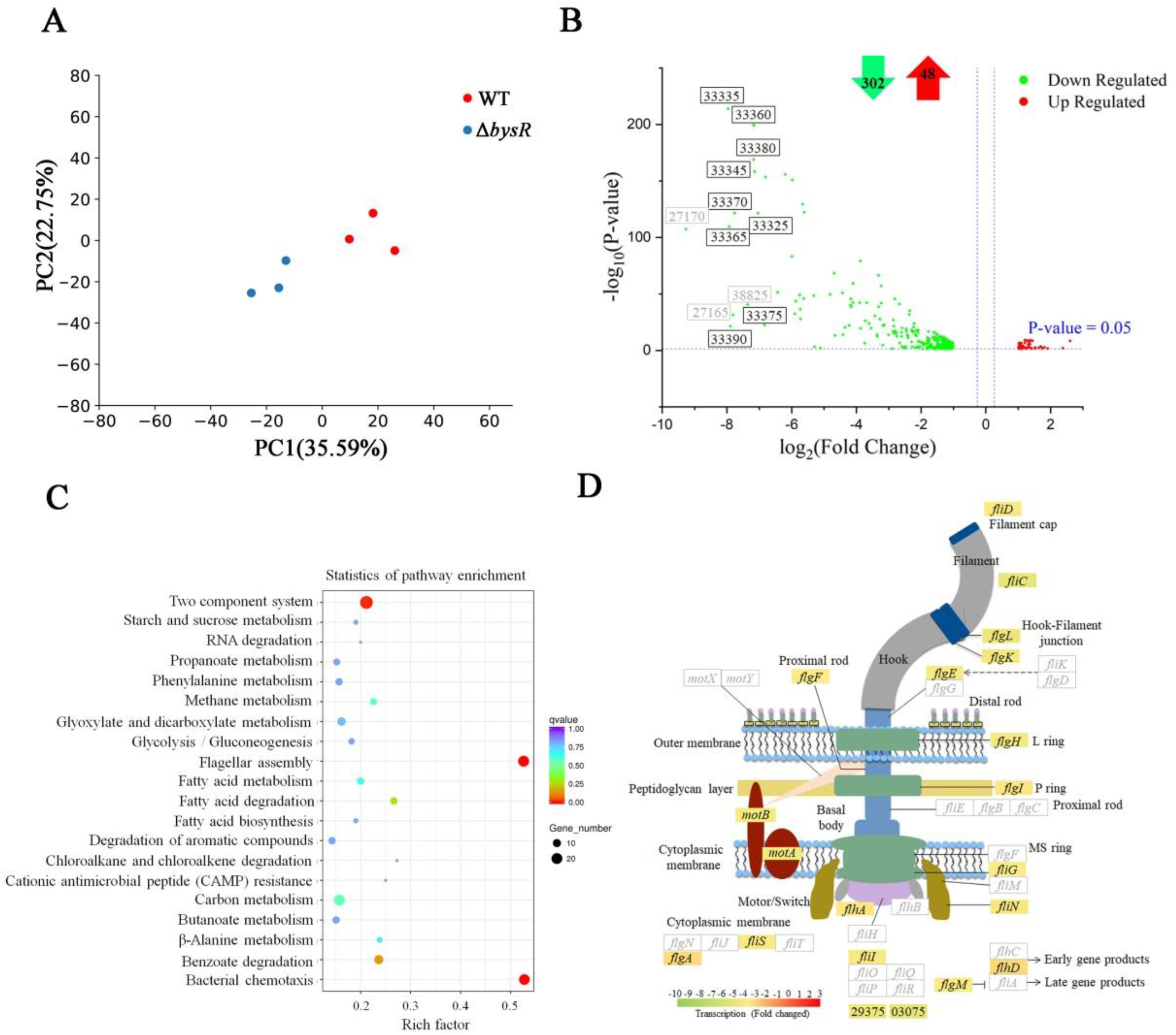
Identification of BysR-dependent genes in *Burkholderia* sp. JP2-270. (A) Principal component analysis (PCA) of six samples from RNA-seq analysis. The blue and red dots represent three different replicates of Δ*bysR* and WT treatment groups, respectively. (B) Volcano plot of RNA-seq data comparing the transcriptomes of Δ*bysR* and WT. Differentially expressed genes (|log2(FoldChange)| > 1, and FDR-adj *P* ≤ 0.01) are indicated in green dot (for down-regulated genes) and in red dot (for up-regulated genes). The genes with the most significant down-regulated in expression are named. The numeric string are the last 5 digits of the locus tag of the gene in NCBI and represent the gene names, eg. 33335 represent DM992_33335. The gene names in black indicate the genes involved in occidiofungin synthesis. Gene names in grey font indicate genes associated with other biological processes. The numbers of up- and down-regulated genes are showed at the top in red and green arrows, respectively. (C) The statistical enrichment of differential expression genes in KEGG pathways. The 20 most significant enrichment pathway items are shown. Rich factor represents the ratio of the number of differential expressed genes enriched in this pathway to the number of background genes. (D) The flagellar assembly model and the expression of related genes. Colored rectangles represent genes regulated by BysR based on the RNA-seq results, and the color scale indicates the relative transcriptional level (fold changed) based on RNA-seq results. The words in the rectangular boxes represent the names of the genes involved in flagellar assembly.

Differential gene expression analysis revealed 350 differentially expressed genes (DEGs) consisting of up-regulated (48) and down-regulated genes (302) (with p-adj < 0.05, |log2(FoldChange)| > 1) (Fig. 2B, Supplementary Table S3). The 12 genes with downregulated most significatly were marked, and 9 of these 12 genes related with the synthesis of occidiofungin (DM992_33325, DM992_33335, DM992_33345, DM992_33360-DM992_33380, DM992_33390) were dramatically downregulated by 3.6-7.9 folds in Δ*bysR* mutant (Fig. 2B and Fig. 3A). Among the down-regulated genes, besides the dramatically affected occidiofungin production-associated gene cluster, four other secondary metabolites biosynthetic gene clusters, including an unknown NRPS type gene clusters (DM992_33030 to DM992_33110, Region 3.1 in Table 1), an unknown bacteriocin type gene cluster (DM992_26140 to DM992_26160, Region 2.6 in Table 1), an *trans*-PKS gene cluster (DM992_38905 to DM992_38940, Region 3.7 in Table 1) involved in gladiostatin production and the pyrrolnitrin biosynthesis gene cluster (DM992_38820 to DM992_38835, Region 3.6 in Table 1) were also identified (Fig. 3A and Supplementary Table S3), implying that BysR regulates the production of secondary metabolites in *Burkholderia* sp. JP2-270. Among the up-regulated genes, the genes responsible for the biosynthesis of pyrroloquinoline quinone (PQQ) were affected significantly (Supplementary Table S3), suggesting that BysR may act as a negative regulator for the PQQ production.

**Fig. 3.**
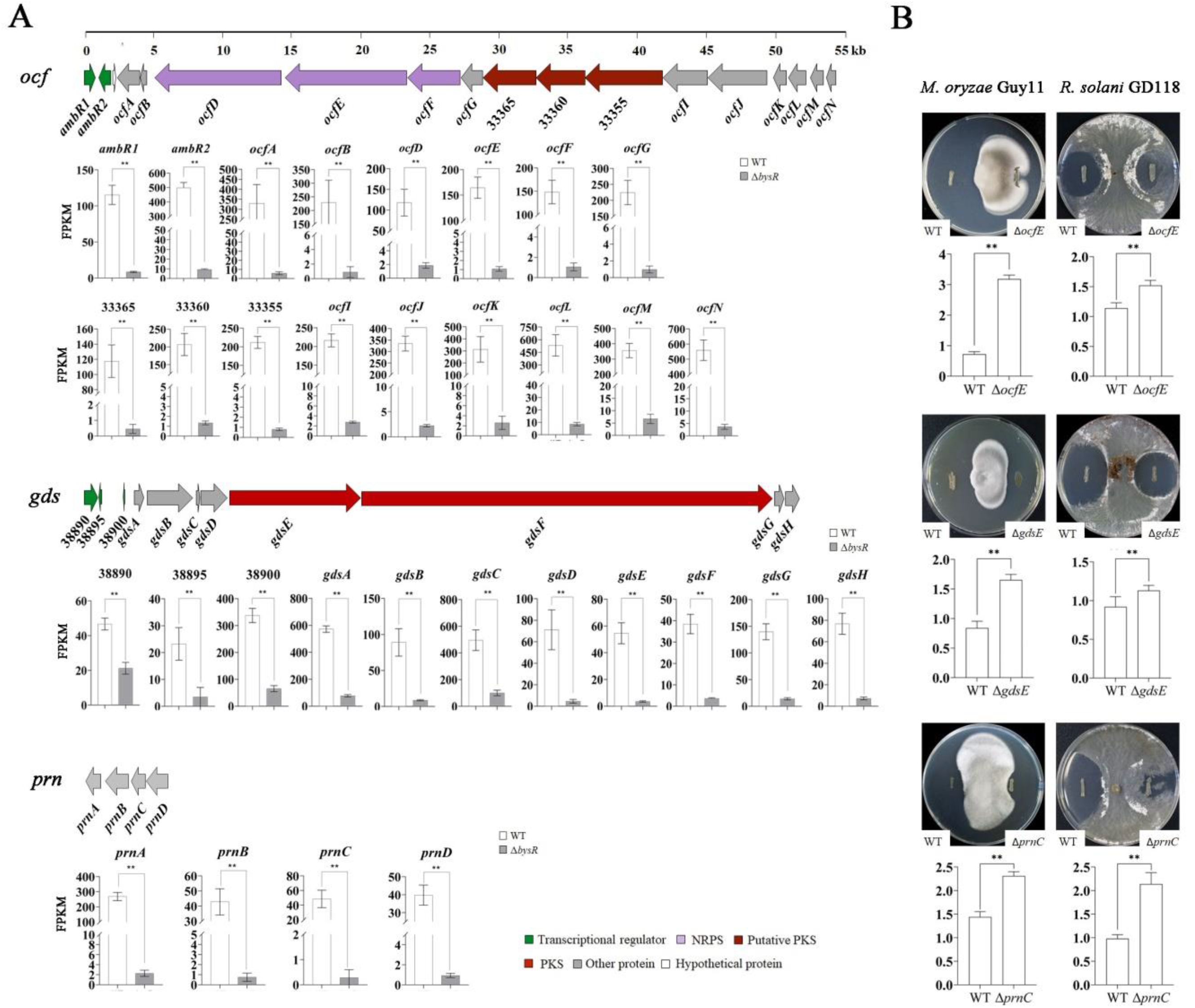
The gene clusters of secondary metabolite synthesis regulated by BysR. (A) Gene organization of occidiofungin (*ocf*), gladiostatin (*gds*) and pyrrolnitrin (*prn*) and the relative expression levels (FPKM) of genes in *ocf*, *gds* and *prn* gene clusters based on RNA-seq. (B) Antifungal activity analysis of key gene deletion mutants in *ocf*, *gds* and *prn* gene clusters. WT: wild type, Δ*ocfE*: the mutant with frame-deletion of *ocfE* encoding NRPS synthase, Δ*gdsE*: the mutant with frame-deletion of *gdsE* encoding PKS synthase, Δ*prnC*: the mutant with frame-deletion of *prnC* encoding FAD-dependent oxidoreductase.

**Table 1.** Gene clusters of *Burkholderia* sp. JP2-270 predicted by antiSMASH and BLAST analysis

| Cluster | Type | From-To | Most similar known cluster |  | Similarity |
| --- | --- | --- | --- | --- | --- |
| Chr1 (CP029824): |  |  |  |  |  |
| Region 1.1 | T1PKS | 1-32,032 | Capsular polysaccharide | Saccharide | 16% |
| Region 1.2 | Arylpolyene | 1,249,332-1,290,543 | APE Vf | Other | 10% |
| Region 1.3 | Terpene | 2,271,135-2,291,968 | - |  |  |
| Region 1.4 | NRPS | 2,482,384-2,529,203 | Burkholderic acid | NRP+Polyketide:Modular type I | 31% |
| Region 1.5 | NRPS | 2,723,377-2,778,049 | Ornibactin | NRP | 100% |
| Chr2 (CP029825): |  |  |  |  |  |
| Region 2.1 | Terpene | 275,900-296,964 | N-acyloxyacyl glutamine | Other | 50% |
| Region 2.2 | Phosphonate | 418,602-460,290 | Phosphinothricintripeptide | NRP | 6% |
| Region 2.3 | redox-cofactor | 1,186,903-1,209,041 |  |  |  |
| Region 2.4 | NRPS-like | 1,651,742-1,709,599 | Malleobactin A-D | NRP:NRP siderophore | 22% |
| Region 2.5 | Terpene | 1,784,391-1,808,491 | - |  |  |
| Region 2.6 | RRE-containing, RiPP-like | 1,834,070-1,847,121 | - |  |  |
| Region 2.7 | Phenazine | 1,874,156-1,894,584 | - |  |  |
| Region 2.8 | Hserlactone | 2,804,503-2,825,111 | - |  |  |
| Region 2.9 | Ectoine | 3,028,116-3,038,514 | - |  |  |
| Chr3 (CP029826): |  |  |  |  |  |
| Region 3.1 | TransAT-PKS,NRPS,NRPS-like | 1-47,903 | lagriamide | Polyketide | 9% |
| Region 3.2 | NRPS-like,T1PKS,NRPS | 77,377-163,367 | Occidiofungin A | NRP+Polyketide | 88% |
| Region 3.3 | Terpene | 448,407-470,461 | - |  |  |
| Region 3.4 | Hserlactone | 707,640-728,338 | - |  |  |
| Region 3.5 | RiPP-like | 1,069,004-1,079,819 | - |  |  |
| Region 3.6 | other | 1,328,932-1,360,585 | Pyrrolnitrin | Other | 100% |
| Region 3.7 | TransAT-PKS | 1,360,644-1,429,994 | Gladiostatin | Polyketide | 100% |
| Plas2 (CP029828): |  |  |  |  |  |
| Region 5.1 | Other | 63,674-106,502 | - |  |  |
| Region 5.2 | Other | 166,130-208,973 | O-antigen | Saccharide | 10% |

Moreover, KEGG pathway enrichment analysis of differentially expressed genes uncovered that a variety of cellular processes involved in carbon metabolism, chloroalkane and chloroalkene degradation, two-component system, bacterial chemotaxis and flagellar assembly were significantly enriched (Fig. 2C and Supplementary Table S4). The genes associated with these enriched pathways were all down-regulated in Δ*bysR* (Supplementary Table S3 and S4). Notably, the proteins involved in flagellar assembly were the most enriched pathways, and 20 of the 38 genes engaged in flagellar assembly were found to be down-regulated in Δ*bysR* (Fig. 2D), indicating that BysR plasys an important role in regulating flagellar assembly.

Collectively, the RNA-seq analysis shows that BysR controls expression of various genes, and the gene encoding proteins engaged in flagellar assembly and chemotaxis are most enriched pathways in diverse cellular processes. Also, the genes associated with secondary metabolites, such as occidiofungin, pyrrolnitrin and gladiostatin, were most significantly affected by BysR. Besides, *bysR* also affects the genes related to co-factor protein synthesis, various transcriptional regulators, transporters, energy metabolism, stress response, electron transfer, nucleotide metabolism, glycometabolism, amino acid metabolism, heavy metal translocating, and other genes, whose importance is not yet known. The number of down-regulated genes (302) was greater than that of up-regulated genes (48) in Δ*bysR* (relative to WT), indicating that this LTTRs-BysR mainly acts as a positive regulator in *Burkholderia* sp. JP2-270.

### BysR may function as a pleiotropic modulator directly regulate gene expression in various cellular processes

The DEGs identified by RNA-seq include those that are regulated directly and indirectly by BysR. To verify genes that are directly regulated by BysR, we used DNA affinity purification sequencing analysis (DAP-seq) for genome-wide recognition of BysR binding sites *in vitro* (26). After affinity purification and sequencing, at least 22 million double-end reads per sample were generated and with > 99% of reads were uniquely mapped to the JP2-270 genome. A total of 400 enriched common peaks of two replicates with –log10(p-value) ≥ 2 were called (Supplementary Table S5). The mean width of DAP-seq peaks was < 1,000 bp as shown in Fig 4A. In total, 89.6% BysR binding peaks were distributed in the promoter, promoter-TSS or intergenic regions, while only 8.3% and 2.3% were in the TTS and exon regions, respectively (Fig. 4B). Among the identified peaks, 372 (93%) of these peaks were found to locate in the −1 kbp to 1 kbp regions by the analysis of peak summit positions relative to the start codons of JP2-270 open reading frames (or the first genes in operons) (Fig. 4C, Supplementary Table S5). These results suggest that the expression of these loci might be directly regulated by BysR. A motif of T-N_11_-A (LTTR box) (Fig. 4D) was found as considerably enriched (e value = 9.8e-010) among all the peaks using MEME-ChIP (27), indicating that the nucleotide sequences of the motif shared by BysR binding regions are similar with binding sites of other LTTRs (17, 28). It is remarkably that *bysR* itself promoter loci (peak ID: Merged-Chr1-3684201-2 in Supplementary Table S5, Fig. 4E) was identified among the peaks, providing a positive control to test the reliability of this approach and verifying the autoregulatory properties of BysR. In addition, the identification of the *ambR1* promoter loci (peak ID: Merged-Chr3-148813-2 in Supplementary Table S5, Fig. 4E) indicated that BysR may regulate the expression of *ocf* gene cluster by directly binding to the *ambR1* promoter region.

**Fig. 4.**
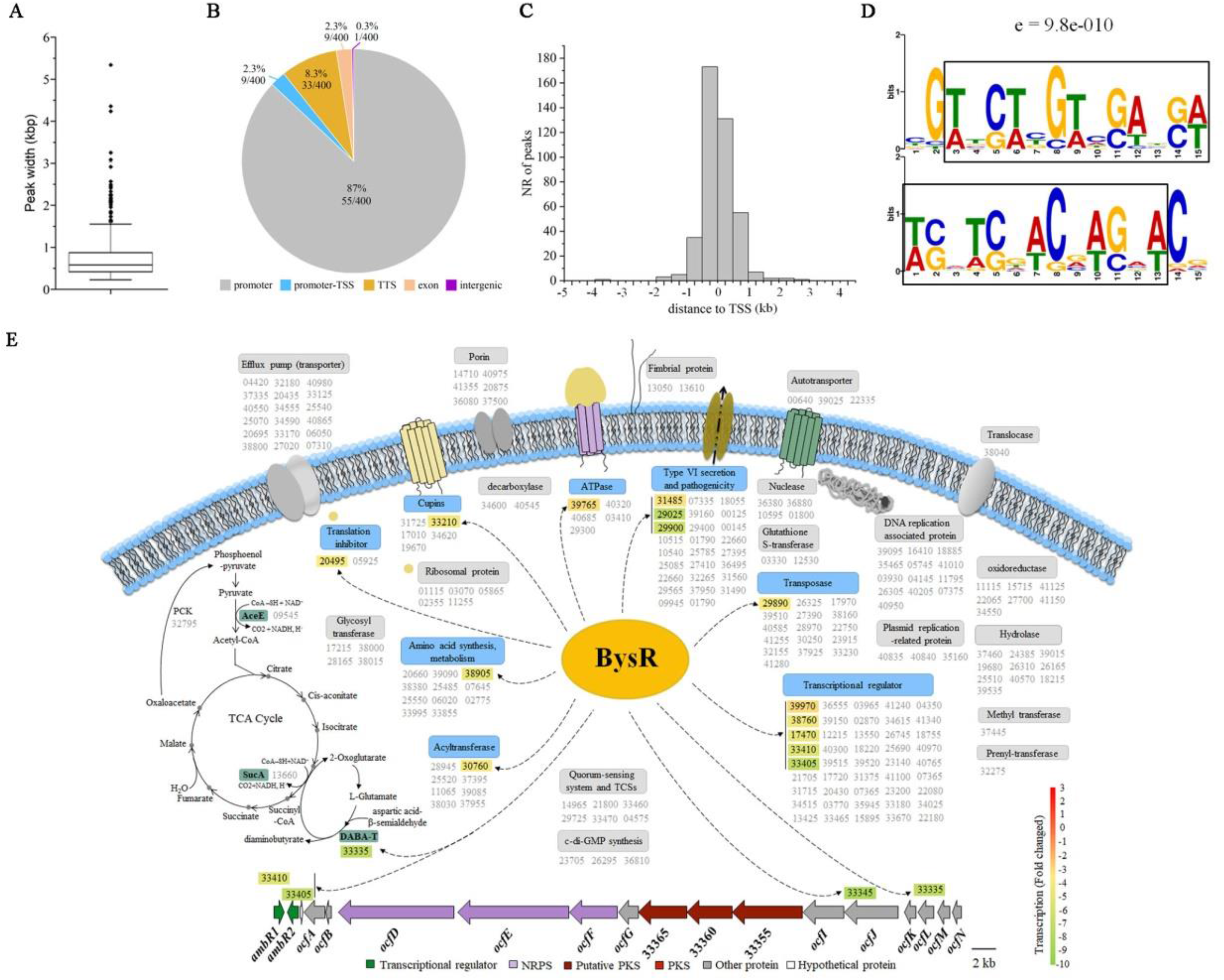
Identification of genes directly regulated by BysR in *Burkholderia* sp. JP2-270. (A) The width of BysR DAP-seq peaks. (B) The number and distribution of BysR binding genes obtained by DAP-seq. (C) Distribution of the distance between DAP-seq peak summit positions relative to the start codons of JP2-270 open reading frames. (D) The predicted binding motif of BysR by MEME-ChIP (e-value = 9.8e-010). The reverse complement of this motif logo are presented below. The putative LTTR box (T-N_11_-A) is framed in black. (E) The proposed model for the regulation of various cellular processes by BysR in *Burkholderia* sp. JP2-270. The black dotted arrows indicates directly activation by BysR. The numeric string are the last 5 digits of the locus tag of the gene in NCBI and represent the gene names, eg. 33335 represents DM992_33335. Gene names in grey font indicate genes putative directly regulated by BysR based on only DAP-seq results. Gene names in black font on a colored background represent genes putative directly regulated by BysR based on both the RNA-seq and DAP-seq results. The color scale indicates the relative transcriptional level (log_2_-fold change) of the genes in Δ*bysR* relative to WT based on RNA-seq results.

Specifically, our data suggested that the genes encoding pyruvate dehydrogenase (DM992_09545), 2-oxoglutarate dehydrogenase (DM992_13660), phosphoenolpyruvate carboxylase (DM992_32795) and diaminobutyrate-2-oxoglutarate transaminase (DM992_33335) envolved in pyruvate metabolism and TCA cycle were the targets directly activated by BysR (Fig. 4E and Supplementary Table S5). In addition, the transcriptional level of gene enconding diaminobutyrate-2-oxoglutarate transaminase (DM992_33335) responsible for the synthesis of 2-oxoglutarate and 2-aminobutyric acid was found significantly down-regulated in Δ*bysR* (Fig. 4E and Supplementary Table S3). Moreover, direct target genes regulated by BysR are also involved in other core metabolism, such as amino acid synthesis and metabolism, and DNA replication (Fig. 4E and Supplementary Table S5). Besides, some genes related to drug resistance, transport, mobility, transposase activity, transcriptional regulation and quorum sensing system were also identified as the direct targets of BysR (Fig. 4E and Supplementary Table S5), and the genes associated with type VI secretion system and transcriptional regulation are account for the largest proportion. It is found that transcriptional regulators-related genes were the largest number of gene group directly regulated by BysR, including DM992_33410 (*ambR1*)/DM992_33405 (*ambR2*) and DM992_38895, which are transcriptional regulators located in the same cluster of occidiofungin (Region 3.2) and gladiostatin (Region 3.7) biosynthesis gene clusters, respectively (Fig. 4E and Table 1).

Overall, BysR may be a pleiotropic transcriptional factor that directly and indirectly regulates gene expression in a variety of cellular processes, like primary and secondary metabolism, antibiotic resistance, motility, transport and others in *Burkholderia* sp. JP2-270.

### *Burkholderia* sp. JP2-270 shows excellent inhibitory activity against *M. oryzae*

In our previously research, we have demonstrated that JP2-270 could suppress the growth of *Rhizoctonia solani* GD118 (16). Furthermore, in this study we also assess the inhibitory activity of JP2-270 against *M. oryzae* Guy11. As shown in Fig. 5A and 5I, the mycelial growth of Guy11 were obviously inhibited by JP2-270, relative to the control, with the mycelial growth radius of Guy11 being 0.72 ± 0.08 cm and 3.44 ± 0.10 cm after seven days of incubation with or without JP2-270, respectively (p < 0.01) (Figs. 5A, 5B and 5I). Our results indicated that *Burkholderia* sp. JP2-270 displayed significant suppressive activity towards *M. oryzae*.

**Fig. 5.**
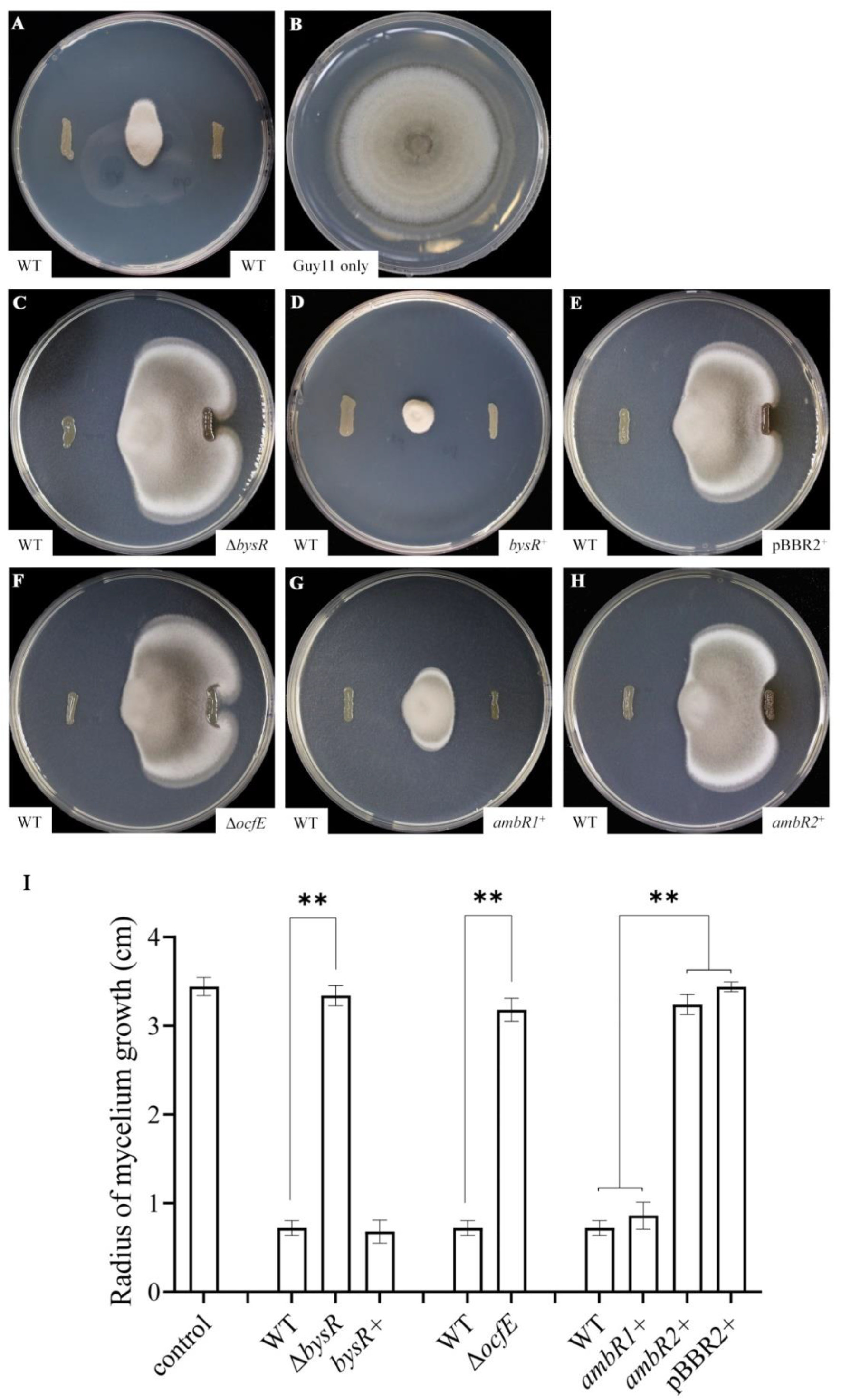
Inhibitory activity of *Burkholderia* sp. JP2-270 and its derivates against *M. oryza*. A, The mycelial growth of *M. oryzae* was inhibited by wild type isolate JP2-270 (WT). B, Uninhibited mycelial growth of *M. oryzae* was used as control. C, The mycelial growth of *M. oryzae* was inhibited by isolates of wild type JP2-270 (WT) and Δ*bysR*. D, The mycelial growth of *M. oryzae* was inhibited by isolates of wild type JP2-270 (WT) and Δ*bysR^+^*. E, The mycelial growth of *M. oryzae* was inhibited by isolates of wild type JP2-270 (WT) and pBBR2*^+^*. F, The mycelial growth of *M. oryzae* was inhibited by isolates of wild type JP2-270 (WT) and Δ*ocfE*. G, The mycelial growth of *M. oryzae* was inhibited by isolates of wild type JP2-270 (WT) and *ambR1^+^*. H, The mycelial growth of *M. oryzae* was inhibited by isolates of wild type JP2-270 (WT) and *ambR2^+^.* Δ*bysR^+^*: Δ*bysR+*pBBR2*-bysR*, pBBR2*^+^*: Δ*bysR+*pBBR2, *ambR1^+^*: Δ*bysR+*pBBR2*-ambR1*, *ambR2^+^*: Δ*bysR+*pBBR2*-ambR2*. Data are expressed as mean ± SD of quintuplicate, ** means extremely significant difference compared with WT (*P* <0.01).

### BysR deficiency reduced inhibitory activity against *M. oryzae*

As a pleiotropic transcriptional factor, BysR could regulate various cellular processes, including but not limited to secondary metabolism. In our previously research, we have demonstrated that BysR played an important role for the suppressive activity of *Burkholderia* sp. JP2-270 against *R. solani* (16). Here, we further demonstrated that while the radius of mycelial growth was significantly inhibited by wild-type JP2-270, the Δ*bysR* mutant almost totally lost the inhibitory activity against *M. oryzae* (Fig. 5C and 5I). The reduced suppressive activity of Δ*bysR* could be restored by introducing pBBR2-*bysR* expressing wild-type *bysR* but not by the empty vector pBBR1MCS-2 (Figs. 5D, 5E and 5I). The strong antagonistic activity against *M. oryzae* and *R. solani* GD118, indicative of wide-spectrum antifungal activity of JP2-270.

### Occidiofungin is the main secondary metabolite with antifungal activites

As mentioned above, we found that several genes involved in occidiofungin (Region 3.2), gladiostatin (Region 3.7) and pyrrolnitrin (Region 3.6) biosynthesis were dramatically downregulated by 3.6-7.9 folds, 1.1-3.9 folds and 5.3-7.4 folds in Δ*bysR* mutant, respectively (Fig. 3A, Supplementary Table S3). To understand the role of these secondary metabolites in inhibiting mycelial growth of fungi, we constructed the marker-less deletion mutants of *ocfE*, *gdsE* and *prnC*, and were tested for inhibitory activities. The *ocfE*, *gdsE* and *prnC* encode the non-ribosomal peptide synthetase, polyketide synthase and FAD-dependent oxidoreductase, respectively, which are key enzymes in the synthesis of occidiofungin, gladiostatin and pyrrolnitrin, respectively. Based on the inhibition assays, we observed that the mycelium growth radius of Guy11 was 3.18 ± 0.13 cm for Δ*ocfE* (relative to 0.72 ± 0.08 cm for JP2-270), while the radius were 1.65 ± 0.09 cm for Δ*gdsE* (relative to 0.84 ± 0.11 cm for JP2-270) and 1.44 ± 0.11 cm for Δ*prnC* (relative to 2.31 ± 0.09 cm for JP2-270) (Fig. 3B). The loss of *ocfE* caused significantly reduced suppressive activity towards *M. oryzae* compared with wild-type JP2-270, indicating that occidiofungin produced by JP2-270 plays more important role than gladiostatin and pyrrolnitrin in inhibiting mycelial growth of *M. oryzae*. While, pyrrolnitrin produced by JP2-270 is the main metabolite inhibiting *R. solani*, with mycelium growth radius were 0.98 ± 0.08 cm, 2.14 ± 0.24 cm for Δ*prnC* and JP2-270, respectively. While occidiofungin and gladiostatin showed relatively lower activity against *R. solani*, with the radius are 1.14 ± 0.09 cm for Δ*ocfE* (relative to 1.52 ± 0.08 cm for JP2-270), 0.92 ± 0.13 cm for Δ*gdsE* (relative to 1.13 ± 0.07 cm for JP2-270). Overall, occidiofungin and pyrrolnitrin had better inhibitory activity against *M. oryzae* and *R. solani*, respectively, while gladiostatin showed moderate inhibitory activities against the tested fungi (Fig. 3B). Collectively, theses results showed that JP2-270 could produce a variety of secondary metabollites, among which occidiofungin is critical for inhibiting *M. oryzae*.

### Comparative evolutionary analysis based on genome and Ocf proteins

Phylograms inferred from genome sequences revealed that the closest species of JP2-270 is *Burkholderia pyrrocinia*, and both of them belonged to the Bcc group (Fig. 6A). Occidiofungin was originally isolated from strain *Burkholderia contaminas* MS14, and the *ocf* gene cluster has also been found in 9 species of *Burkholderia*, including *Burkholderia cepacia*, *Burkholderia ubonensis*, *Burkholderia stabilis*, *Burkholderia anthina*, *Burkholderia contaminans*, *Burkholderia vietnamiensis*, *Burkholderia ambifaria*, *Burkholderia pyrrocinia* and *Burkholderia* sp. JP2-270 (Fig. 6B). The phylogenetic tree based on the concatenated sequences of AmbR1/2 and OcfA-OcfN proteins was constructed (Fig. 6B), and the inconsistent evolutionary trajectory of Ocf proteins and *Burkholderia* genome was founded by contrasting the corresponding phylogenetic trees (Fig. 6A and 6B). This result indicated that occidiofungin genes may acquired via horizontal gene transfer.

**Fig. 6.**
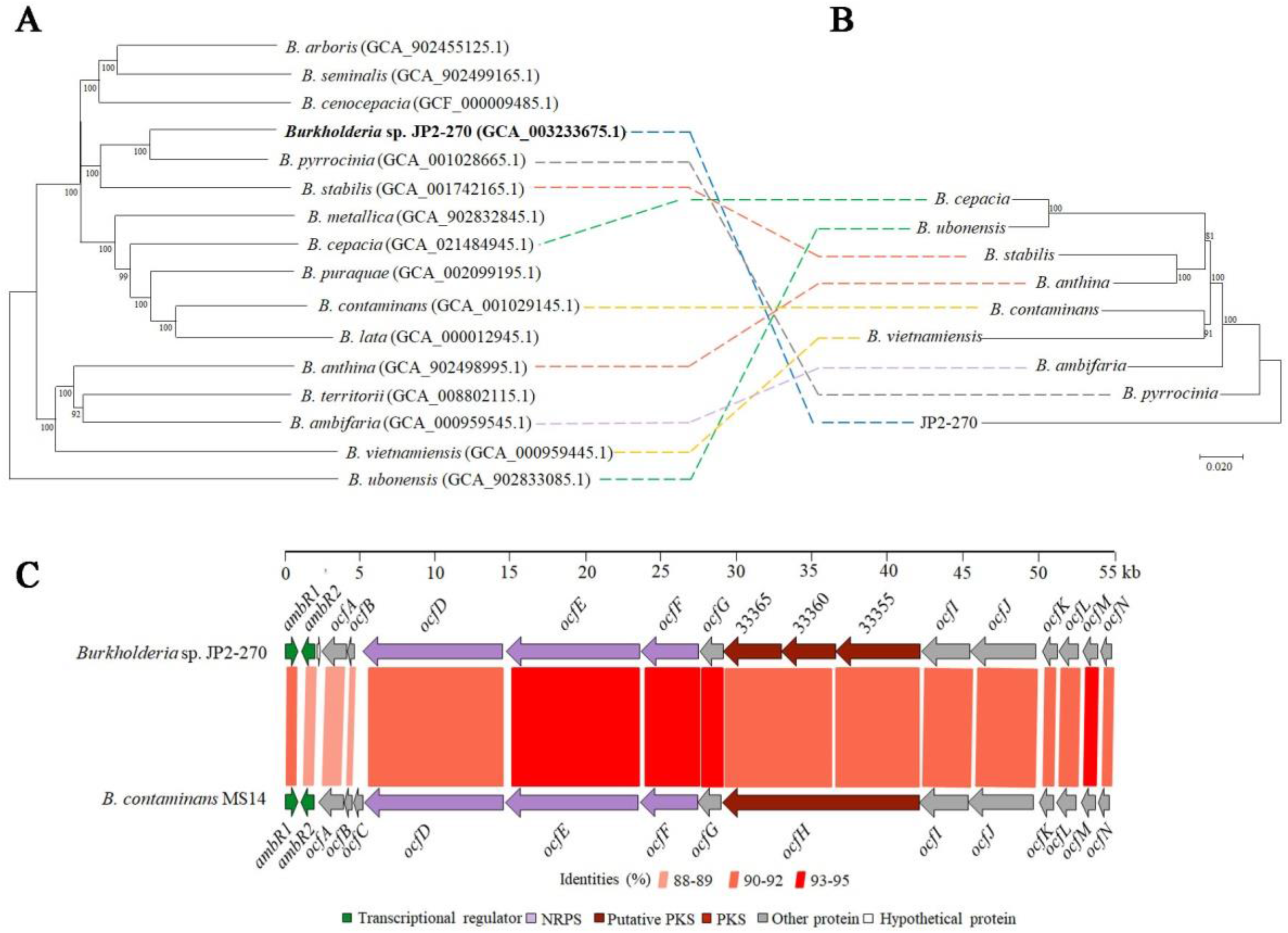
Comparative evolutionary analysis based on genome and Ocf proteins. (A) Phylogentic tree inferred with FastME 2.1.6.1 [7] from GBDP distances calculated based on genome sequences of representative species belonging to *Burkholderia*. The branch lengths are scaled in terms of GBDP distance formula *d*_5_. The numbers above branches are GBDP pseudo-bootstrap support values > 60 % from 100 replications, with an average branch support of 98.7 %. (B) The phylogenetic tree of Ocf protein generated by MEGA7. Ocf protein was found only in these 8 known species of *Burkholderia* and JP2-270. The amino acid substitution model was poisson model. The evolutionary history was inferred using the neighbor-joining method. Bootstrap values (> 60%) from 1000 replicates are showed at the nodes. The tree was drawn to scale, with branch lengths in the same units as those of the evolutionary distances used to infer the phylogenetic tree. (C) Comparison of the gene organization for the occidiofungin gene clusters of *Burkholderia* sp. JP2-270 and *B. contaminans* MS14. The predicted occidiofungin biosynthesis genes are highly conserved between JP2-270 and MS14.

In detail, we compared the *ocf* genes between *Burkholderia* sp. JP2-270 and *B. contaminans* MS14, and found that *ocfC* enconding glycosyl transferase was absent in the *ocf* gene cluster of JP2-270 (Fig. 6C). Besides, the *ocfH* in MS14 was highly homologous with three independent genes DM992_33365, DM992_33360 and DM992_33355 in JP2-270 (Fig. 6C). Overall, the genes in the *ocf* gene cluster of JP2-270 have highly similarity with the corresponding genes in MS14 (Fig. 6C).

### AmbR1 is a downstream component of the BysR regulation system

It is known that *ambR1* is a pathway specific regulatory gene, which positively regulates the biosynthesis of occidiofungin. Based on the fact that Δ*ocfE* displays significantly decreased inhibitory activity against *M. oryzae* compared to JP2-270 and the expression of *ambR1* is significantly down-regulated in Δ*bysR* as shown both in RNA-seq and qRT-PCR results, we infered that BysR might regulate the synthesis of occidiofungin by modulating *ambR1* expression. Thus, we enhance the expression of *ambR1* in the *bysR* mutant strain by introducing *ambR1* expression vector into Δ*bysR*, and the result showed that the overexpression of *ambR1* in Δ*bysR* increases suppressive activity towards *M. oryzae* (Fig. 5G and 5I). The observed radius of mycelial growth were 0.86 ± 0.15 cm for *ambR1^+^* (Δ*bysR+*pBBR*-ambR1*) and 0.72 ± 0.08 cm for JP2-270, indicating that *ambR1* could almost fully restore the inhibitory activity against *M. oryzae* to the wild-type level (Fig. 5G and 5I). However, the *ambR2* could not recover the inhibitory activity of Δ*bysR* against *M. oryzae* (Fig. 5H and 5I). In conclusion, these results suggested that *ambR1* is a downstream component of the BysR regulation system in JP2-270.

### BysR regulates *ambR1* expression by directly binding to the promoter

To assess whether transcriptional regulation of *ambR1* was performed by directly binding of BysR to the promoter, we performed electrophoretic mobility shift analyses (EMSAs). BysR fused with a N-terminal GST tag was purified using affinity chromatography, and then about 39 kDa BysR protein was obtained by excising GST tag using PreScission protease (Fig. 7A). Three DNA fragments of the *ambR1* promoter 5’-labled with Cy5 were obtained by PCR amplification for use as the probes. Probe 1 consists of fragment from positions −677 to −844 relative to the translation initiation site of *ambR1*, abbreviated as probe 1 (−677 to −844) (Fig. 7B). Similarity, the other two probes were named as probe 2 (−7 to −336) and probe 3 (−160 to −692) (Fig. 7B). The EMSAs showed that the complex of BysR and probe 3 migrated slower than the unbound probe, while there were no shift bands observed for probe 1 and probe 2 (Fig. 7B). Moreover, the probe 3 that bound to BysR significantly increased with increasing amounts of BysR protein (Fig. 7C). These results indicated that BysR could bind to the promoter region of *ambR1*, and the promoter region from −160 to −692 is essential for binding with BysR.

**Fig. 7.**
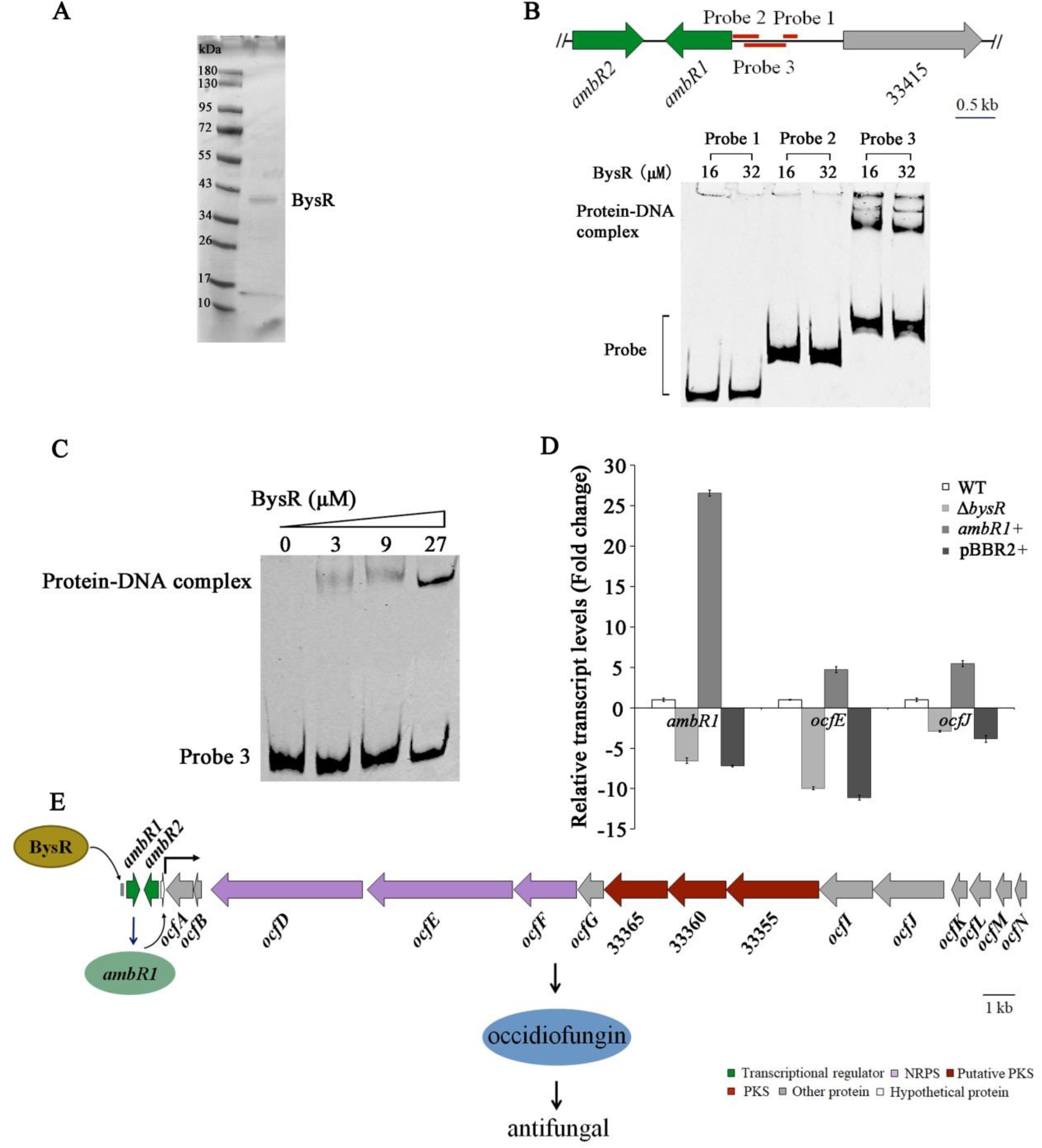
The regulatory mode of BysR on occidiofungin synthesis. (A) SDS-PAGE of purified BysR protein. (B) The relative positions of different probe fragments in *ambR1* promoter region and the binding of the probes to BysR protein by EMSA assays. (C) EMSA assays of BysR binding to the probe 3 of *ambR1* promoter region with increasing concentrations of BysR. (D) The expression level of *ambR1*, *ocfE* and *ocfJ* in isolates of wild-type JP2-270 (WT), Δ*bysR*, Δ*bysR*+pBBR2-*ambR1* andΔ*bysR*+pBBR2. The *recA* gene is used as a control. The relative gene expression levels are presented as expression ratios of the indicated genes. (E) The proposed model of BysR controls the production of occidiofungin by directly activating the transcription of *ambR1*.

### BysR regulates occidiofungin biosynthesis at the transcriptional level

To clarify the regulatory effect of BysR on the *ocf* operon, we applied qRT-PCR analysis to assess the transcriptional changes of *ocfJ*, *ocfE* and *ambR1* in wild-type JP2-270, Δ*bysR, ambR1^+^*(Δ*bysR+*pBBR2*-ambR1*) and pBBR2*^+^*(Δ*bysR+*pBBR2). As shown in Fig. 7D, the deficiency of *bysR* resulted in 7.2-fold to 28.6-fold reduction in the transcriptional level of these three selected genes, relative to the wild-type JP2-270. Moreover, overexpression of *ambR1* in Δ*bysR* could increase the expression level of *ocfJ* and *ocfE*, resulted in 5.45-fold and 4.79-fold up-regulations of *ocfJ* and *ocfE*, relative to WT, respectively (Fig. 7D). While, the expression level of *ocfJ*, *ocfE* and *ambR1* in pBBR2*^+^* are similar to that of wild type JP2-270. The 26.56-fold up-regulated of *ambR1* transcriptional level was observed in Δ*bysR* harboring pBBR2-*ambR1* (Fig. 7D), suggesting that *ambR1* had been successfully overexpressed in Δ*bysR*.

Based on our results, we could concluded that BysR binds directly to the promoter region of *ambR1*, and activates the transcription of *ambR1*, then AmbR1 promotes transcription of *ocf* genes located in the downstream of *ambR1* and thus controls the production of occidiofungin (Fig. 7E).

## DISCUSSION

Occidiofungin is a natural product produced by *Burkholderia* spp., and possesses a broad-spectrum antifungal, antiparasitic and anticancer activity with limited toxicity and chemical stability (2, 11, 29–31). Besides, its great application potential in agricultural practices for controlling fungi-causing plant diseases, occidiofungin also has been developing as a lead compound for clinical antifungal therapeutics (e.g., candidiasis) and antiparasitic treatment owing to the distinctive mode of action (2, 11, 30, 31). In addition, occidiofungin and its derivatives could also be developed as anticancer drugs due to the targeting effect on some cancer cell lines (29). However, the lack of knowledge on the regulatory mechanisms of occidiofungin production significantly hinders us from effectively utilization of the occidiofungin-producing strains, such as identification of highly productive strains and manipulation of the occidiofungin production pathway. In this study, using our identified beneficial strain *Burkholderia* sp. JP2-270 as a model, we demonstrated that the occidiofungin production pathway is regulated by a novel global LTTR, BysR, by directly regulating the transcription of *ambR1* that is known as the direct regulator of occidiofungin biosynthesis located in the *ocf* gene cluster. Previous studies suggested *ambR1* and *ambR2* were the only two genes in charge of *ocfD*-*ocfJ* expression regulation (12, 13, 15, 32). Our results demonstrated that *ambR1* but not *ambR2* is a direct downstream regulatory gene of BysR, and *bysR* may directly regulate the expression of occidiofungin synthesis such as *ocfJ* as revealed by the DAP-seq analysis. These results suggested more complex and well-tuned regulation maps of occidiofungin synthesis in *Burkholderia*.

BysR is identified as a member in the LysR-type transcription regulators (LTTRs) family. As global regulators, LTTRs are the most widespread in prokaryotes and can regulate the expression of genes involved various cellular processes in different microorganisms, such as the control of carbon catabolism, amino acid metabolism, antibiotic resistance and motility (11, 33–37). Here, we explored the RNA-seq and DAP-seq data, and the genes and associated pathways directly or indirectly regulated by BysR in JP2-270 were identified.

RNA-seq results showed that the expression of genes associated with various cellular processes were positively regulated by BysR, of which the genes related to flagellar assembly and bacterial chemotaxis are the most enriched pathways in KEGG analysis (Fig. 2C). However, all of DEGs related to the flagellar assembly and bacterial chemotaxis identified in RNA-seq are absent in DAP-seq results, indicating that these genes may be indirectly regulated by BysR. The Lrp family transcriptional regulator and H-NS histone family protein, known as regulators act to modulate flagellar production (38), were identified as genes directly regulated by BysR in our DAP-seq (Fig. 4E, Supplementary Table S5), and implying that BysR may indirectly regulate flagellar assembly by directly regulating Lrp and H-NS protein expression. Previous study had shown that *ocf* genes (including *ocfA-N* and *ambR2*), flagellar assembly-related genes (including *fliT*, *fliD* and *motB*) and the genes associated with type VI secretion system (including *hcp*, *tssC* and *tssB*) were significantly down-regulated in *ambR1* mutant (MS455MT38) compared to the wild type *Burkholderia* sp. MS455 revealed by RNA-seq analysis (2), which are consistent with our study that *bysR* defective mutant affect the expression of *ambR1*, and in turn affects the expression of *ocf* gene cluster, flagellar assembly and type VI secretion system related genes. Therefore, based on the results of this study and previous study, we infered that *ambR1* not only regulates occidiofungin production, but also modulates other cellular processes, and AmbR1 is a downstream component of BysR in the regulatory network, and plays a direct role in regulating flagellar assembly and type VI secretion system related genes expression. However, more studies are needed to confirm the speculation that *ambR1*, as a downstream regulator of BysR, directly regulates genes involved in flagellar assembly and type VI secretion system.

DAP-seq as an alternative and a complementary method to ChIP-seq, it is powerful and convenient. Due to it is not limited to one specific growth condition, DAP-seq usually can reveal the binding events under conditions that are not suitable for ChIP-seq (26). DAP-seq has been successfully applied to identify TFBSs in several organisms, especially in eukaryotes, such as *Arabidopsis thaliana* (26), rice (39, 40) and maize (41). However, DAP-seq was not widely used in prokaryotes, and only several reports have been reported in the study of TFs in *Pseudomonas*, such as the studies on XRE-like family TFs and two-component systems response regulators in *Pseudomonas aeruginosa* (42, 43). More studies in different species will be encouraged to evaluate the applicability of DAP-seq in prokaryote. In this study, we applied DAP-seq in *Burkholderia* to identify the binding sites of BysR in the genome of JP2-270 for the first time and the results uncovered 400 putative binding loci of BysR (Supplementary Table S5). And most remarkably, the genes encoding transcriptional regulators are the largest number of gene group directly regulated by BysR (Fig. 4E), and DM992_17470 (*bysR*), DM992_33405 (*ambR2*), DM992_33410 (*ambR1*), DM992_38895 and DM992_39970 were also found down-regulated in Δ*bysR* as revealed by RNA-seq (Fig. 3A and 4E, Supplementary Table S3). Interestingly, an unidentified MarR family transcriptional regulator DM992_38895, located in the gladiostatin biosynthesis gene cluster (Region 3.7 in Table 1), was identified as the target of BysR in RNA-seq and DAP-seq (Fig. 3A and 4E). This suggests that, similar to *ambR1*, BysR may control the expression of genes responsible for gladiostatin production by directly binding to the promoter region of DM992_38895 in JP2-270. However, more experimental evidence is needed to confirm this assumption. Besides the genes related to the production of occidiofungin, our analyses suggested BysR regulates a variety of cellular processes, such as primary and secondary metabolism, drug resistance, transport, mobility, transposase activity, transcriptional regulation and quorum sensing system (Fig. 4E and Supplementary Table S5), indicating that BysR is a global and pleiotropic transcriptional regulator in *Burkholderia* sp. JP2-270. Previously characterized target genes regulated by other LTTRs from other bacterial species using ChIP-seq, ChIP-on-chip or genome scale SELEX were also found in our DAP-seq results, such as genes related to type VI secretion system, H-NS family transcription regulator and multi-drug efflux transporter (28, 44, 45), indicating that the results of our DAP-seq are credible and some of the regulatory functions in cellular processes are shared by LTTRs from different microorganisms. Nevertheless, it is distinctive from previous reports, as a higher level of transcriptional regulator, BysR could regulate various types of transcriptional regulators, such as LTTRs, LuxR family regulators, MarR family regulators, H-NS family regulators, IclR regulators, GntR family regulators, Lrp/AsnC family regulators, TerR/AcrR family regulators and so on.

Of note, the diaminobutyrate-2-oxoglutarate transaminase (DABAT, EC2.6.1.76) encoded by gene *ocfL* (DM992_33335) presumably catalyzes the formation of L-2,4-aminobutyric acid (DABA) and 2-oxoglutarate (46). L-2,4-aminobutyric acid was added to the intermediate of occidiofungin by OcfE (12), and 2-oxoglutarate may be participate in the TCA cycle. The *ocfL* may not only provide aminobutyric acid during the synthesis of occidiofungin, but also supplement the source of 2-oxoglutarate in the TCA cycle. Thus, we predicted that *ocfL* may be involved in both primary and secondary metabolic processes. In addition to the regulatory mechanisms related to secondary metabolite synthesis, we need more studies to depict the other function of BysR. A deeper research about the other function of BysR will provide better understanding on how Bcc isolates adapt to the environmental conditions.

Similar to ScmR and ShvR, as the global regulator, BysR not only regulates multiple cellular processes (including DNA replication, carbohydrate metabolism, amino metabolism, TCA cycles, drug resistance, transport, mobility, transposase activity, transcriptional regulation and quorum sensing system), but also participates in the regulation of secondary metabolites (20, 23). The pathway specific transcriptional regulatory genes *ambR1* is a key target gene of BysR, and BysR activates the transcription of *ocf* gene cluster by directly binding the promoter region of *ambR1*. Aside from *Burkholderia*, LTTRs function as a global and pleiotropic transcriptional regulator had been reported in other genera, such as PsrA in *Serratia Marcescens*, LeuO in *Escherichia coli* and BsrA in *Pseudomonas aeruginosa* (28, 36, 37). The cellular processes related to secondary metabolism, virulence, drug resistence, transporter and motility may be the common regulation targets of these LTTRs as revealed by previous studies and this study (20, 28, 36, 37, 44, 47, 48).

## MATERIALS AND METHODS

### Bacterial/fungal strains and culture conditions

Bacterial, fungal strains and plasmids used in this study are listed in Supplementary Table S1. The strain *Burkholderia* sp. JP2-270 and derivates were routinely cultured in Luria-Bertani medium (LB medium) at 28 °C (49). *Escherichia coli* strains were routinely maintained in LB medium at 37 °C. The concentrations of antibiotics, when necessary, used for *E. coli* were 50 μg/ml for carbenicillin, 50 μg/ml for kanamycin, 50 μg/ml for streptomycin and 30 μg/ml for nalidixic acid. *M. oryzae* isolate Guy11 and *R. solani* GD118 were routinely cultivated on complete medium (CM medium) and PDA plates, respectively, at 25 °C.

### *In vitro* inhibition assay

The petri dish assay was used to test *in vitro* antagonistic activity of JP2-270 and derivates against *M. oryzae* Guy11 and *R. solani* GD118. The overnight culture of JP2-270 and derivates were streak inoculation at both sides, 2 cm away from the center of CM or PDA plates. After incubation for 24 h at 25 °C, the mycelium plugs (5-mm-diameter) from the fresh edge of *M. oryzae* Guy11 or *R. solani* GD118 were placed at the center of the plates. Following 2 to 10 days co-culturing at 25 °C, the radius of mycelium growth were measured to evaluate the inhibitory effect. Five biological replicates were performed and an average value was calculated to evaluate the inhibitory activity.

### Overexpression of *ambR1* and *ambR2* in Δ*bysR*

To construct plasmid expressing *ambR1* gene, the fragment containing coding sequences of *ambR1* (DM992_33410) from JP2-207 genome was cloned into a multiple cloning site of pBBR1MCS-2. The plasmid pBBR2-*ambR1* expressing *ambR1* was obtained. The primers C*ambR1*-F/C*ambR1*-R were used to amplify the corresponding fragment. All constructs were verified by PCR and sequencing. Similary, the pBBR2-*ambR2* expressing *ambR2* gene was constructed. The obtained plasmids were electroporated into Δ*bysR* to obtain the *ambR1*-overexpression and *ambR2*-overexpression strain *ambR1^+^* and *ambR2^+^,* respectively. The primers used in this study were listed in Table S2.

### In-frame *ocfE*, *gdsE* and *prnC* gene deletion in JP2-270

The marker free site-directed deletion was carried out using pK18mobSacB (50). The upstream and downstream regions of *ocfE*, *gdsE* and *prnC* to be deleted were fused using overlap extension PCR. The fusion products were then subcloned into the suicide vector pK18mobSacB. The resultant recombinant plasmids were introduced into JP2-270 by electroporation transformation, and subsequently the plasmids were integrated within the target gene via homologous recombination. *Burkholderia sp.* JP2-270 is not sensitive to nalidixic acid. *Burkholderia* isolates containing the plasmids were selected on LB with 50 μg/ml kanamycin, 30 μg/ml nalidixic acid. The *Burkholderia* isolates containing plasmids were subsequently cultured in LB without any antibiotic for several generations. The deletion mutants that grew on LB plate with 30 μg/ml nalidixic acid but did not grew on LB plate with 50 μg/ml kanamycin, 30 μg/ml nalidixic acid were selected, and the resultant allelic exchange mutants of *ocfE* (Δ*ocfE*), *gdsE* (Δ*gdsE*) and *prnC* (Δ*prnC*) were verified by PCR and subsequent DNA sequencing. The primers used in this study were listed in Table S2.

### RNA isolation, RNA-seq and quantitative RT-PCR

*Burkholderia* sp. JP2-270 (WT) and in-frame *bysR* deletion isolate (Δ*bysR*) strains were grown in LB medium at 28 °C. An overnight cultures (1%) were inoculated into CM medium and grown at 28 °C. The cells were harvested after 24 h incubation. Three biological replicates of each strain were used. The collected cells were sent directly to the company (Beijing Novogene Bioinformatics Technology Co., Ltd.) for further treatments. Briefly, a total of 3 µg RNA per sample was obtained and rRNA were depleted using Ribo-Zero rRNA removal kit (G^-^ bacteria). The mRNA were fragmented using diavalent cations under elevated temperature in NEBNext First Strand Synthesis Reaction Buffer (5 X). Then, the cDNA libraries were generated using NEBNext Ultra Directional RNA Library Prep Kit for Illumina (NEB, USA) following manufacture’s instructions. The cDNA fragments were purified with AMPure XP system (Beckman Coulter, Beverly, USA), and then 3 µl USER Enzyme (NEB, USA) was used with size-selected, adaptor-ligated cDNA at 37 °C for 15 min followed by 5 min at 95 °C before PCR. Then PCR was performed with Phusion HighFidelity DNA polymerase, Universal PCR primers and Index Primer. At last, products were purified (AMPure XP system) and library quality was assessed on the Agilent Bioanalyzer 2100 system. The resultant samples were sequenced on an Illumina Hiseq 2000 platform.

For qRT-PCR analysis, an RNA prep pure cell/bacterial kit (Tiangen Biotech, Beijing, China) was used to prepare total RNA according to the instructions. The quality and concentration was analyzed by 1% agarose gel electrophoresis. Genomic DNA (gDNA) was digested with gDNA remover (TOYOBO) at 37 °C for 5 min. Total RNA (∼0.5 μg) was reversed to cDNA using a ReverTra Ace qPCR RT Master Mix (TOYOBO). Samples were diluted and about 25 ng of cDNA were added to the PCR system in a total volume of 20 μl, and quantitative PCR (qPCR) was conducted using THUNDERBIRD SYBR qPCR Mix (TOYOBO). The qRT-PCR was performed in an Applied Biosystems 7500 according the manufacturer’s instructions. The *recA* gene was used as an internal standard, and relative expression was quantified using the 2^-ΔΔCt^ threshold cycle (Ct) method. Primers are listed in Table S2. Three independent experiments were conducted, each with three replicates.

### DAP-seq analysis

DAP-seq was carried out in duplicate on *Burkholderia* sp. JP2-270 genomic DNA as previously described (26). Specifically, the Halo-BysR protein was expressed using the TNT SP6 Wheat Germ Protein Expression System (Promega, Fitchburg, WI, USA). The Halo Tag-BysR protein was purified using magnetic HaloTag beads (Promega) and verified by Western blotting with the anti-HaloTag antibody (Promega). Genomic DNA was extracted from JP2-270 using DNAiso (Takara, Japan). Genomic DNA library was generated using NEXTflex Rapid DNA Seq kit (Bioo Scientific, USA) following the manufacturer’s instructions. The purified protein was incubated with DNA library at 30 °C for 2 h, and then the unbound DNA fragments were washed away. Then, the BysR-bound DNA fragments were recovered and PCR-amplified. The resultant products were sequenced using the Illumina NovaSeq platform with the sequencing strategy of PE150. The two technical duplicates were performed and the DNA sample without incubated with BysR protein was used as input negative control.

### Bioinformatics analysis

For RNA-seq data, clean reads were obtained by removing reads containing adapter, reads containing ploy-N and low quality reads from raw data. Clean reads were mapped to the *Burkholderia* sp. JP2-270 reference genome (CP029824-CP029828) using Bowtie 2-2.2.3 (51). Then Htseq v0.6.1 was used to count the reads numbers mapped to each gene (52). FPKM (Fragment Per Kilobase of transcript sequence per Millions base pairs sequenced) was used to estimate gene expression levels (53). Differential expression analysis of two groups (three biological replicates per group) was performed using the DESeq R package (1.18.0). The square (R^2^) of Pearson correlation coefficient was caculated to estimate the reproducibility of the replicates. DESeq provide statistical routines for determining differential expression in digital gene expression data using a model based on the negative binomial distribution. The resulting p-values were adjusted using the Benjamini and Hochberg’s approach for controlling the false discovery rate. Genes with an adjusted p-value < 0.05 and |log2(FoldChange)| > 1 found by DESeq were assigned as differentially expressed. The volcano plot drawn by Origin 2022 was applied to display significantly different gene expression. KEGG (Kyoto Encyclopedia of Genes and Genomes) enrichment analysis was performed by KOBAS software (54).

For DAP-seq analysis, the clean reads were obtained by filtering the raw data using FASTP with the default parameters. The clean reads were mapped to the *Burkholderia* sp. JP2-270 reference genome (CP029824-CP029828) using BWA-MEM (55). Peaks were generated with MACS2 (56) (p-value < 0.01), and IDR software were used to merge the peaks present in the two replicates. The MEME-CHIP was used to analyze the conservative motifs in the peaks (57). The HOMER software (58) was used to annotate peaks. The box plots of the peaks width distribution, the pie chart of distribution of BysR bound genes and the histogram-distance to TSS (Transcription Terminate Site) were drew using Origin 2022. Venn plot for RNA-Seq and DAP-Seq data was produced by Origin 2022.

### Protein expression, purification and EMSA

The coding region of BysR was obtained by PCR with the primers BysR-F/BysR-R listed in Table S2 and cloned into vector pGEX-6P-1 to generate a N-terminally GST-tagged BysR protein fusion. The resulting plasmid was transformed into *E. coli* BL21(DE3) (Table S1) for protein expression. The resultant strain was cultivated in LB medium (containing 100 μg/mL ampicillin) overnight at 37 °C. Two millilitres of the overnight culture was transferred into 200 mL fresh LB at 37 °C and grown with shaking at 200 rpm, until OD_600_ = 0.4 ∼ 0.6. Isopropyl-β,D-thiogalactopyranoside (IPTG, Sangon, Shanghai, China) was added to a final concentration of 0.4 mM. The culture was incubated for an additional 16 h at 18 °C with shaking at 100 rpm. Cells were collected by centrifugation at 4 °C, resuspended in 25 mL phosphate buffered saline (PBS) lysis buffer containing 10 mM protease inhibitor (PMSF, Beyotime, China), and lysed by sonication (Scientz, JY92-IIDN, Ningbo, China)(power 200 W, ultrasonic 5 s, intermittent 10 s). Following centrifugation at 4500 rpm at 4 °C for 30 min, the soluble protein was collected by incubation with GST beads (Sangon, Shanghai, China) for 30 min. The purified GST-BysR protein was eluted with buffer containing glutathione. According to the instructions, the GST tag was removed by PreScission protease (P2303, Beyotime, Shanghai, China). The purity of BysR protein was assessed by sodium dodecyl sulphate– polyacrylamide gel electrophoresis (SDS-PAGE) and the concentration of BysR determined using a BCA assay kit (Sangon Biotech, Shanghai, China). Aliquots of the proteins were stored at −80 °C.

EMSA (Electrophoretic mobility shift assay) was performed as follows. Cy5-labelled probes of the *ambR1* promoter region were obtained by PCR using EMSA-ambR1-F/EMSA-ambR1-R (for probe 1), EMSA-ambR2-F/EMSA-ambR2-R (for probe 2) and EMSA-ambR3-F/EMSA-ambR3-R (for probe 3), respectively. Probe and protein extract were mixtured according to the specifications of EMSA/Gel-shift binding buffer (5×) (GS005, Beyotime, Shanghai, China), and incubated for 20 min at room temperature under darkness. The binding mixture was loaded onto the 6% non-denatured polyacrylamide gel and electrophoresed at 100 V for 4 h in dark. Odyssey CLx infrared fluorescence imaging system (LI-COR, Lincoln, NE, USA) was used to detect the fluorescence signal of Cy5-labeled DNA fragments. Probes used were listed in Table S2.

### Statistics

Data analysis was performed using Microsoft Excel 2010. The Student’s T-test was used to compare the differences between two sets of data. The results were considered to be statistically significant different when *p* < 0.05 and extremely significant difference when *p* < 0.01.

### Availability of supporting data

Raw sequence data from our RNA-seq and DAP-seq analysis can be accessed via the National Center for Biotechnology Information Sequence Read Archive server under accession number GSE193778 and GSE193916, respectively, and the complete genome sequence of *Burkholderia* sp. JP2-270 has been deposited in NCBI under accession number CP029824 to CP029828. We thank Dr. Guozhong Feng (CNRRI, Hangzhou, China) for platform support provided in the initial stage of this research. We thank Prof. Changfu Tian (College of Biological Sciences, China Agricultural University, Beijing, China) for his constructive comments on this munuscript.

## SUPPLEMENTAL MATERIAL

Supplemental material is available online only.

## ACKNOWLEDGEMENTS

This research was financially supported by National Natural Science Foundation of China (NSFC) (31901924, 31972318) and supported by Zhejiang Provincial Key Research and Development Program of China (2021C02056-3), Zhejiang Provincial Natural Science Foundation of China (LY21C130004). We thank Dr. Guozhong Feng (CNRRI, Hangzhou, China) for the platform support in the initial stage of this research. We thank Prof. Changfu Tian (College of Biological Sciences, China Agricultural University, Beijing, China) for his constructive comments on this manuscript.

## Supplementary Information

**Supplementary Figure 1:**
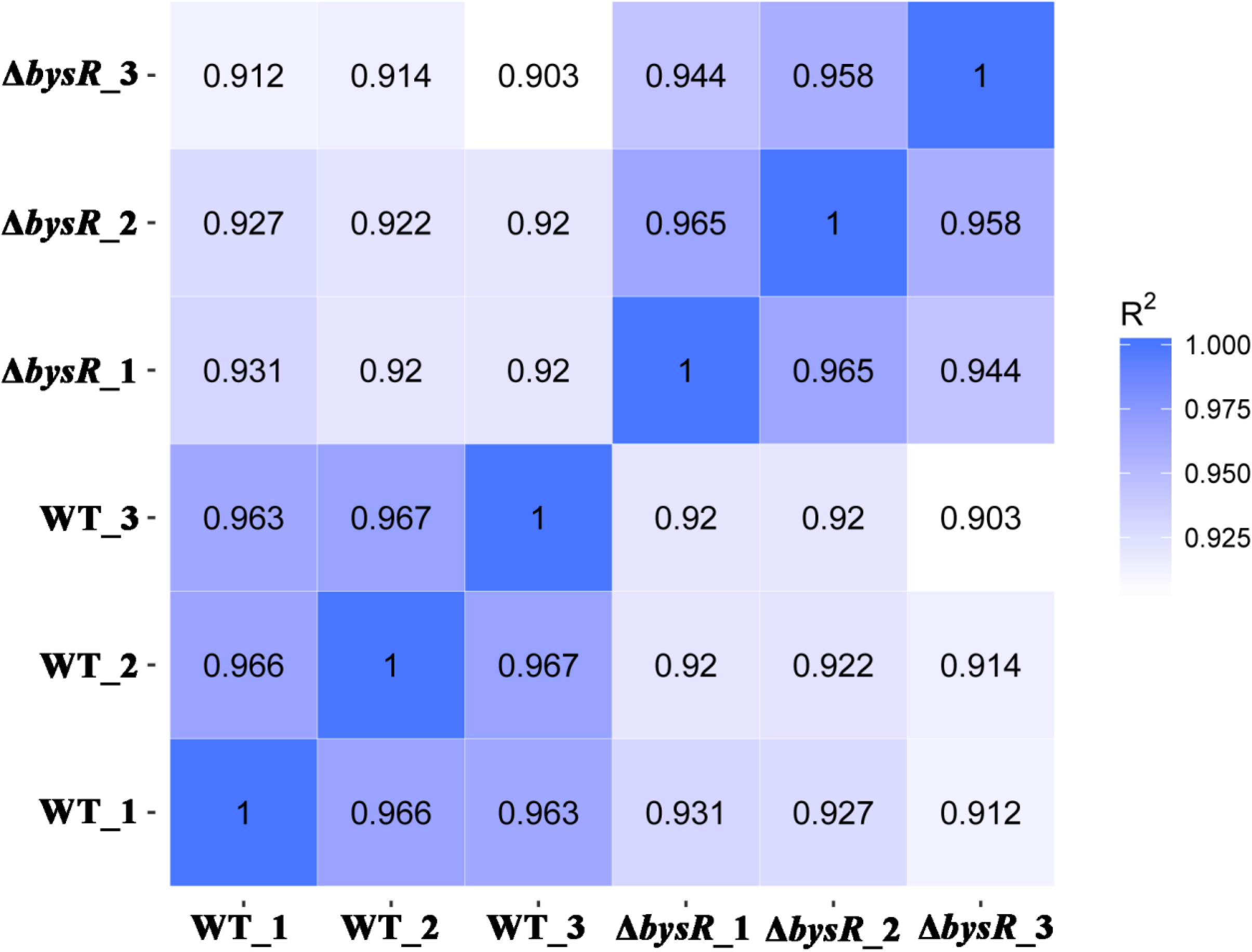
The pearson correlation coefficient between replicates of RNA-seq. WT_1,WT_2,WT_3 and Δ*bysR_*1, Δ*bysR_*2, Δ*bysR_*3 represent the three replication of wild type isolate JP2-270 and *bysR* deletion isolate Δ*bysR*, respectively.

**Supplementary Figure 2:**
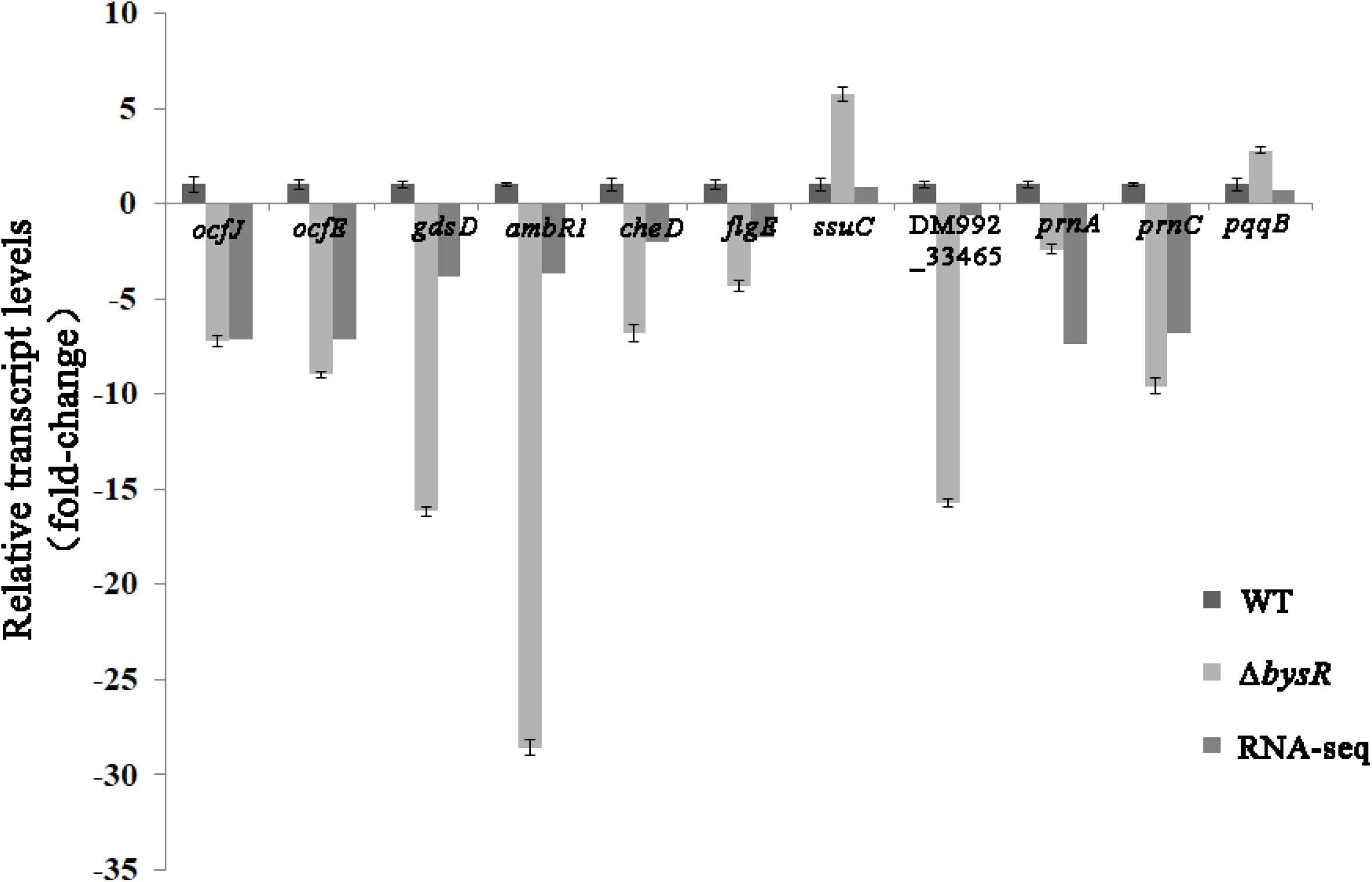
The transcription levels of representative genes in WT, Δ*bysR* and RNA-Seq results. The *recA* gene is used as a control. The relative gene expression levels are presented as expression ratios of the indicated genes.

---

**Supplementary Table S1:**
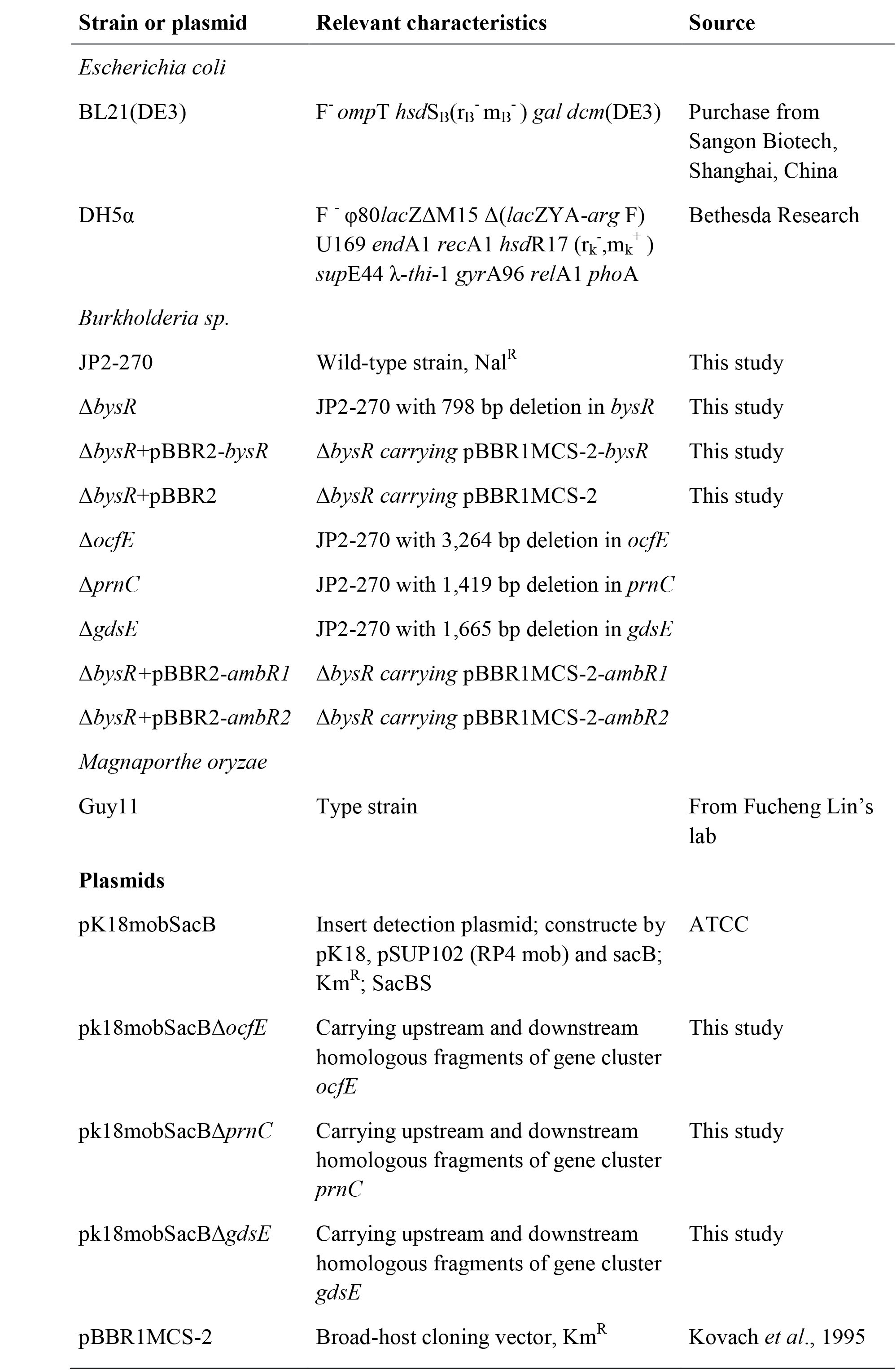
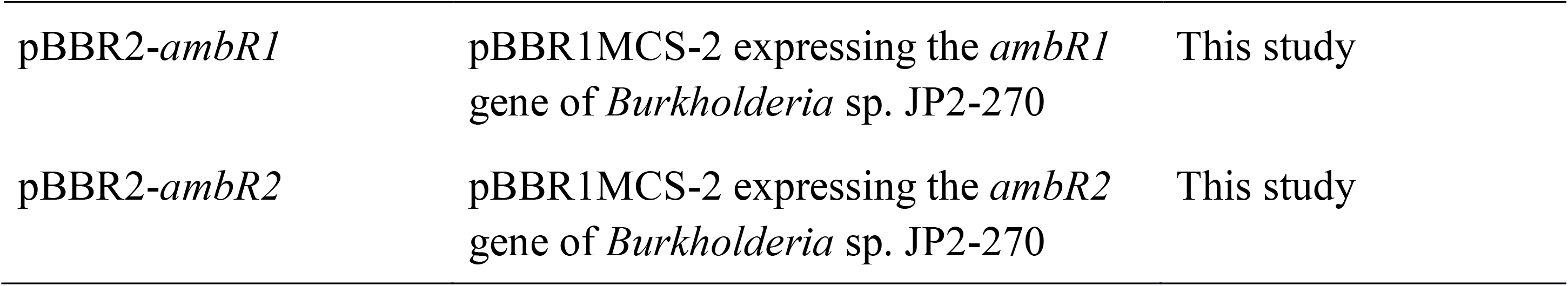
Bacterial strains and plasmids in this study

**Supplementary Table S2:**
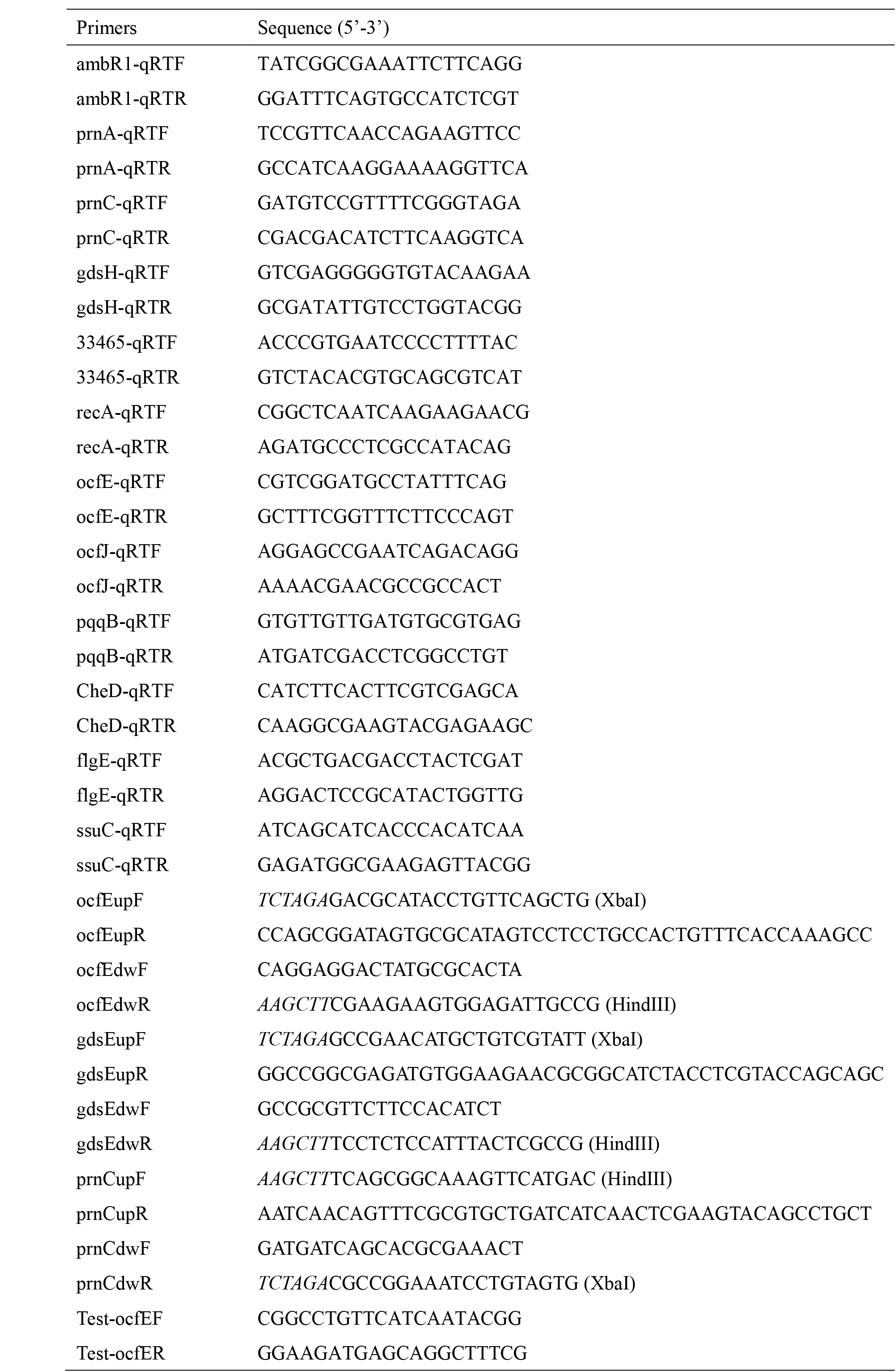
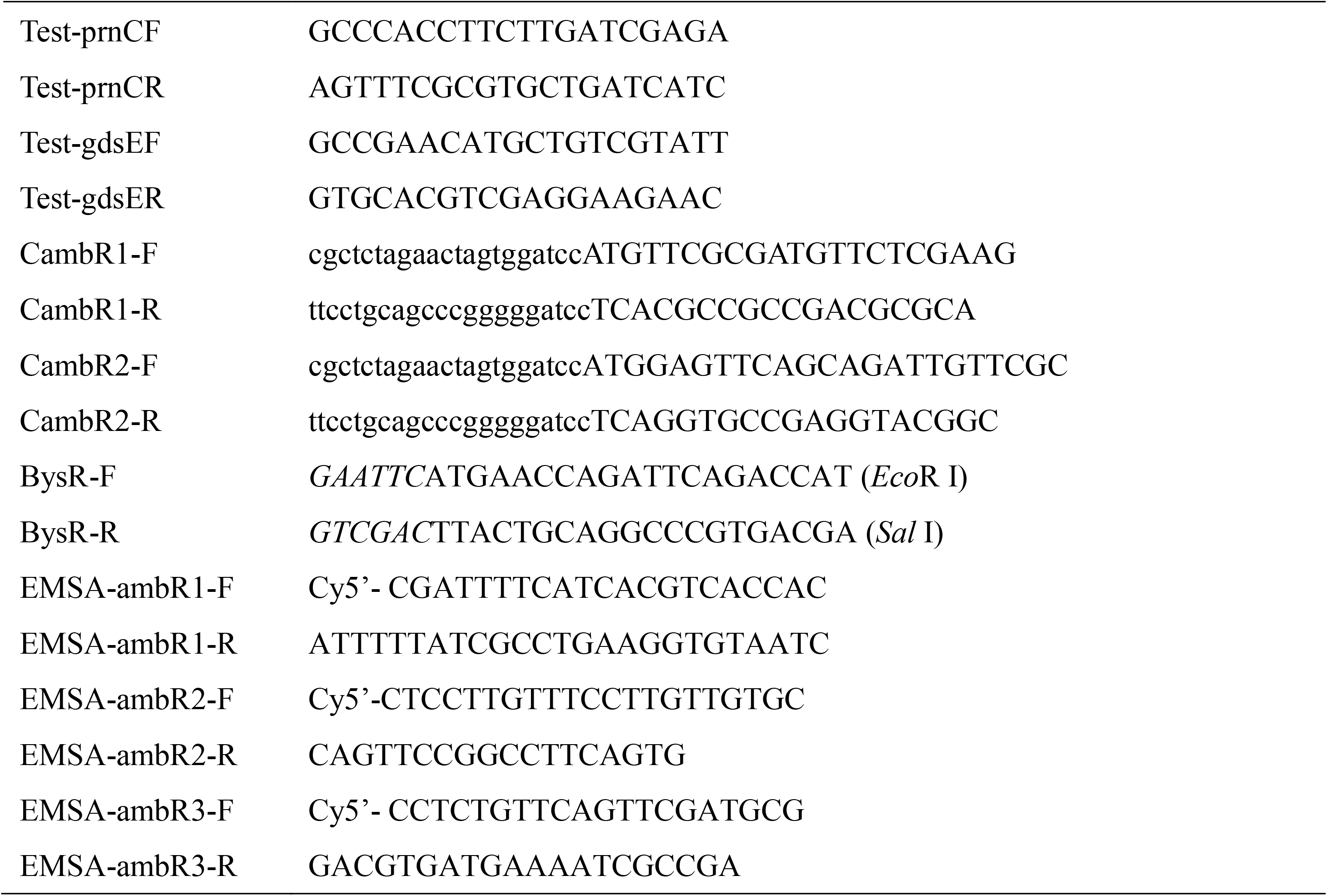
Primers used in this study

**Supplementary Table S3.**
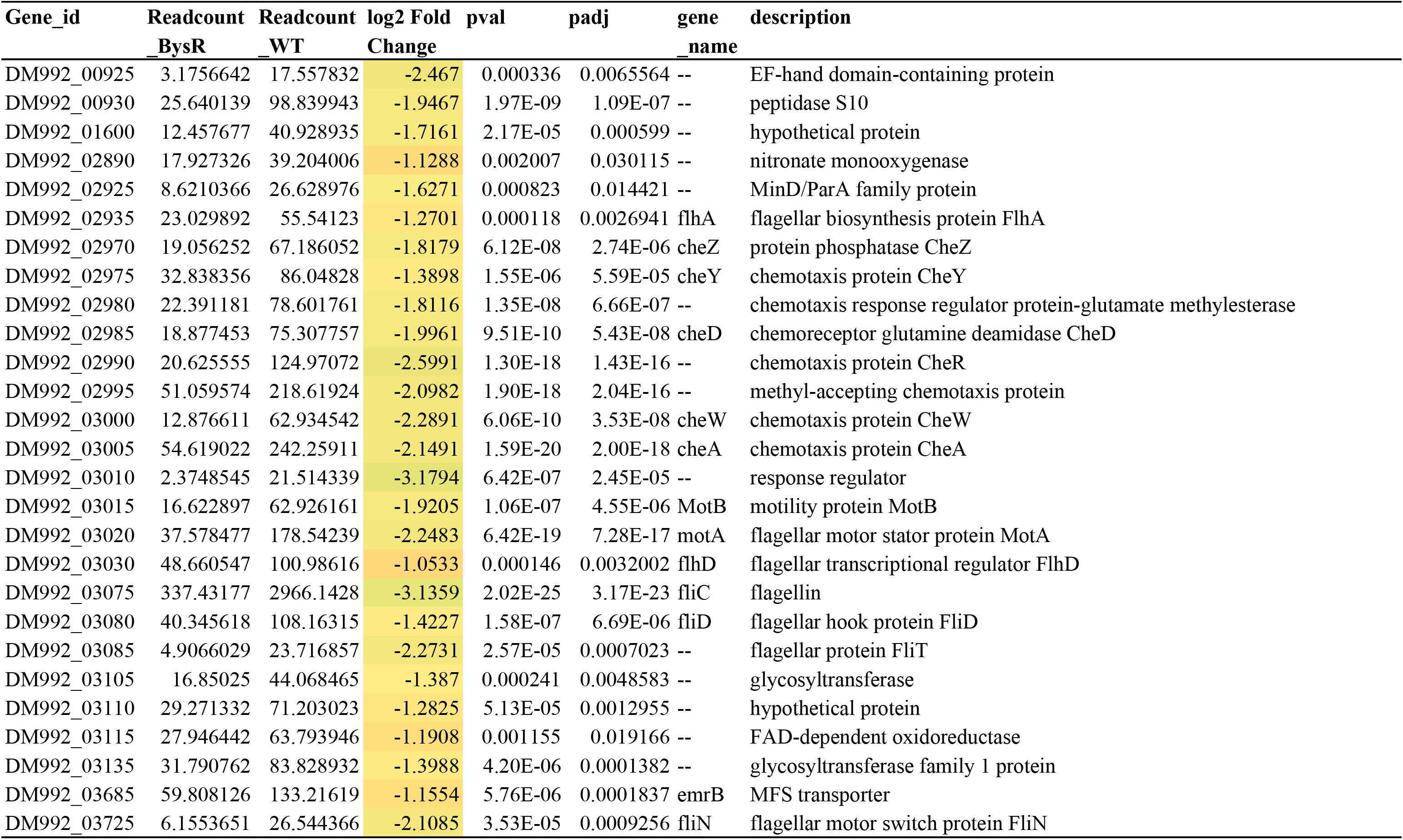
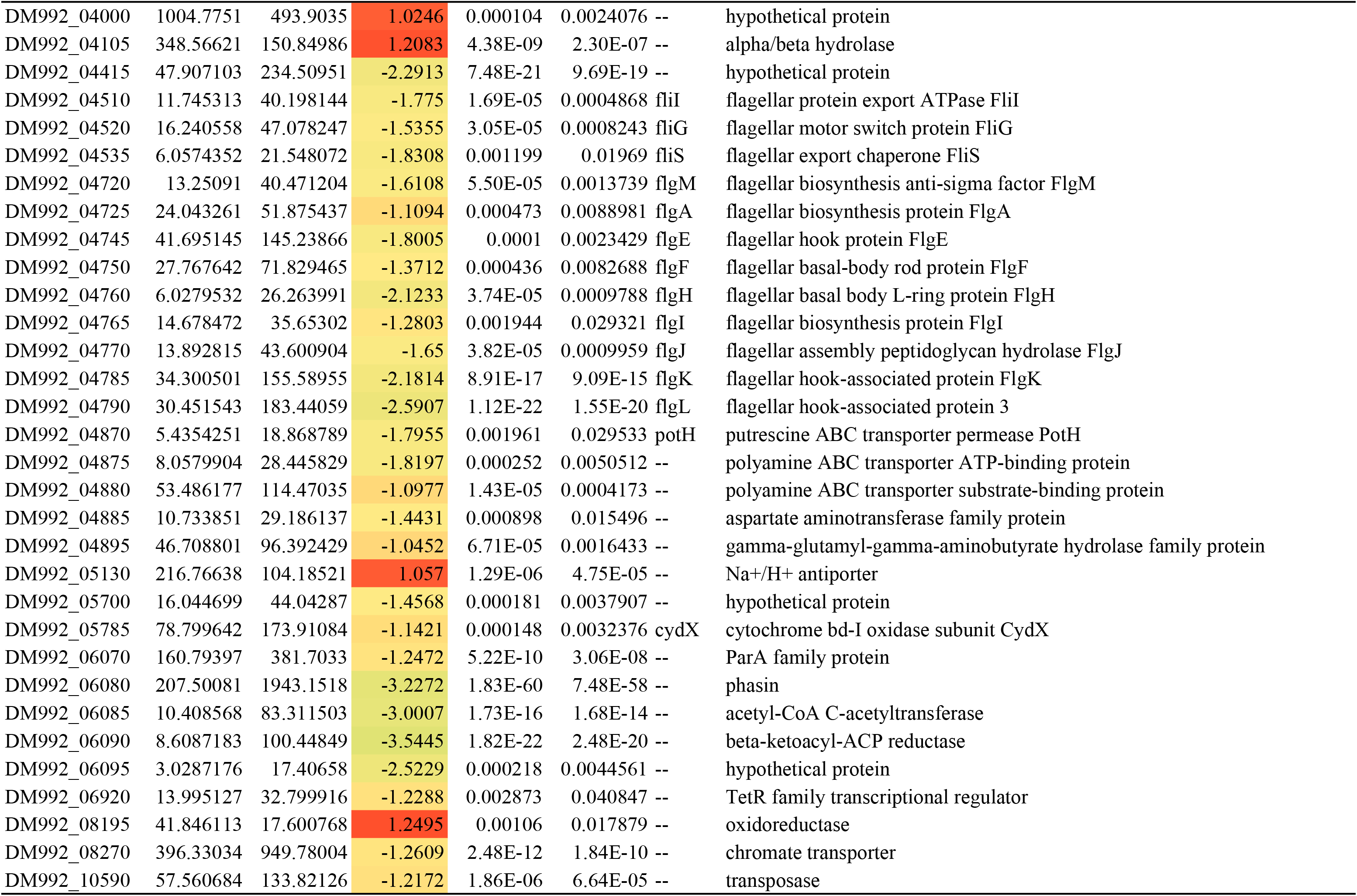
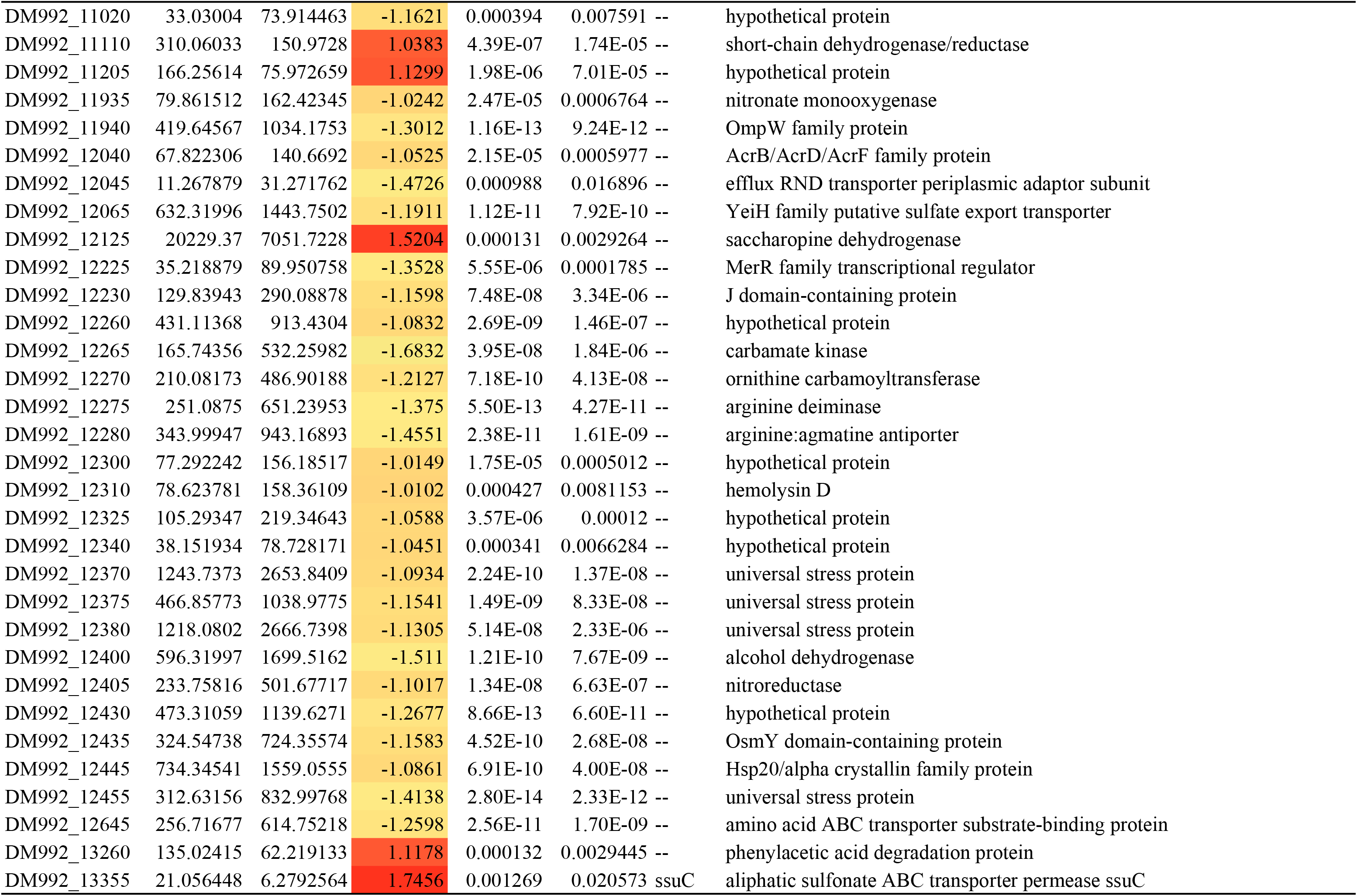
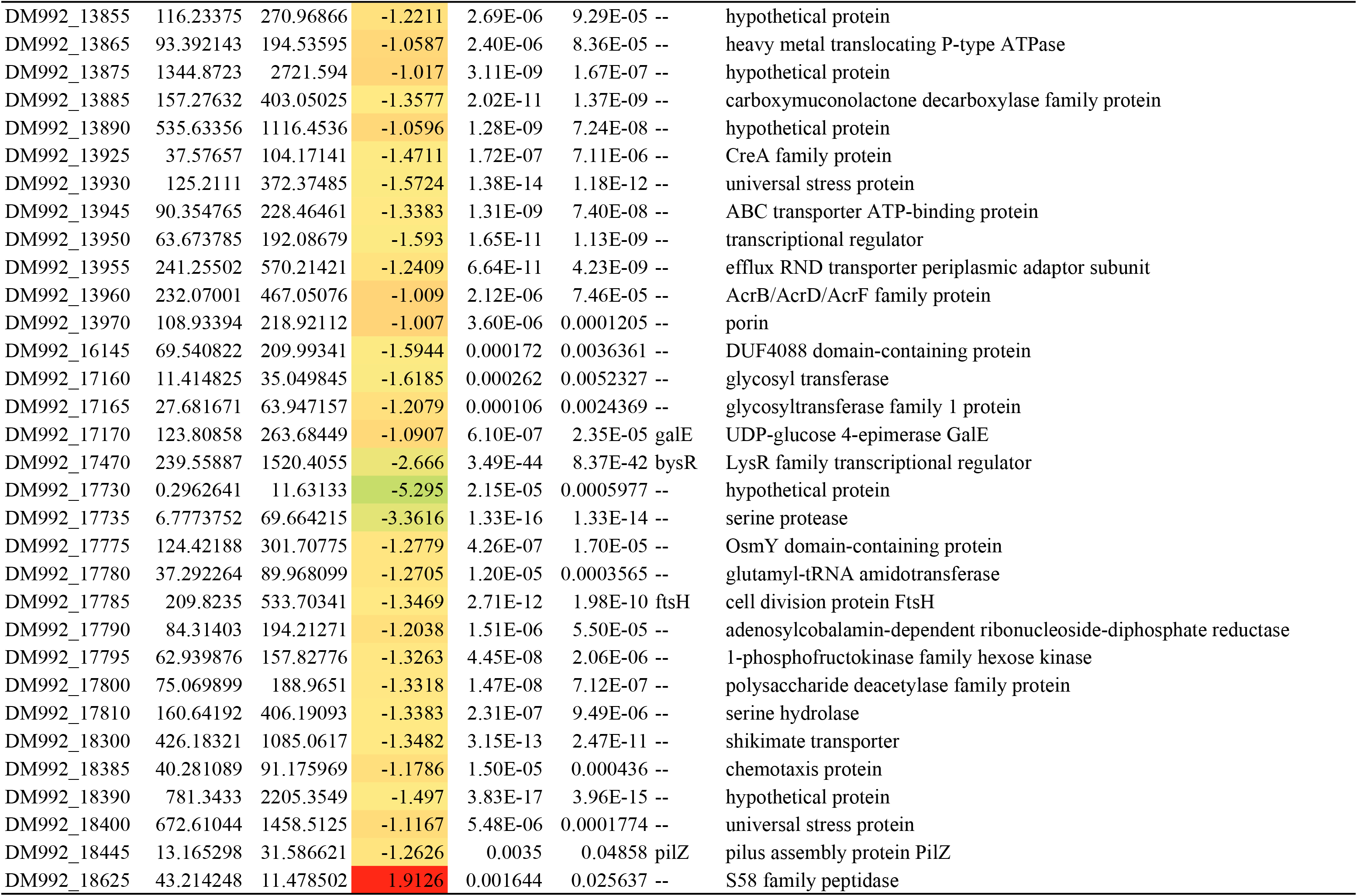
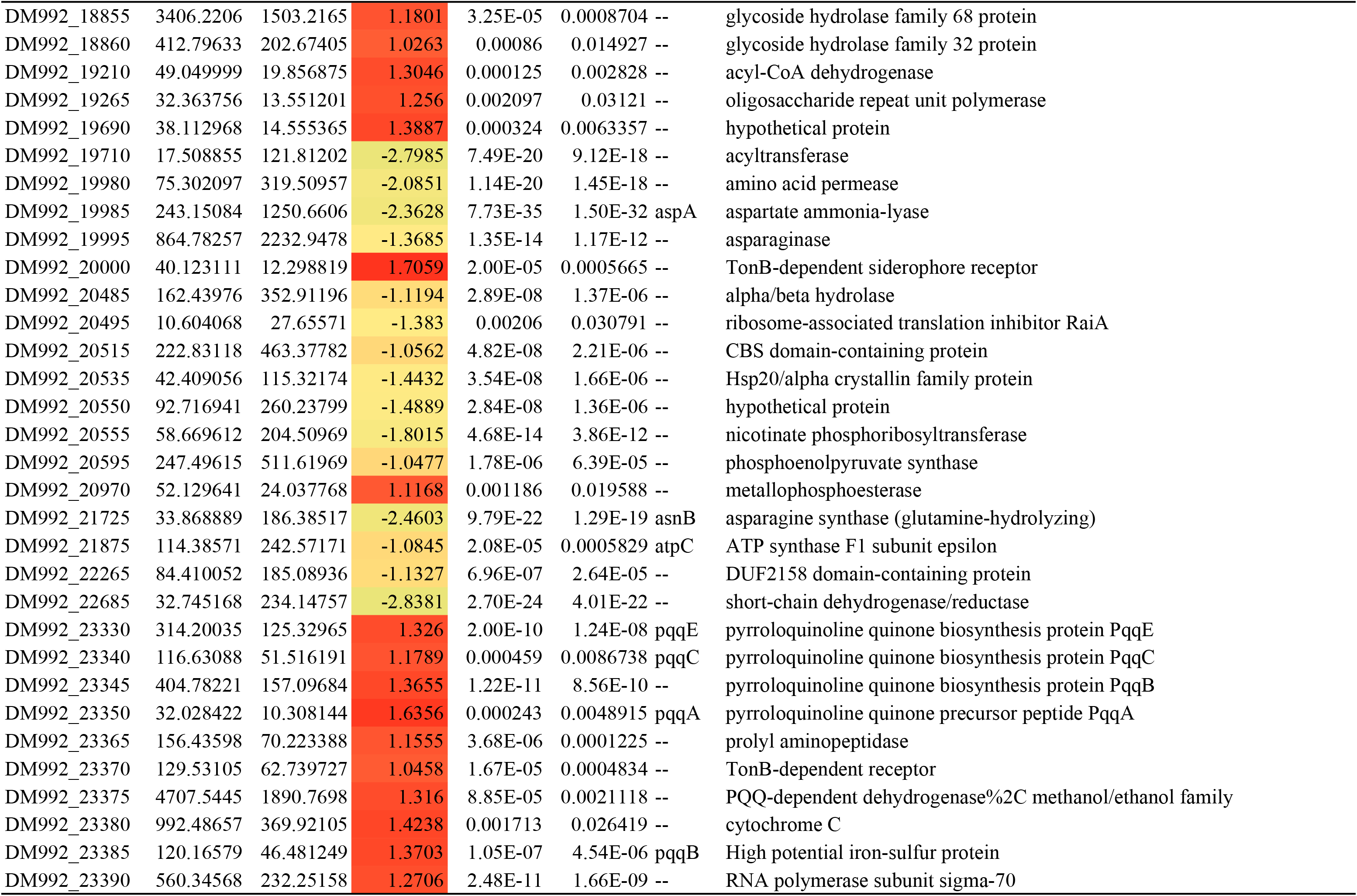
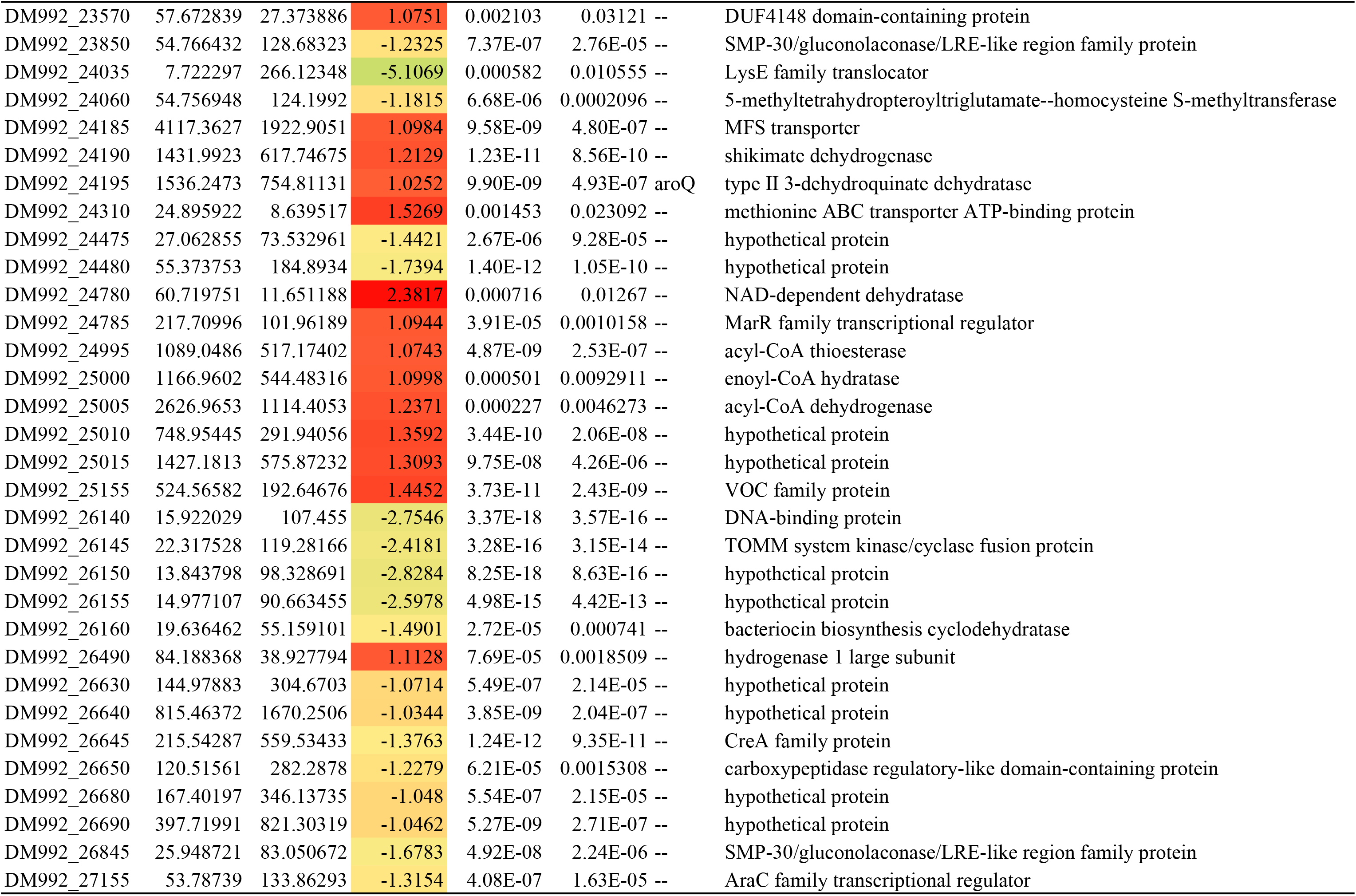
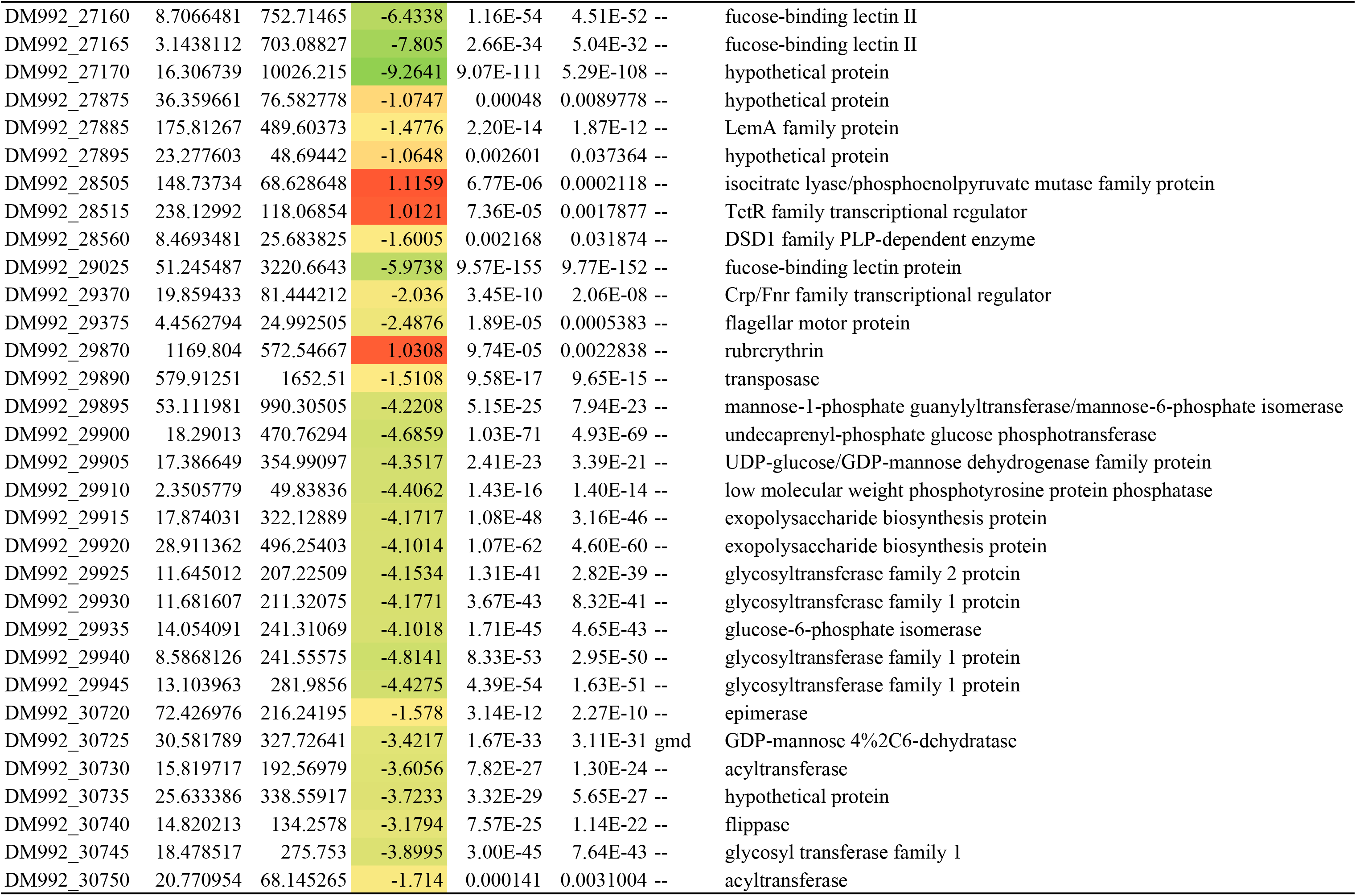
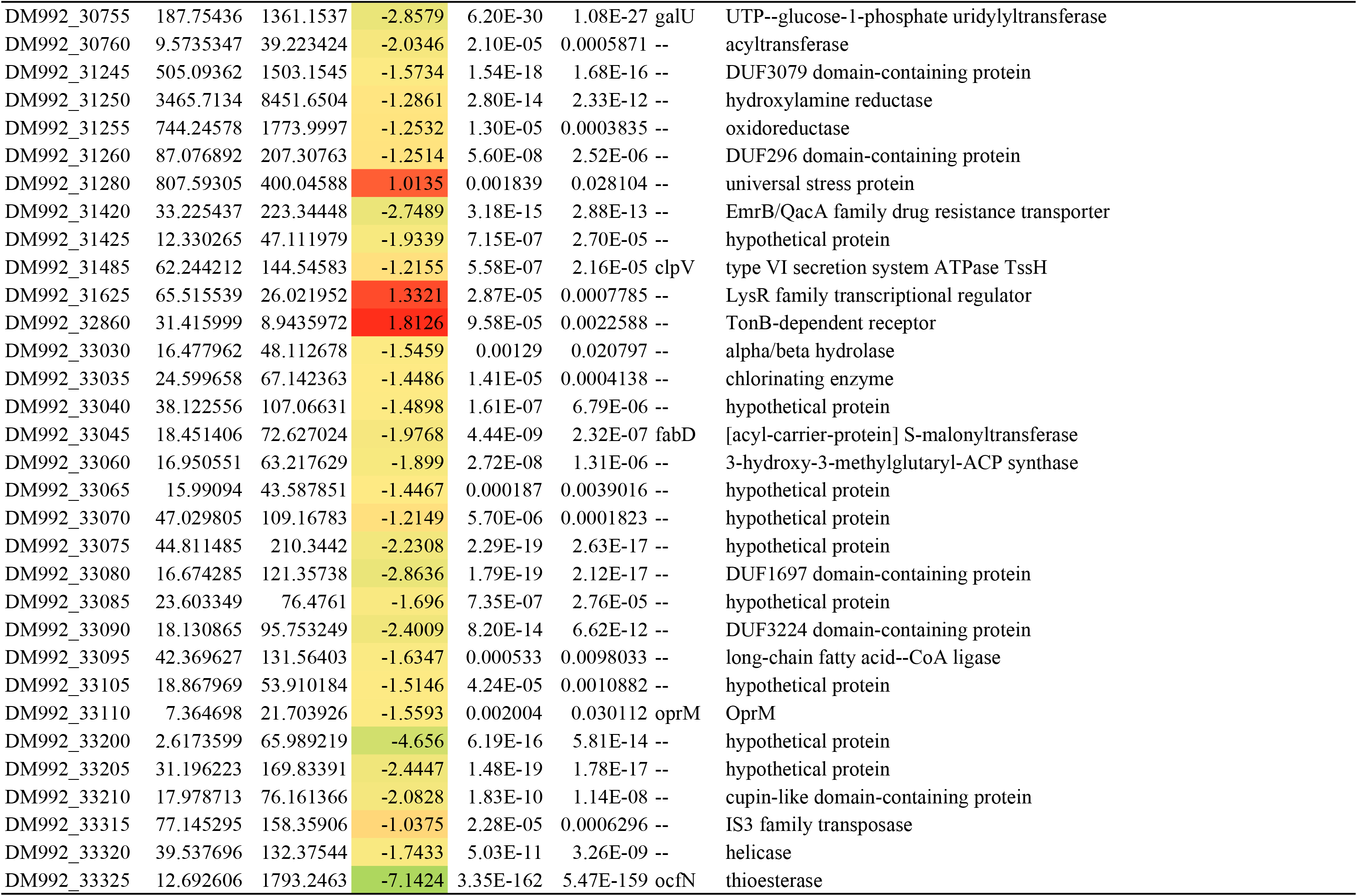
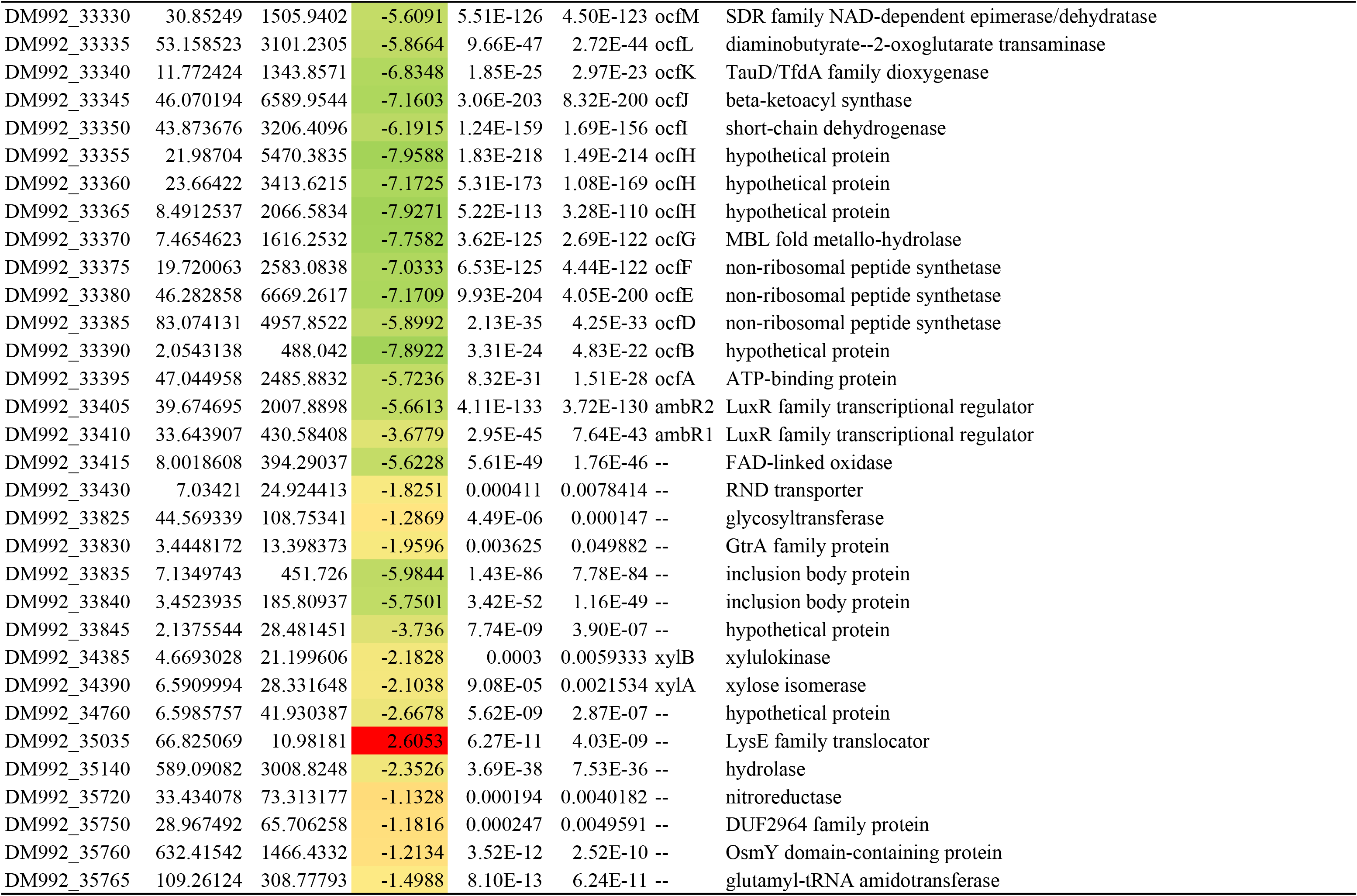
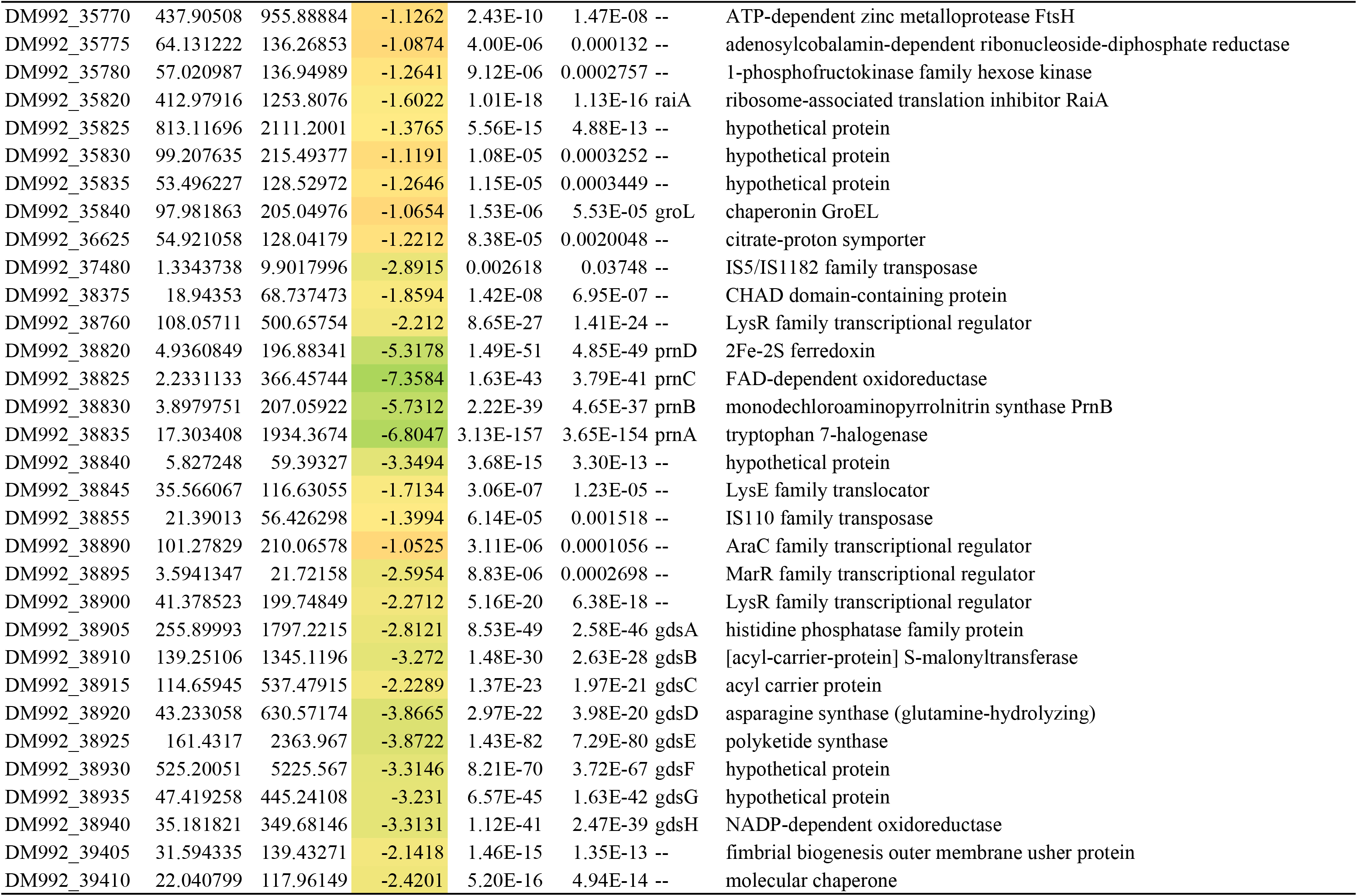
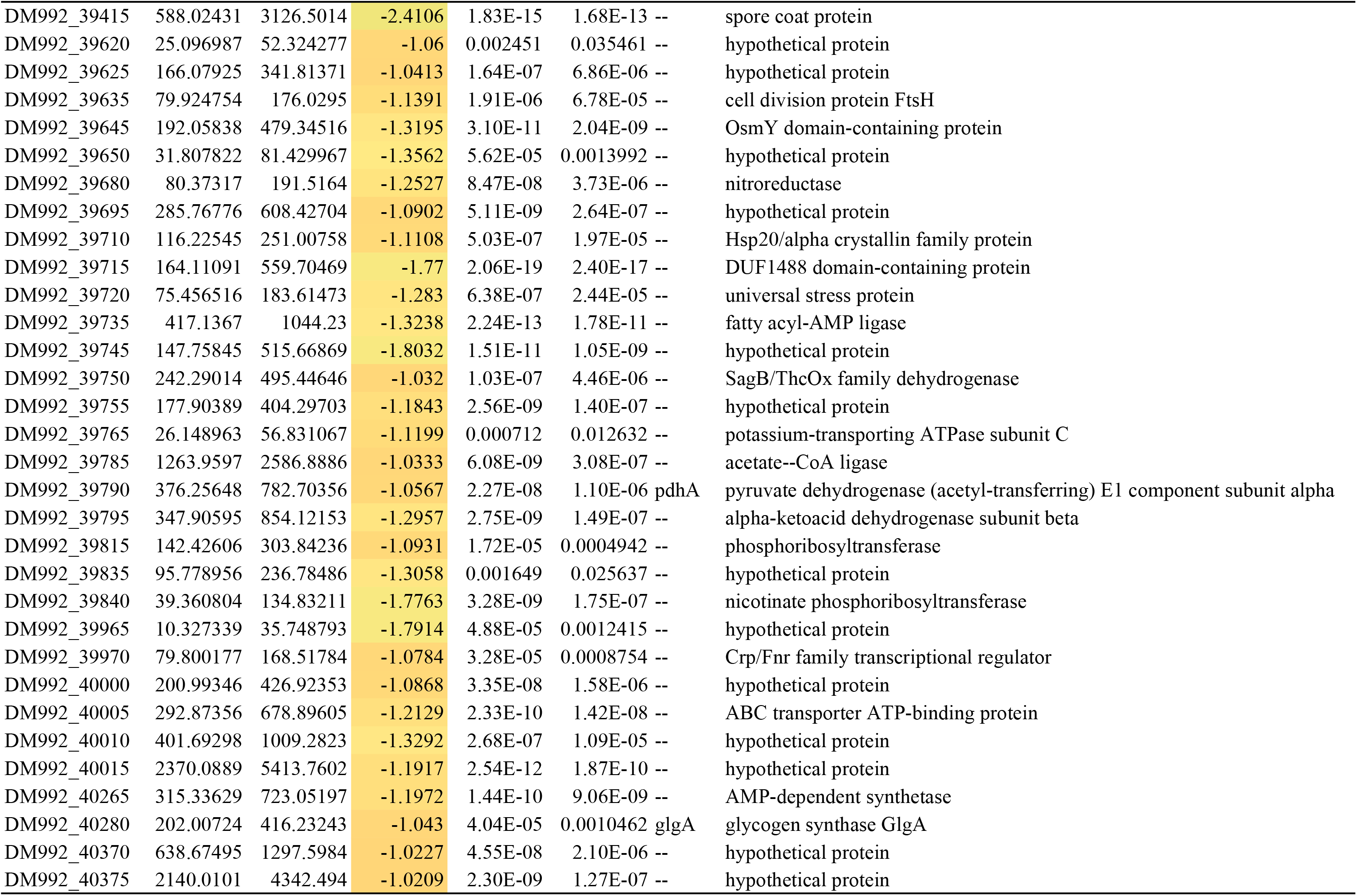
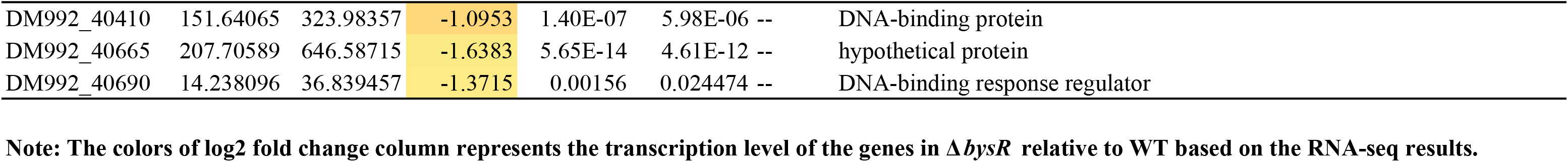
The DEGs in Δ*bysR* relative to WT

**Supplementary Table S4.** The KEGG pathway enrichment results

| Term | ID | Input<br>number | Background<br>number | P-Value | Corrected<br>P-Value | Input |
| --- | --- | --- | --- | --- | --- | --- |
| Flagellar assembly | bct02040 | 20 | 38 | 5.77E-11 | 3.69E-09 | DM992_03030\|DM992_03080\|DM992_04500\|DM992_04720\|DM992_04535\|DM992_04785\|DM992_04765\|DM992_03075<br>DM992_04720\|DM992_04535\|DM992_04785\|DM992_03020\|DM992_04750\|DM992_03725\|DM992_04725\|DM992_04790<br>DM992_03020\|DM992_04750\|DM992_03725\|DM992_04510\|DM992_04520\|DM992_29375\|DM992_03015\|DM992_04760<br>DM992_04745\|DM992_04745\|DM992_02935\|DM992_02935\|DM992_04765\|DM992_03075\|DM992_04725\|DM992_04790<br>DM992_04510\|DM992_04520\|DM992_29375\|DM992_03015\|DM992_04760 |
| Bacterial chemotaxis | bct02030 | 19 | 36 | 1.71E-10 | 5.46E-09 | DM992_02990\|DM992_03020\|DM992_03005\|DM992_24480\|DM992_02980\|DM992_05700\|DM992_04625\|DM992_29375<br>DM992_04520\|DM992_03725\|DM992_02985\|DM992_02975\|DM992_31425\|DM992_03000\|DM992_18385\|DM992_03010<br>DM992_03015\|DM992_02970\|DM992_24475 |
| Two-component system | bct02020 | 26 | 137 | 4.11E-06 | 8.77E-05 | DM992_39765\|DM992_33890\|DM992_03020\|DM992_03000\|DM992_03005\|DM992_13970\|DM992_24475\|DM992_02990<br>DM992_05790\|DM992_05795\|DM992_03010\|DM992_03030\|DM992_06085\|DM992_03075\|DM992_24480\|DM992_02980<br>DM992_40690\|DM992_13985\|DM992_05700\|DM992_12640\|DM992_29375\|DM992_31425\|DM992_18385\|DM992_33455<br>DM992_12410\|DM992_02975 |
| Carbon metabolism | bct01200 | 14 | 127 | 0.045771221 | 0.562894776 | DM992_20510\|DM992_20505\|DM992_06085\|DM992_06090\|DM992_21865\|DM992_39795<br>DM992_39790\|DM992_10125\|DM992_39830\|DM992_39785\|DM992_12350\|DM992_03180\|DM992_12360\|DM992_20595 |
| Chloroalkane and chloroalkene degradation | bct00625 | 3 | 11 | 0.048891576 | 0.562894776 | DM992_32870\|DM992_12400\|DM992_12350 |
| Fatty acid biosynthesis | bct00061 | 4 | 21 | 0.061332921 | 0.562894776 | DM992_38910\|DM992_40265\|DM992_21850\|DM992_33045 |
| Methane metabolism | bct00680 | 5 | 31 | 0.064043202 | 0.562894776 | DM992_20510\|DM992_20505\|DM992_20595\|DM992_39785\|DM992_12350 |
| Glycolysis / Gluconeogenesis | bct00010 | 5 | 33 | 0.077394647 | 0.562894776 | DM992_32870\|DM992_39795\|DM992_12400\|DM992_39790\|DM992_39785 |
| Pyruvate metabolism | bct00620 | 7 | 56 | 0.085271629 | 0.562894776 | DM992_06085\|DM992_21865\|DM992_39795\|DM992_39790\|DM992_27175\|DM992_39785\|DM992_20595 |
| Glyoxylate and dicarboxylate metabolism | bct00630 | 8 | 68 | 0.087952309 | 0.562894776 | DM992_20510\|DM992_20505\|DM992_06085\|DM992_06090\|DM992_10125\|DM992_03180\|DM992_12360\|DM992_03185 |
| Pentose and glucuronate interconversions | bct00040 | 3 | 18 | 0.131263488 | 0.662833539 | DM992_34385\|DM992_30755\|DM992_29905 |
| Fatty acid metabolism | bct01212 | 5 | 40 | 0.134638063 | 0.662833539 | DM992_38910\|DM992_06085\|DM992_40265\|DM992_21850\|DM992_33045 |
| Amino sugar and nucleotide sugar metabolism | bct00520 | 5 | 40 | 0.134638063 | 0.662833539 | DM992_17170\|DM992_30755\|DM992_30725\|DM992_29905\|DM992_29895 |
| Fatty acid degradation | bct00071 | 4 | 30 | 0.148174712 | 0.677370114 | DM992_32870\|DM992_06085\|DM992_12400\|DM992_40265 |
| Nicotinate and nicotinamide metabolism | bct00760 | 3 | 20 | 0.160295325 | 0.683926718 | DM992_39840\|DM992_20555\|DM992_39825 |
| Galactose metabolism | bct00052 | 2 | 11 | 0.184464944 | 0.737859776 | DM992_17170\|DM992_30755 |
| Alanine, aspartate and glutamate metabolism | bct00250 | 4 | 38 | 0.249116096 | 0.883205108 | DM992_38920\|DM992_33335\|DM992_19985\|DM992_21725 |
| Quorum sensing | bct02024 | 10 | 120 | 0.263278825 | 0.883205108 | DM992_03030\|DM992_40690\|DM992_27160\|DM992_02885\|DM992_33455\|DM992_27165\|DM992_27170\|DM992_33845<br>DM992_12305\|DM992_40265 |
| Butanoate metabolism | bct00650 | 5 | 53 | 0.274558578 | 0.883205108 | DM992_33335\|DM992_12355\|DM992_06085\|DM992_06090\|DM992_12360 |
| RNA degradation | bct03018 | 2 | 15 | 0.276001596 | 0.883205108 | DM992_35840\|DM992_20465 |
| Pentose phosphate pathway | bct00030 | 3 | 28 | 0.289807091 | 0.88322161 | DM992_17795\|DM992_39830\|DM992_35780 |
| Nitrogen metabolism | bct00910 | 3 | 29 | 0.306759148 | 0.892390248 | DM992_02890\|DM992_11935\|DM992_31250 |
| beta-Lactam resistance | bct01501 | 3 | 34 | 0.391411443 | 0.964259232 | DM992_12305\|DM992_13960\|DM992_13970 |
| Starch and sucrose metabolism | bct00500 | 2 | 21 | 0.411062415 | 0.964259232 | DM992_30755\|DM992_29905 |
| Fructose and mannose metabolism | bct00051 | 2 | 22 | 0.432478931 | 0.964259232 | DM992_30725\|DM992_29895 |
| Degradation of aromatic compounds | bct01220 | 4 | 56 | 0.498183912 | 0.964259232 | DM992_32870\|DM992_33030\|DM992_12400\|DM992_34755 |
| Biosynthesis of unsaturated fatty acids | bct01040 | 1 | 11 | 0.528486316 | 0.964259232 | DM992_21850 |
| Sulfur relay system | bct04122 | 1 | 11 | 0.528486316 | 0.964259232 | DM992_04415 |
| Oxidative phosphorylation | bct00190 | 4 | 60 | 0.549852983 | 0.964259232 | DM992_21875\|DM992_05790\|DM992_21870\|DM992_05795 |
| Cationic antimicrobial peptide (CAMP) resistance | bct01503 | 1 | 12 | 0.557154764 | 0.964259232 | DM992_13970 |
| Steroid degradation | bct00984 | 1 | 12 | 0.557154764 | 0.964259232 | DM992_38820 |
| Propanoate metabolism | bct00640 | 3 | 46 | 0.578517418 | 0.964259232 | DM992_33335\|DM992_06085\|DM992_39785 |
| Carbon fixation in photosynthetic organisms | bct00710 | 1 | 13 | 0.584085092 | 0.964259232 | DM992_21865 |
| Tyrosine metabolism | bct00350 | 2 | 30 | 0.586840792 | 0.964259232 | DM992_32870\|DM992_12400 |
| Citrate cycle (TCA cycle) | bct00020 | 2 | 30 | 0.586840792 | 0.964259232 | DM992_39795\|DM992_39790 |
| Synthesis and degradation of ketone bodies | bct00072 | 1 | 14 | 0.609382378 | 0.964259232 | DM992_06085 |
| Arginine and proline metabolism | bct00330 | 3 | 50 | 0.632396729 | 0.964259232 | DM992_04885\|DM992_20325\|DM992_20335 |
| Biotin metabolism | bct00780 | 1 | 16 | 0.655466855 | 0.964259232 | DM992_21850 |
| Selenocompound metabolism | bct00450 | 1 | 16 | 0.655466855 | 0.964259232 | DM992_24060 |
| One carbon pool by folate | bct00670 | 1 | 17 | 0.676434031 | 0.964259232 | DM992_03180 |
| Benzoate degradation | bct00362 | 3 | 55 | 0.692773142 | 0.964259232 | DM992_06085\|DM992_33030\|DM992_34755 |
| Cysteine and methionine metabolism | bct00270 | 2 | 37 | 0.694615486 | 0.964259232 | DM992_24060\|DM992_02915 |
| Microbial metabolism in diverse environments | bct01120 | 20 | 341 | 0.700898341 | 0.964259232 | DM992_20510\|DM992_32870\|DM992_34755\|DM992_33030\|DM992_12410\|DM992_06090\|DM992_21865\|DM992_12360<br>DM992_39795\|DM992_39790\|DM992_12400\|DM992_10125\|DM992_33335\|DM992_06085\|DM992_20505\|DM992_39830<br>DM992_38820\|DM992_20595\|DM992_39785\|DM992_12350 |
| Folate biosynthesis | bct00790 | 1 | 19 | 0.714628255 | 0.964259232 | DM992_04415 |
| Terpenoid backbone biosynthesis | bct00900 | 1 | 19 | 0.714628255 | 0.964259232 | DM992_06085 |
| Valine, leucine and isoleucine biosynthesis | bct00290 | 1 | 21 | 0.74832598 | 0.964259232 | DM992_27175 |
| beta-Alanine metabolism | bct00410 | 1 | 21 | 0.74832598 | 0.964259232 | DM992_33335 |
| Porphyrin and chlorophyll metabolism | bct00860 | 2 | 42 | 0.7565512 | 0.964259232 | DM992_17040\|DM992_33360 |
| Ascorbate and aldarate metabolism | bct00053 | 1 | 22 | 0.763656142 | 0.964259232 | DM992_29905 |
| Metabolic pathways | bct01100 | 53 | 893 | 0.766075921 | 0.964259232 | DM992_21725\|DM992_03180\|DM992_34755\|DM992_17170\|DM992_40265\|DM992_33045\|DM992_39840\|DM992_19985<br>DM992_20510\|DM992_32870\|DM992_04415\|DM992_05790\|DM992_34385\|DM992_20325\|DM992_24060\|DM992_04885<br>DM992_21870\|DM992_03185\|DM992_20595\|DM992_33030\|DM992_21850\|DM992_39795\|DM992_35775\|DM992_39790<br>DM992_38820\|DM992_33335\|DM992_12400\|DM992_39830\|DM992_12350\|DM992_20555\|DM992_30725\|DM992_20505<br>DM992_30755\|DM992_12410\|DM992_27175\|DM992_33360\|DM992_39825\|DM992_39785\|DM992_17795\|DM992_17790<br>DM992_21865\|DM992_38910\|DM992_02915\|DM992_29895\|DM992_17040\|DM992_21875\|DM992_29905\|DM992_05795<br>DM992_06085\|DM992_35780\|DM992_38920\|DM992_20335\|DM992_10125 |
| Bacterial secretion system | bct03070 | 2 | 45 | 0.788274095 | 0.964259232 | DM992_13970\|DM992_31485 |
| Lysine degradation | bct00310 | 1 | 25 | 0.804281894 | 0.964259232 | DM992_06085 |
| Pantothenate and CoA biosynthesis | bct00770 | 1 | 26 | 0.816212408 | 0.964259232 | DM992_33065 |
| Pyrimidine metabolism | bct00240 | 2 | 48 | 0.816306954 | 0.964259232 | DM992_35775\|DM992_17790 |
| Glycine, serine and threonine metabolism | bct00260 | 2 | 52 | 0.848513321 | 0.964259232 | DM992_03180\|DM992_03185 |
| Tryptophan metabolism | bct00380 | 1 | 31 | 0.865827298 | 0.964259232 | DM992_06085 |
| Biosynthesis of secondary metabolites | bct01110 | 17 | 334 | 0.868028958 | 0.964259232 | DM992_32870\|DM992_06085\|DM992_38920\|DM992_39795\|DM992_03180\|DM992_39790\|DM992_33360\|DM992_03185<br>DM992_27175\|DM992_10125\|DM992_24060\|DM992_21725\|DM992_12400\|DM992_39830\|DM992_39785\|DM992_17040<br>DM992_29895 |
| Purine metabolism | bct00230 | 3 | 80 | 0.887846092 | 0.964259232 | DM992_35775\|DM992_17790\|DM992_39830 |
| 2-Oxocarboxylic acid metabolism | bct01210 | 1 | 34 | 0.88892648 | 0.964259232 | DM992_27175 |
| Valine, leucine and isoleucine degradation | bct00280 | 1 | 43 | 0.937025003 | 0.988252759 | DM992_06085 |
| Sulfur metabolism | bct00920 | 1 | 49 | 0.956884033 | 0.988252759 | DM992_12410 |
| Biosynthesis of antibiotics | bct01130 | 10 | 248 | 0.95736986 | 0.988252759 | DM992_32870\|DM992_06085\|DM992_39795\|DM992_30755\|DM992_39790\|DM992_12400\|DM992_39830\|DM992_39785<br>DM992_03180\|DM992_03185 |
| ABC transporters | bct02010 | 7 | 236 | 0.993402699 | 0.995107464 | DM992_33395\|DM992_13985\|DM992_02885\|DM992_04625\|DM992_04880\|DM992_04870\|DM992_04875 |
| Biosynthesis of amino acids | bct01230 | 3 | 145 | 0.995107464 | 0.995107464 | DM992_24060\|DM992_39830\|DM992_27175 |

**Supplementary Table S5.**
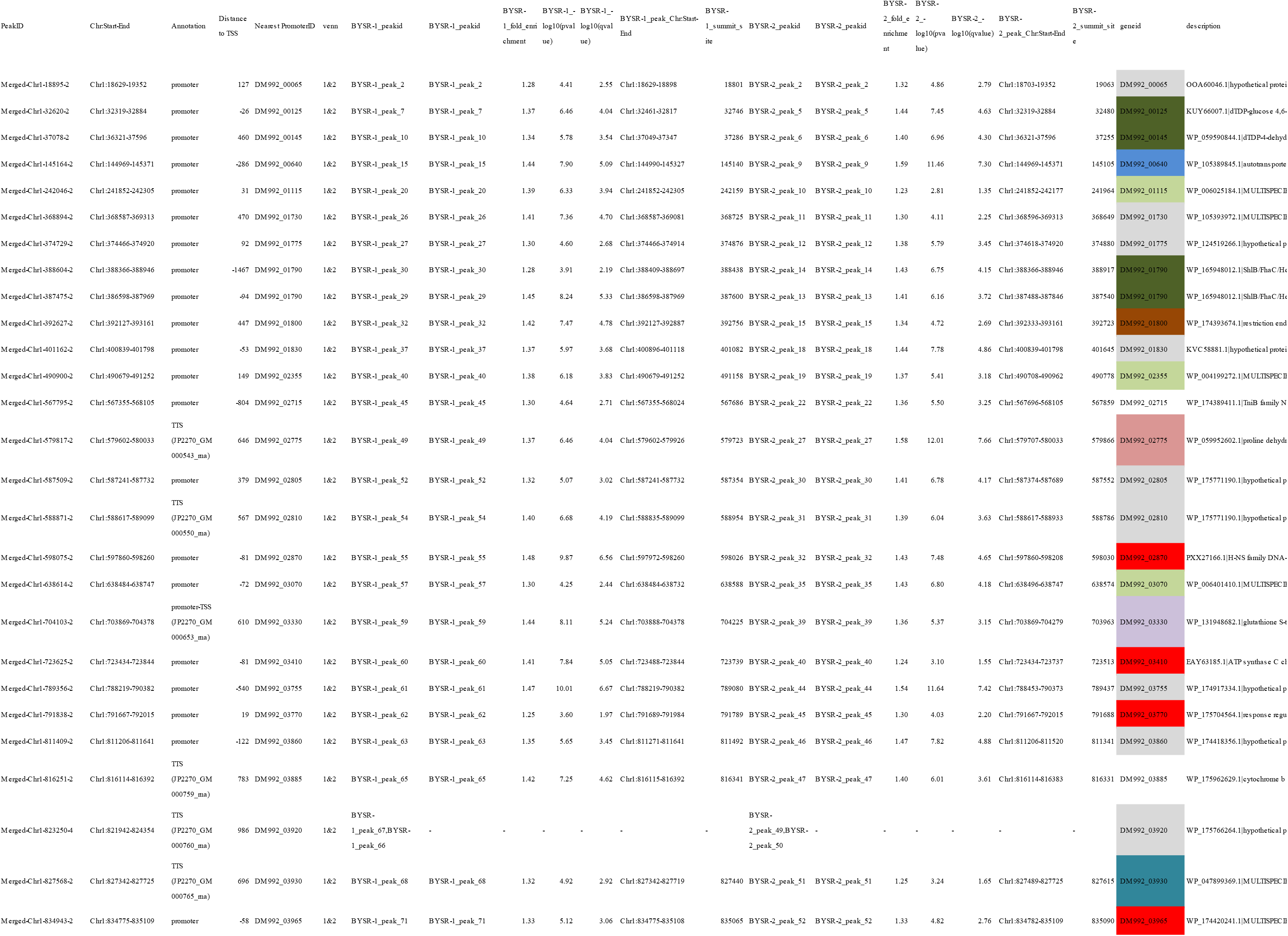
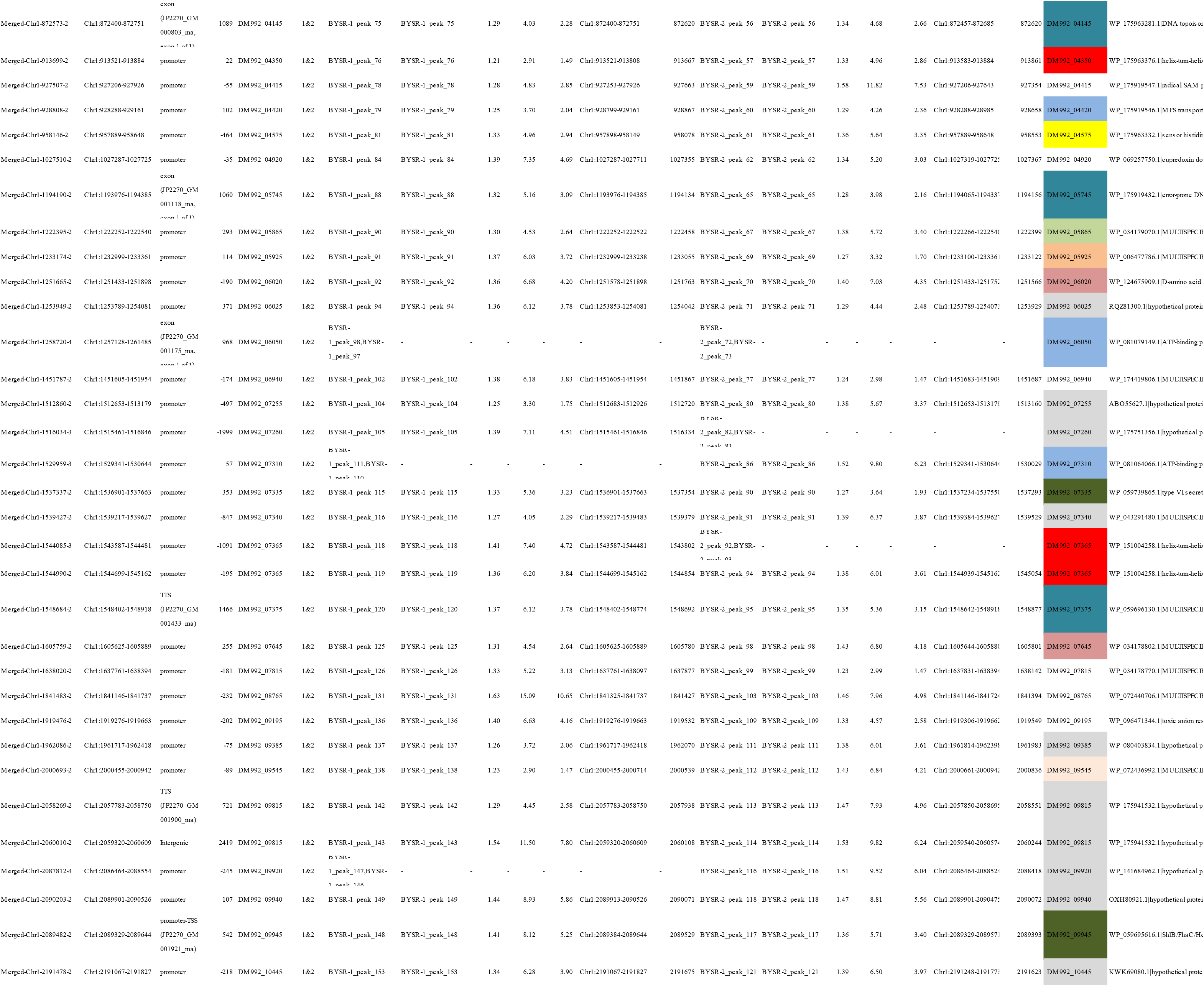
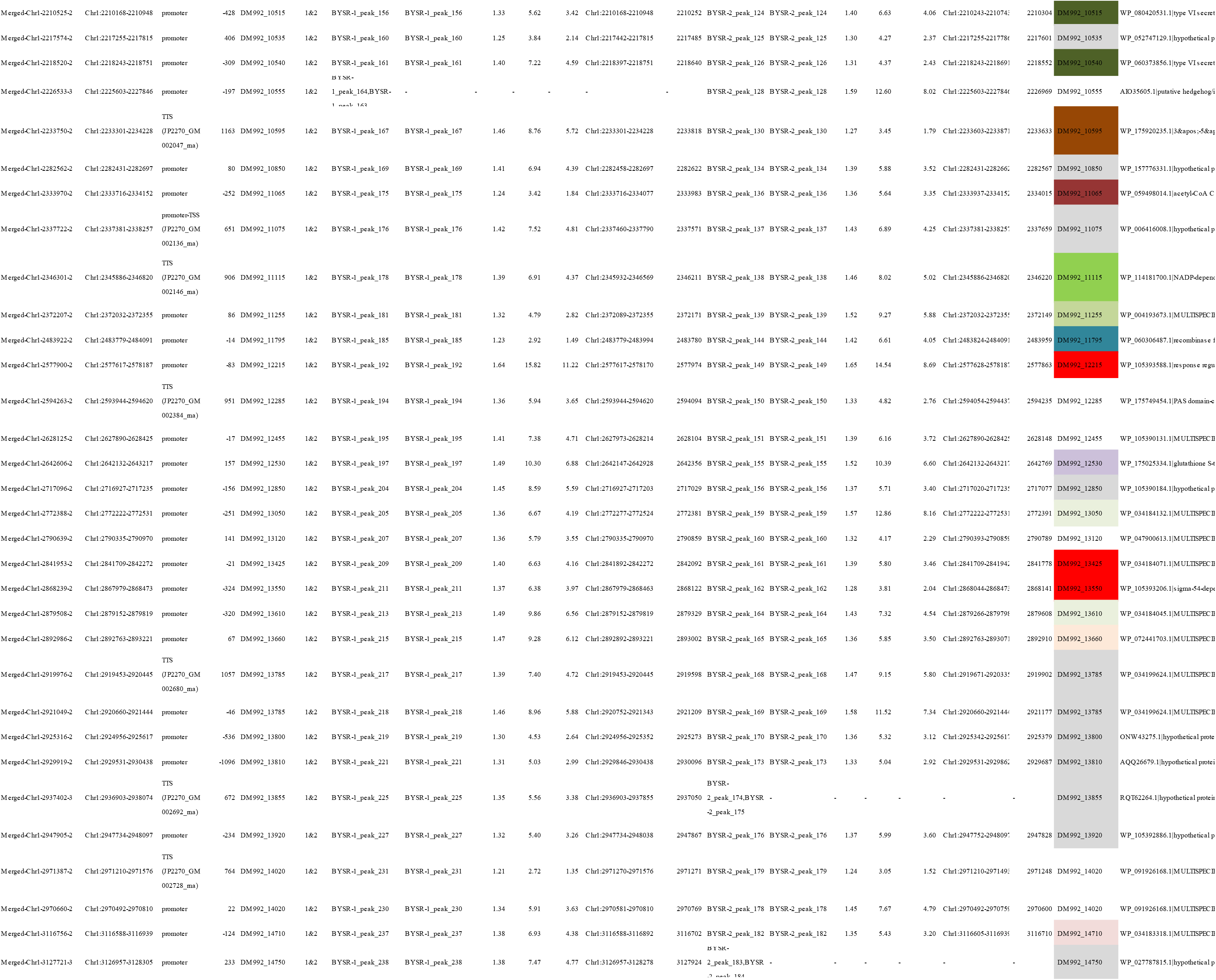
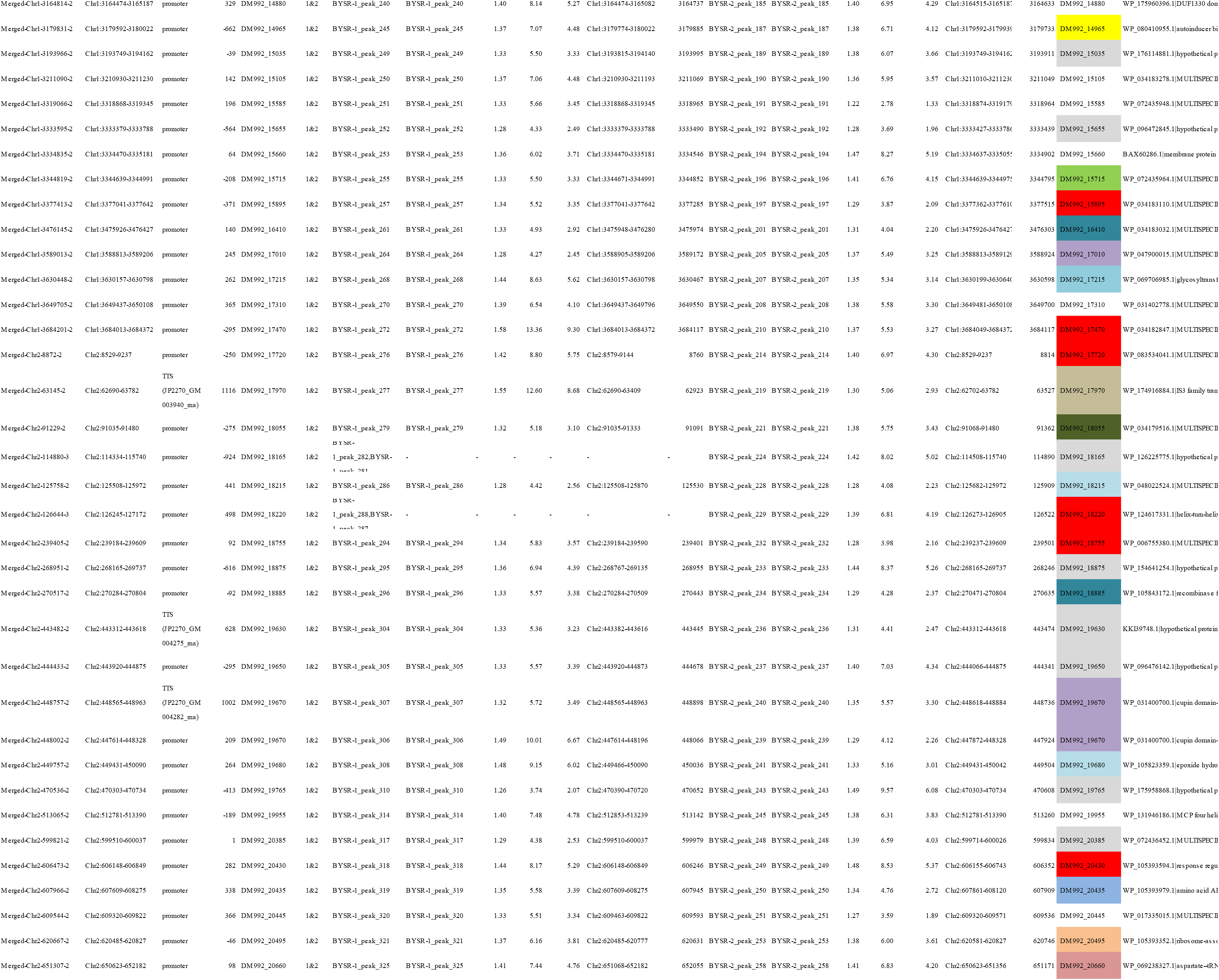
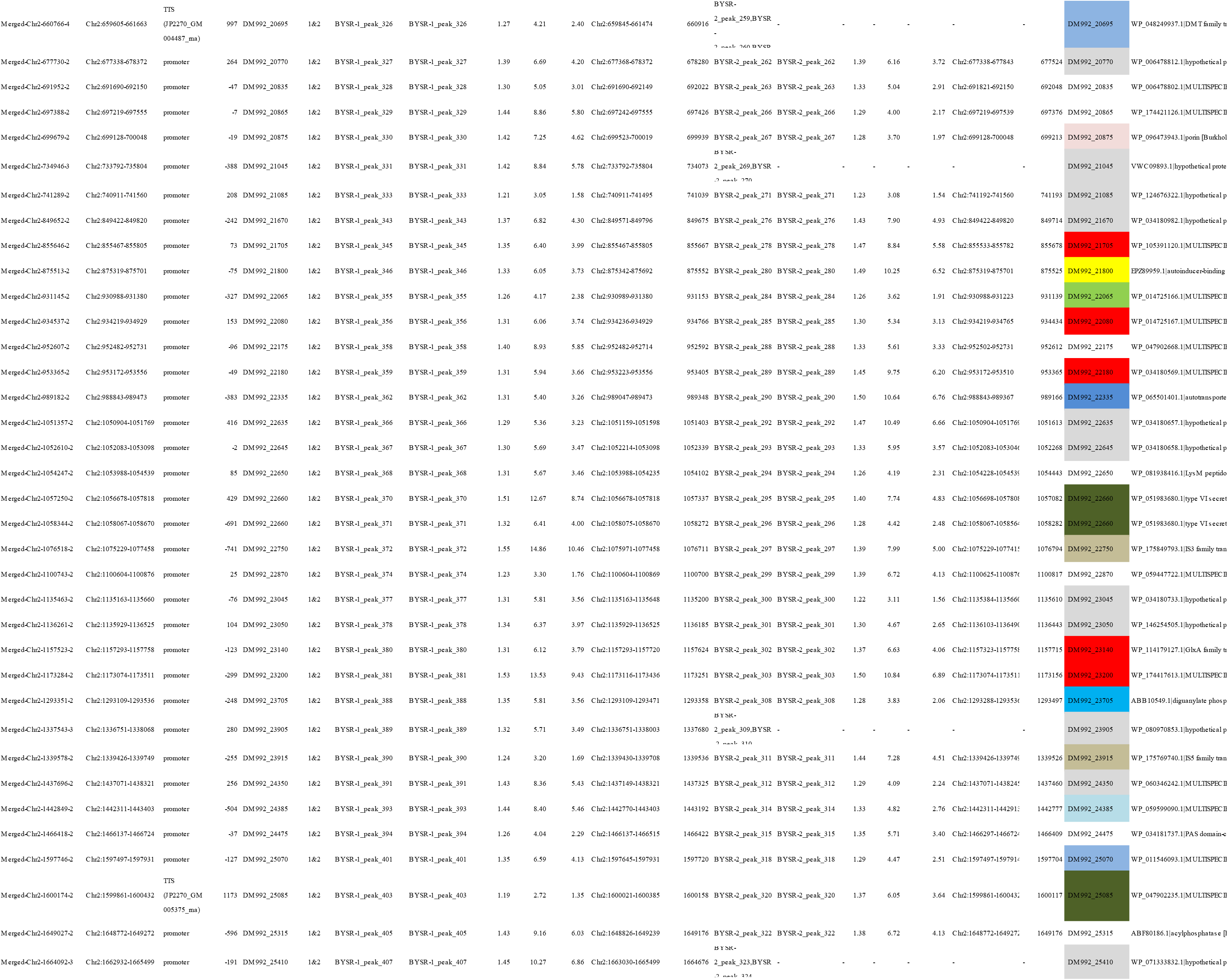
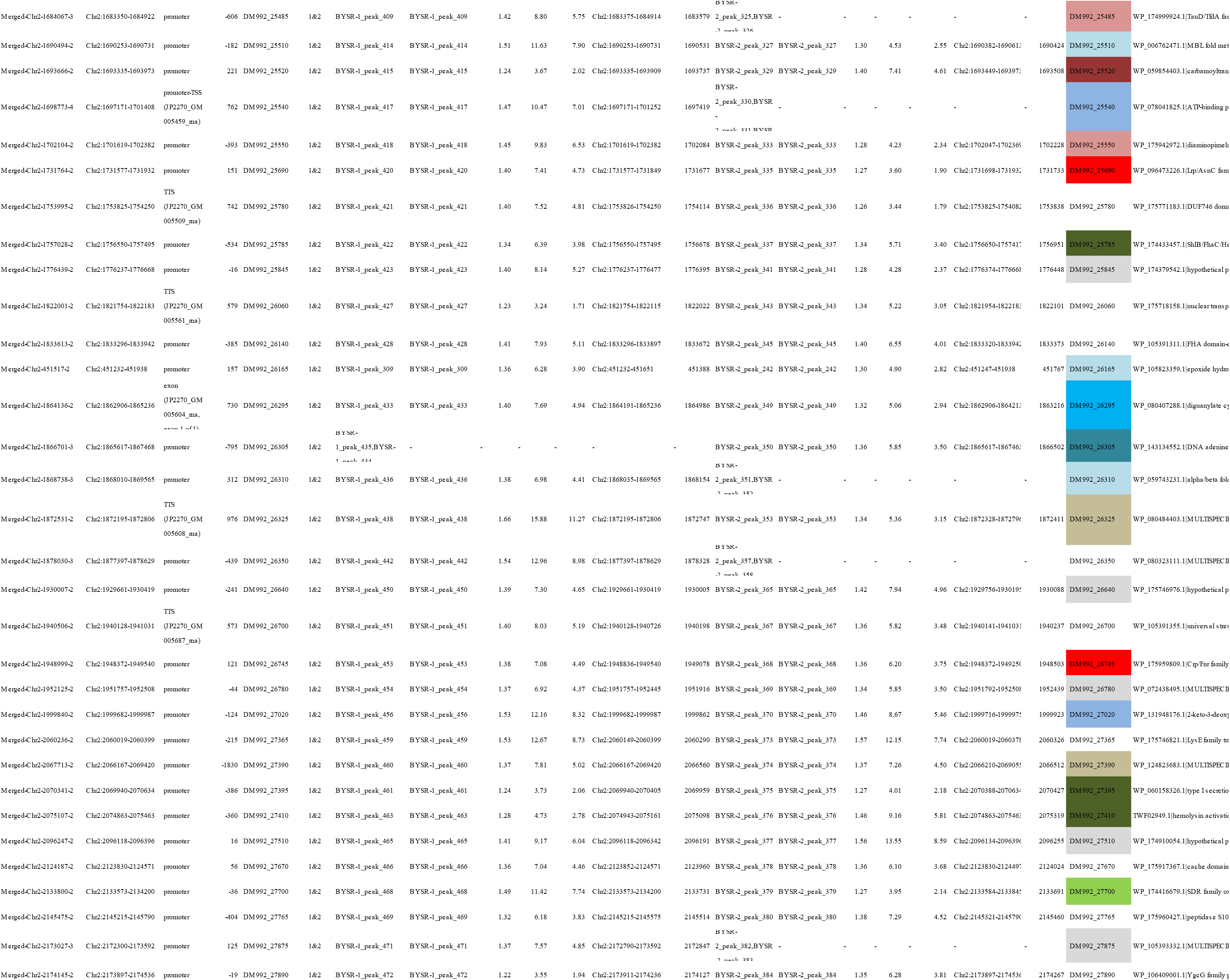
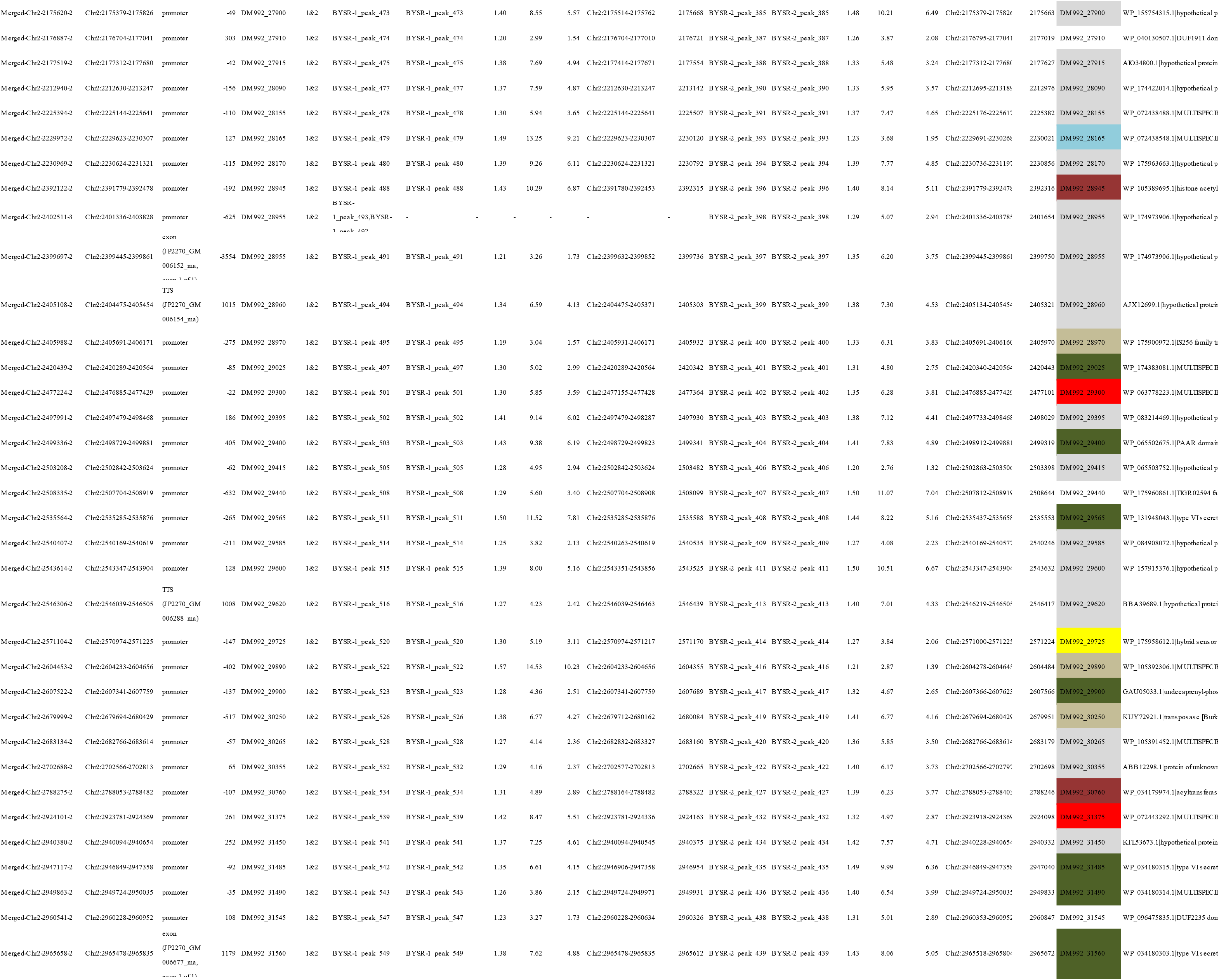
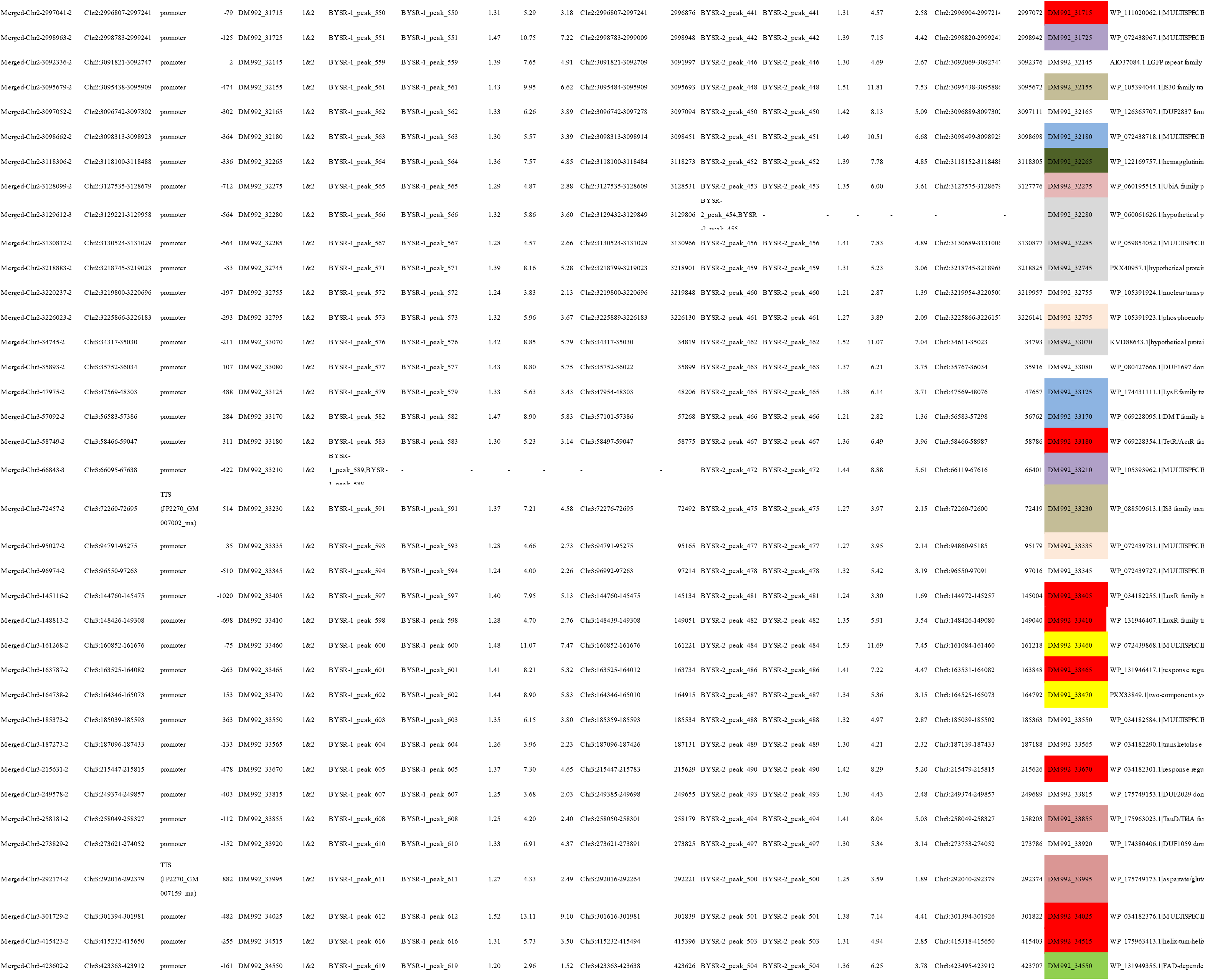
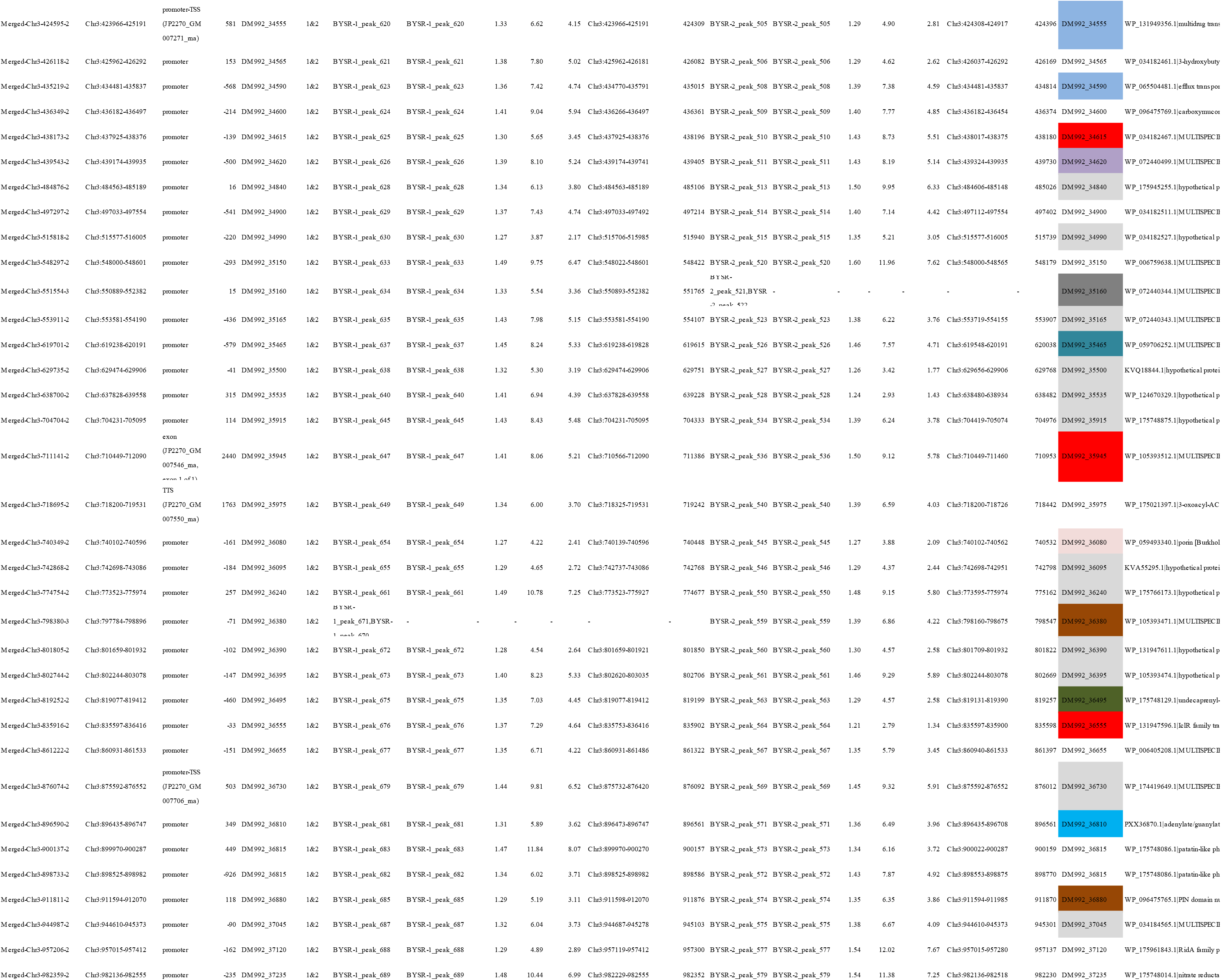
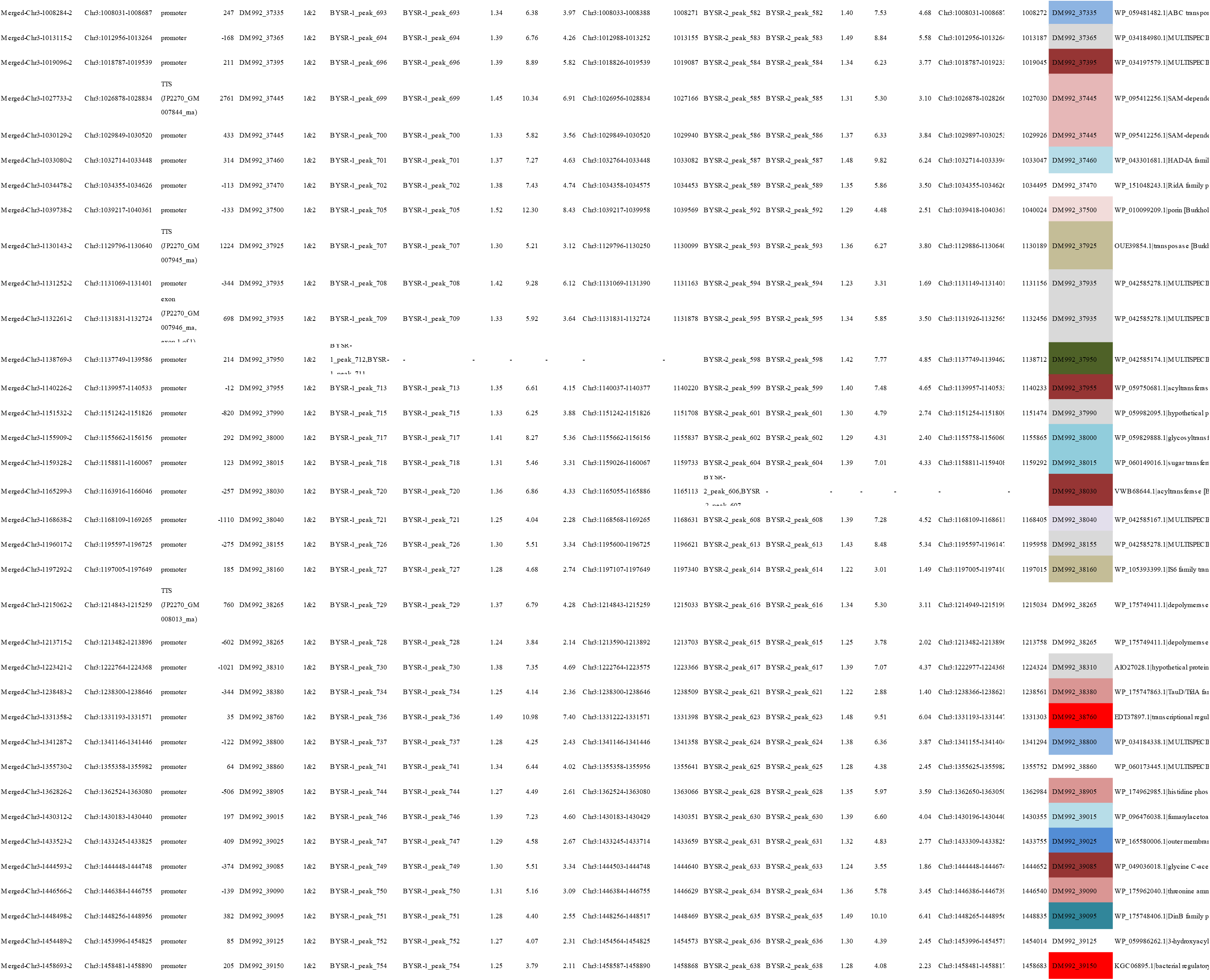
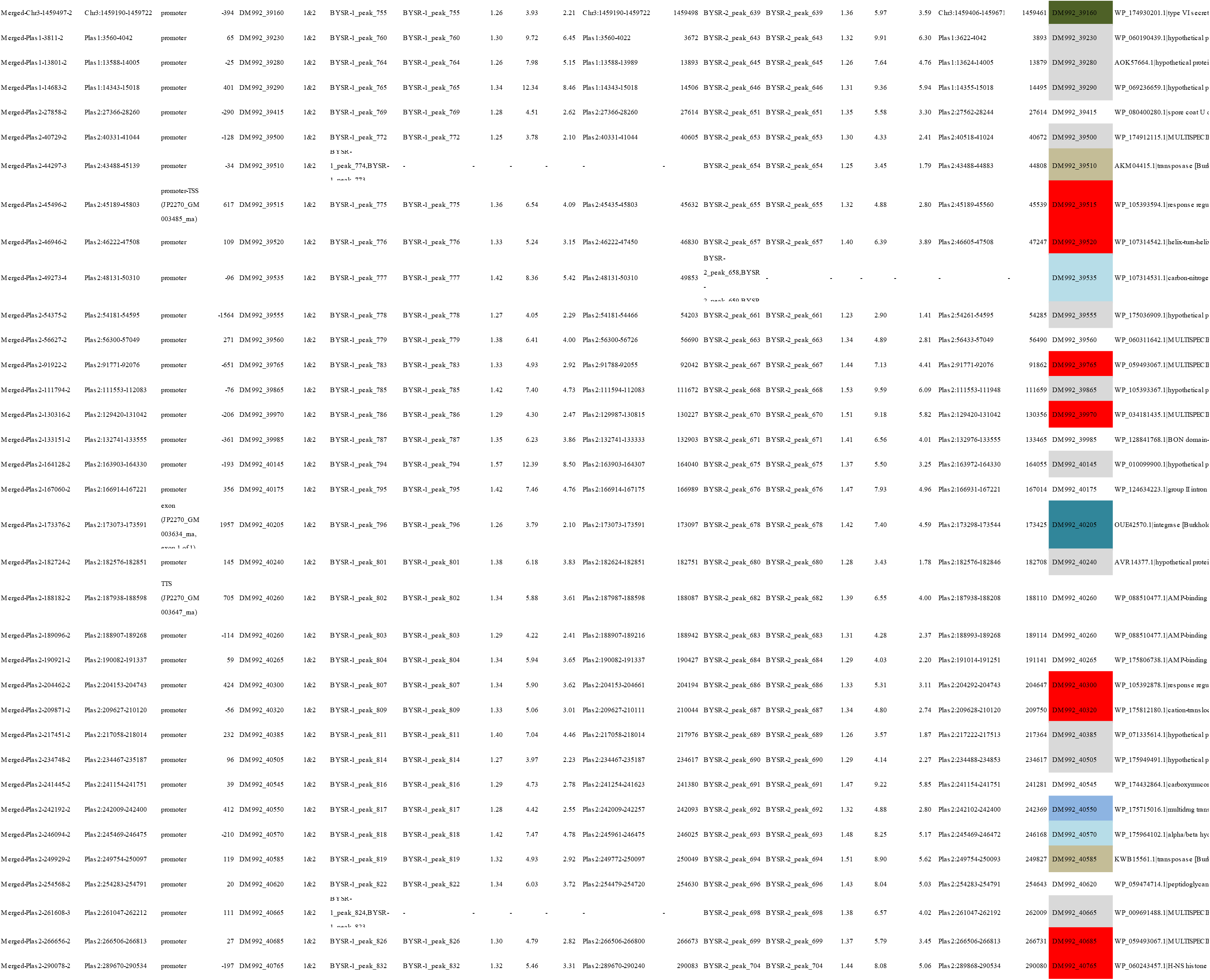
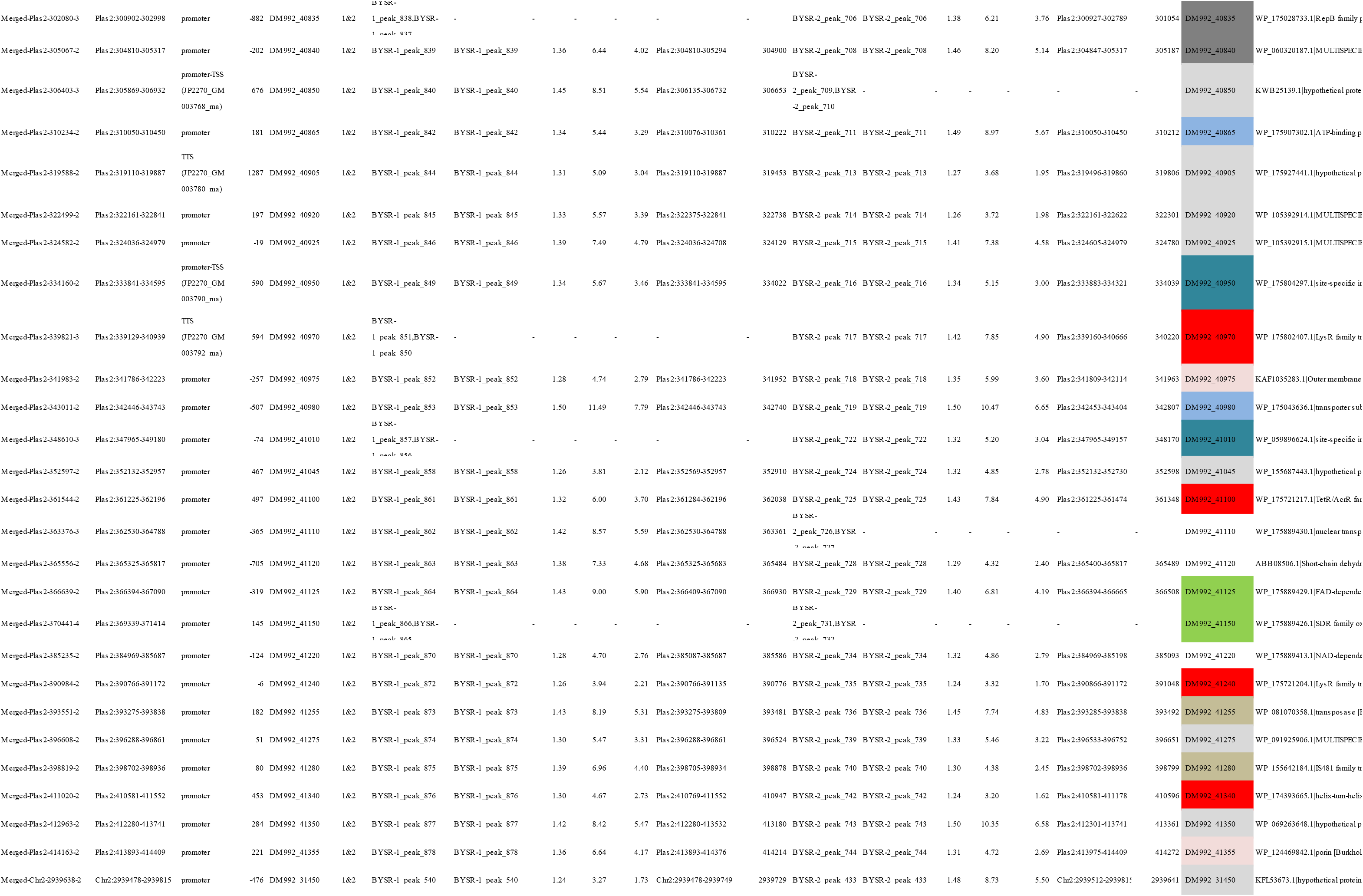
The enriched peaks identified by DAP-seq

## REFERENCES

1. Elshafie HS, Camele I. 2021. An Overview of Metabolic Activity, Beneficial and Pathogenic Aspects of Burkholderia Spp. Metabolites 11.

2. Jia J, Ford E, Hobbs S, Baird S, Lu S. 2021. Occidiofungin is the key metabolite for antifungal activity of the endophytic bacterium Burkholderia sp. MS455 against Aspergillus flavus. Phytopathology doi:10.1094/PHYTO-06-21-0225-R.

3. Hendry S, Steinke S, Wittstein K, Stadler M, Harmrolfs K, Adewunmi Y, Sahukhal G, Elasri M, Thomashow L, Weller D, Mavrodi O, Blankenfeldt W, Mavrodi D. 2021. Functional Analysis of Phenazine Biosynthesis Genes in Burkholderia spp. Appl Environ Microbiol 87.

4. Imade EE, Babalola OO. 2021. Biotechnological utilization: the role of Zea mays rhizospheric bacteria in ecosystem sustainability. Appl Microbiol Biotechnol 105:4487–4500.

5. Parke JL, Gurian-Sherman D. 2001. Diversity of the Burkholderia cepacia complex and implications for risk assessment of biological control strains. Annu Rev Phytopathol 39:225–58.

6. Mullins AJ, Murray JAH, Bull MJ, Jenner M, Jones C, Webster G, Green AE, Neill DR, Connor TR, Parkhill J, Challis GL, Mahenthiralingam E. 2019. Genome mining identifies cepacin as a plant-protective metabolite of the biopesticidal bacterium Burkholderia ambifaria. Nat Microbiol 4:996–1005.

7. Foxfire A, Buhrow AR, Orugunty RS, Smith L. 2021. Drug discovery through the isolation of natural products from Burkholderia. Expert Opin Drug Discov 16:807–822.

8. Masschelein J, Jenner M, Challis GL. 2017. Antibiotics from Gram-negative bacteria: a comprehensive overview and selected biosynthetic highlights. Nat Prod Rep 34:712–783.

9. Mahenthiralingam E, Song L, Sass A, White J, Wilmot C, Marchbank A, Boaisha O, Paine J, Knight D, Challis GL. 2011. Enacyloxins are products of an unusual hybrid modular polyketide synthase encoded by a cryptic Burkholderia ambifaria Genomic Island. Chem Biol 18:665–77.

10. Lu SE, Novak J, Austin FW, Gu G, Ellis D, Kirk M, Wilson-Stanford S, Tonelli M, Smith L. 2009. Occidiofungin, a unique antifungal glycopeptide produced by a strain of Burkholderia contaminans. Biochemistry 48:8312–21.

11. Ma J, Guo F, Jin Z, Geng M, Ju M, Ravichandran A, Orugunty R, Smith L, Zhu G, Zhang H. 2020. Novel Antiparasitic Activity of the Antifungal Lead Occidiofungin. Antimicrob Agents Chemother 64.

12. Gu G, Smith L, Liu A, Lu SE. 2011. Genetic and biochemical map for the biosynthesis of occidiofungin, an antifungal produced by Burkholderia contaminans strain MS14. Appl Environ Microbiol 77:6189–98.

13. Chen KC, Ravichandran A, Guerrero A, Deng P, Baird SM, Smith L, Lu SE. 2013. The Burkholderia contaminans MS14 ocfC gene encodes a xylosyltransferase for production of the antifungal occidiofungin. Appl Environ Microbiol 79:2899–905.

14. Deng P, Wang X, Baird SM, Showmaker KC, Smith L, Peterson DG, Lu S. 2016. Comparative genome-wide analysis reveals that Burkholderia contaminans MS14 possesses multiple antimicrobial biosynthesis genes but not major genetic loci required for pathogenesis. Microbiologyopen 5:353–69.

15. Gu G, Wang N, Chaney N, Smith L, Lu SE. 2009. AmbR1 is a key transcriptional regulator for production of antifungal activity of Burkholderia contaminans strain MS14. FEMS Microbiol Lett 297:54–60.

16. Song D, Chen G, Liu S, Khaskheli MA, Wu L. 2019. Complete genome sequence of Burkholderia sp. JP2-270, a rhizosphere isolate of rice with antifungal activity against Rhizoctonia solani. Microb Pathog 127:1-6.

17. Maddocks SE, Oyston PCF. 2008. Structure and function of the LysR-type transcriptional regulator (LTTR) family proteins. Microbiology (Reading) 154:3609–3623.

18. Tang G, Wang Y, Luo L. 2014. Transcriptional regulator LsrB of Sinorhizobium meliloti positively regulates the expression of genes involved in lipopolysaccharide biosynthesis. Appl Environ Microbiol 80:5265–73.

19. Le Guillouzer S, Groleau MC, Mauffrey F, Deziel E. 2020. ScmR, a Global Regulator of Gene Expression, Quorum Sensing, pH Homeostasis, and Virulence in Burkholderia thailandensis. J Bacteriol 202.

20. Mao D, Bushin LB, Moon K, Wu Y, Seyedsayamdost MR. 2017. Discovery of scmR as a global regulator of secondary metabolism and virulence in Burkholderia thailandensis E264. Proc Natl Acad Sci U S A 114:E2920–E2928.

21. Chen L, Wang Y, Miao J, Wang Q, Liu Z, Xie W, Liu X, Feng Z, Cheng S, Chi X, Ge Y. 2021. LysR-type transcriptional regulator FinR is required for phenazine and pyrrolnitrin biosynthesis in biocontrol Pseudomonas chlororaphis strain G05. Appl Microbiol Biotechnol 105:7825–7839.

22. Mao XM, Sun ZH, Liang BR, Wang ZB, Feng WH, Huang FL, Li YQ. 2013. Positive feedback regulation of stgR expression for secondary metabolism in Streptomyces coelicolor. J Bacteriol 195:2072–8.

23. Gomes MC, Tasrini Y, Subramoni S, Agnoli K, Feliciano JR, Eberl L, Sokol P, O’Callaghan D, Vergunst AC. 2018. The afc antifungal activity cluster, which is under tight regulatory control of ShvR, is essential for transition from intracellular persistence of Burkholderia cenocepacia to acute pro-inflammatory infection. Plos Pathogens 14.

24. Kim J, Kim JG, Kang Y, Jang JY, Jog GJ, Lim JY, Kim S, Suga H, Nagamatsu T, Hwang I. 2004. Quorum sensing and the LysR-type transcriptional activator ToxR regulate toxoflavin biosynthesis and transport in Burkholderia glumae. Mol Microbiol 54:921–34.

25. Finn RD, Tate J, Mistry J, Coggill PC, Sammut SJ, Hotz HR, Ceric G, Forslund K, Eddy SR, Sonnhammer EL, Bateman A. 2008. The Pfam protein families database. Nucleic Acids Res 36:D281–8.

26. Bartlett A, O’Malley RC, Huang SC, Galli M, Nery JR, Gallavotti A, Ecker JR. 2017. Mapping genome-wide transcription-factor binding sites using DAP-seq. Nat Protoc 12:1659–1672.

27. Bailey TL, Boden M, Buske FA, Frith M, Grant CE, Clementi L, Ren JY, Li WW, Noble WS. 2009. MEME SUITE: tools for motif discovery and searching. Nucleic Acids Research 37:W202–W208.

28. Modrzejewska M, Kawalek A, Bartosik AA. 2021. The LysR-Type Transcriptional Regulator BsrA (PA2121) Controls Vital Metabolic Pathways in Pseudomonas aeruginosa. mSystems 6:e0001521.

29. SL. H, Ravichandran A, J. E, J. C, Austin F, SE. L, S. P, F. S. 2014. Toxicological evaluation of occidiofungin against mice and human cancer cell lines. Pharmacol Pharm 5:1085–1093.

30. Ellis D, Gosai J, Emrick C, Heintz R, Romans L, Gordon D, Lu SE, Austin F, Smith L. 2012. Occidiofungin’s Chemical Stability and In Vitro Potency against Candida Species. Antimicrobial Agents and Chemotherapy 56:765–769.

31. Tan W, Cooley J, Austin F, Lu SE, Pruett SB, Smith L. 2012. Nonclinical Toxicological Evaluation of Occidiofungin, a Unique Glycolipopeptide Antifungal. International Journal of Toxicology 31:326–336.

32. Gu G, Smith L, Wang N, Wang H, Lu SE. 2009. Biosynthesis of an antifungal oligopeptide in Burkholderia contaminans strain MS14. Biochem Biophys Res Commun 380:328–32.

33. Schell MA. 1993. Molecular biology of the LysR family of transcriptional regulators. Annu Rev Microbiol 47:597–626.

34. Fan X, Zhao Z, Sun T, Rou W, Gui C, Saleem T, Zhao X, Xu X, Zhuo T, Hu X, Zou H. 2020. The LysR-Type Transcriptional Regulator CrgA Negatively Regulates the Flagellar Master Regulator flhDC in Ralstonia solanacearum GMI1000. J Bacteriol 203.

35. Nguyen Le Minh P, Velazquez Ruiz C, Vandermeeren S, Abwoyo P, Bervoets I, Charlier D. 2018. Differential protein-DNA contacts for activation and repression by ArgP, a LysR-type (LTTR) transcriptional regulator in Escherichia coli. Microbiol Res 206:141–158.

36. Pan X, Tang M, You J, Osire T, Sun C, Fu W, Yi G, Yang T, Yang ST, Rao Z. 2022. PsrA is a novel regulator contributes to antibiotic synthesis, bacterial virulence, cell motility and extracellular polysaccharides production in Serratia marcescens. Nucleic Acids Res 50:127–148.

37. Fragel SM, Montada A, Heermann R, Baumann U, Schacherl M, Schnetz K. 2019. Characterization of the pleiotropic LysR-type transcription regulator LeuO of Escherichia coli. Nucleic Acids Res 47:7363–7379.

38. McCarter LL. 2006. Regulation of flagella. Curr Opin Microbiol 9:180–6.

39. Zhou L, Zhu C, Fang X, Liu H, Zhong S, Li Y, Liu J, Song Y, Jian X, Lin Z. 2021. Gene duplication drove the loss of awn in sorghum. Mol Plant 14:1831–1845.

40. Wei H, Xu H, Su C, Wang X, Wang L. 2022. Rice CIRCADIAN CLOCK ASSOCIATED1 transcriptionally regulates ABA signaling to confer multiple abiotic stress tolerance. Plant Physiol doi:10.1093/plphys/kiac196.

41. Cao Y, Zeng H, Ku L, Ren Z, Han Y, Su H, Dou D, Liu H, Dong Y, Zhu F, Li T, Zhao Q, Chen Y. 2020. ZmIBH1-1 regulates plant architecture in maize. J Exp Bot 71:2943–2955.

42. Trouillon J, Imbert L, Villard AM, Vernet T, Attree I, Elsen S. 2021. Determination of the two-component systems regulatory network reveals core and accessory regulations across Pseudomonas aeruginosa lineages. Nucleic Acids Res 49:11476–11490.

43. Trouillon J, Ragno M, Simon V, Attree I, Elsen S. 2021. Transcription Inhibitors with XRE DNA-Binding and Cupin Signal-Sensing Domains Drive Metabolic Diversification in Pseudomonas. mSystems 6.

44. Dillon SC, Espinosa E, Hokamp K, Ussery DW, Casadesus J, Dorman CJ. 2012. LeuO is a global regulator of gene expression in Salmonella enterica serovar Typhimurium. Molecular Microbiology 85:1072–1089.

45. Ishihama A, Shimada T, Yamazaki Y. 2016. Transcription profile of Escherichia coli: genomic SELEX search for regulatory targets of transcription factors. Nucleic Acids Res 44:2058–74.

46. Vandenende CS, Vlasschaert M, Seah SY. 2004. Functional characterization of an aminotransferase required for pyoverdine siderophore biosynthesis in Pseudomonas aeruginosa PAO1. J Bacteriol 186:5596–602.

47. Subramoni S, Nguyen DT, Sokol PA. 2011. Burkholderia cenocepacia ShvR-regulated genes that influence colony morphology, biofilm formation, and virulence. Infect Immun 79:2984–97.

48. Lu J, Huang X, Li K, Li S, Zhang M, Wang Y, Jiang H, Xu Y. 2009. LysR family transcriptional regulator PqsR as repressor of pyoluteorin biosynthesis and activator of phenazine-1-carboxylic acid biosynthesis in Pseudomonas sp. M18. J Biotechnol 143:1-9.

49. Shubeita HE, Sambrook JF, McCormick AM. 1987. Molecular cloning and analysis of functional cDNA and genomic clones encoding bovine cellular retinoic acid-binding protein. Proc Natl Acad Sci U S A 84:5645–9.

50. Kvitko BH, Collmer A. 2011. Construction of Pseudomonas syringae pv. tomato DC3000 mutant and polymutant strains. Methods Mol Biol 712:109–28.

51. Langmead B, Salzberg SL. 2012. Fast gapped-read alignment with Bowtie 2. Nat Methods 9:357–9.

52. Kim D, Langmead B, Salzberg SL. 2015. HISAT: a fast spliced aligner with low memory requirements. Nat Methods 12:357–60.

53. Trapnell C, Pachter L, Salzberg SL. 2009. TopHat: discovering splice junctions with RNA-Seq. Bioinformatics 25:1105–11.

54. Mao XZ, Cai T, Olyarchuk JG, Wei LP. 2005. Automated genome annotation and pathway identification using the KEGG Orthology (KO) as a controlled vocabulary. Bioinformatics 21:3787–3793.

55. Houtgast EJ, Sima VM, Bertels K, Al-Ars Z. 2018. Hardware acceleration of BWA-MEM genomic short read mapping for longer read lengths. Comput Biol Chem 75:54–64.

56. Zhang Y, Liu T, Meyer CA, Eeckhoute J, Johnson DS, Bernstein BE, Nusbaum C, Myers RM, Brown M, Li W, Liu XS. 2008. Model-based analysis of ChIP-Seq (MACS). Genome Biol 9:R137.

57. Ma W, Noble WS, Bailey TL. 2014. Motif-based analysis of large nucleotide data sets using MEME-ChIP. Nat Protoc 9:1428–50.

58. Heinz S, Benner C, Spann N, Bertolino E, Lin YC, Laslo P, Cheng JX, Murre C, Singh H, Glass CK. 2010. Simple combinations of lineage-determining transcription factors prime cis-regulatory elements required for macrophage and B cell identities. Mol Cell 38:576–89.

